# Correlative Imaging for Comprehensive Molecular Mapping of Individual Cell Types in Biological Tissues

**DOI:** 10.1101/2024.09.04.611280

**Authors:** Manxi Yang, Mushfeqa Iqfath, Frederick Nguele Meke, Zihan Qu, Emerson L. Hernly, Pei Su, Zhong-Yin Zhang, Julia Laskin

**Affiliations:** Department of Chemistry, Purdue University, West Lafayette, IN, USA, 47907; Borch Department of Medicinal Chemistry and Molecular Pharmacology, Purdue University, West Lafayette, IN, USA, 47907; Department of Chemistry, Northwestern University, Evanston, IL, 60208

## Abstract

Mass spectrometry imaging (MSI) is a powerful technique for label-free spatial mapping of multiple classes of biomolecules in tissue sections. However, differences in desorption and ionization efficiency of different classes of molecules make it challenging to simultaneously map biomolecules at each omics layer in the same tissue sample. Herein, we present a correlative imaging method using nanospray desorption electrospray ionization (nano-DESI) MSI, which enables the spatial mapping of lipids, metabolites, peptides, and proteins with cellular-level spatial resolution in a single tissue section. We demonstrate the molecular profiling of specific cell types and identify truncated peptides in mouse pancreatic tissue. Distinct chemical gradients of peptides and lipids extending from endocrine cells to exocrine cells indicate their different roles in endocrine-exocrine crosstalk and intracellular signaling. The results underscore the power of the developed imaging approach for spatial multi-omics analysis that provides deep insights into cellular diversity and the intricate molecular interactions that occur within heterogenous biological tissues.

## Introduction

Spatial multi-omics enables a comprehensive characterization of gene expression, protein levels, and metabolite profiles within tissue microenvironment, which is critical to mapping genotype to phenotype.^1^ A detailed molecular description of an organism provides insights into the biological processes at the genome level and establishes molecular phenotypes at the proteome and metabolome level.^2^ From genomics and transcriptomics to proteomics and metabolomics, the genetic and phenotypic variation between organisms drastically expands due to differences in cellular states and environmental factors.^3^ Proteome and metabolome respond dynamically to different conditions and processes that modulate the functional state of cells, and are therefore the most sensitive descriptors of the phenotype of a biological system. Spatial multi-omics of tissues with high spatial resolution advances our understanding of cellular activity, communication, and phenotype variations of the same cell type due tissue microenvironment. These insights are important to understanding the development, aging, and disease etiology of biological systems.^3–6^

Despite the burgeoning next-generation sequencing and antibody-based tools for multi-omics spatial profiling, the grand challenge is to enable the untargeted analysis and integration of spatial proteomics, metabolomics, and lipidomics of the same sample.^7^ Mass spectrometry imaging (MSI) is a label-free approach that enables sensitive, highly multiplexed spatial mapping of different classes of biomolecules in tissue sections offering unique capabilities for spatial metabolomics, lipidomics, and proteomics experiments.^8,9^ Matrix-assisted laser desorption/ionization (MALDI) and desorption electrospray ionization (DESI) MSI are established techniques for imaging of both small molecules and proteins in biological samples.^10^ MALDI imaging of spatial metabolomics has been successfully coupled to transcriptomics.^11^ Recent developments have demonstrated the power of correlative imaging of lipids and proteins using MADLI-MSI for understanding the mechanisms and molecular pathology of diseases.^12–15^ Furthermore, several studies have combined different MSI modalities with different microscopy techniques to increase the spatial resolution, expand the depth of molecular information, and place the molecular maps obtained using MSI into the context of tissue anatomy.^14,16–22^ Despite these impressive advancements, the trade-off between the spatial resolution and molecular coverage and challenges associated with the identification of molecules observed in MSI experiments limit the applications of these techniques.

In this study, we present a platform for correlative and multimodal molecular imaging of biological tissues using nanospray desorption electrospray ionization (nano-DESI) MSI and immunofluorescence microscopy (IF). Nano-DESI is an ambient ionization technique, in which molecules are extracted from tissue sections into a localized liquid bridge formed using a specially-designed probe.^23^ Images are generated by scanning the sample under the probe and acquiring mass spectra from different locations on the sample. Nano-DESI does not require sample pretreatment and generates ions using soft, electrospray-like ionization. The ability to tailor the solvent composition to improve the extraction and ionization efficiency of different classes of molecules enhances the sensitivity and expands molecular coverage of lipids, metabolites, small molecules, and glycans.^24–26^ Furthermore, nano-DESI generates multiply charged ions of intact proteoforms extracted from tissues, which enables their imaging and identification using top-down proteomics approaches.^27–29^ These capabilities have been used for molecular imaging of biological tissues with ∼10 µm spatial resolution.^24,30,31^ Because of the strong suppression of protein signals by lipids, delipidation of the tissue sections is performed prior to imaging of proteins. This presents a challenge to obtaining spatial distributions of both lipids/metabolites and proteins in the same tissue sample. In this study, we integrated IF imaging and two different nano-DESI MSI workflows for imaging of lipids and metabolites^23^ and imaging of proteins and peptides,^27^ which addresses this challenge and enables cell-type-specific mapping of these different classes of biomolecules in the same tissue section. We use this platform to identify molecules involved in endocrine-exocrine communication in mouse pancreatic tissues, which are difficult to characterize using other methods.

The pancreatic islets comprising only ∼2% of the pancreas play a vital role in regulating blood sugar levels and various body functions.^32^ Different cell types in the islets contribute to the delicate balance of hormones involved in glucose metabolism and overall metabolic regulation within the body.^33^ Disruptions in hormone secretion or metabolism can lead to diseases such as diabetes, pancreatic cancer, and pancreatitis.^33^ For example, β-cell dysfunction resulting in insufficient insulin production and hyperglycemia, is a major factor in the development of type 1 diabetes.^34^ Although omics research has provided insights into the cellular activity and molecular mechanisms of pancreatic diseases,^35–39^ bulk analysis falls short in understanding the impact of tissue microenvironment and cellular heterogeneity on the disease progression.

The small size of pancreatic islets (typically 100-200 µm) presents a challenge to their characterization using MSI. Recent developments of high-resolution MSI techniques have provided insights into the localization of lipids and metabolites in both mouse and human pancreatic islets.^40,41^ These studies demonstrated that small differences in lipid composition have a pronounced effect on their localization to either endocrine or exocrine pancreas and provided insights into islet heterogeneity.^40,41^ In one of these studies, IF microscopy performed on serial sections was used to visualize islet locations in tissue sections.^41^

Herein, we demonstrate molecular imaging of lipids, metabolites, peptides, and proteins in the same pancreatic tissue section with high spatial resolution. We use IF microscopy of either the same tissue section or a serial section both as a roadmap for islet localization and as a validation method for protein imaging using nano-DESI MSI.^24^ By alternating between positive and negative ionization modes, we obtain a deep coverage of small molecules in tissue sections. We observe that most lipids are relatively tightly localized to either endocrine or exocrine pancreas. Meanwhile, a majority of pancreatic peptides identified in this study show distinctly different localization. We observed ∼43 pancreatic peptides and visualized their distribution with cellular-level spatial resolution. We have identified 16 truncated C-peptides that exhibit distinct chemical gradients extending from endocrine cells to acinar cells. The presence of these gradients and their dependence on the amino acid sequence of different C-peptides indicate that these peptides play distinct roles in endocrine-exocrine communication. This study opens opportunities for studying biological systems using multimodal MSI-based approaches.

## Results

### Overview of the multimodal imaging workflow

The multimodal imaging workflow is illustrated in Fig. 1 and described in more detail in the Materials and Methods section. Pancreatic tissues are first sectioned to 18 µm slices. Consecutive sections are mounted on serial glass slides and their brightfield optical images are acquired prior to imaging experiments. IF staining of insulin and glucagon is performed on one tissue section to identify the localization of β-cells and α-cells in the islets. IF images are then used to guide the selection of regions of interest for the following nano-DESI MSI analysis. Nano-DESI MSI experiments are performed on a small region of an adjacent tissue section containing ∼2-10 islets to map the spatial distribution of biomolecules. Imaging of lipids, metabolites, and peptides is performed using MeOH/H_2_O (9/1, *v/v*) and by alternating between positive and negative ionization modes in line scan acquisitions. The tissue is subsequently washed using ethanol and chloroform to precipitate proteins and remove lipids, respectively.^31^ Protein imaging is performed using the delipidated tissue section and ACN/H_2_O/HCOOH (80/20/0.07, *v/v/v*) as the extraction solvent. On-tissue MS/MS experiments are performed and compared with predicted *in-silico* generated fragmentation spectra in databases for molecular identification. In addition, several tissue sections were stained and imaged using IF after the first nano-DESI imaging step.

**Fig. 1.**
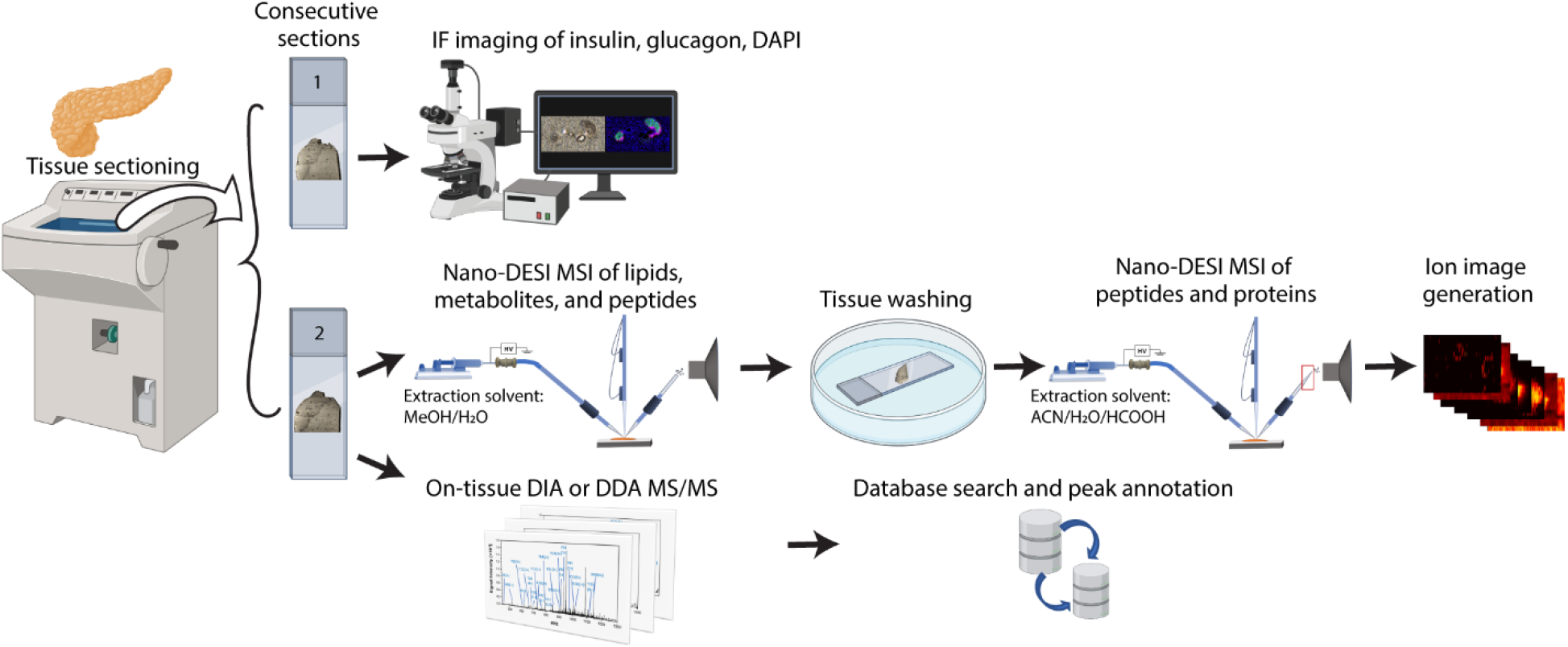
Workflow for multimodal imaging of pancreatic tissue sections. The individual steps are described in detail in the Materials and Methods Section.

### Dual-polarity, corelative nano-DESI MSI of the same pancreatic tissue section maps the spatial distribution of lipids, metabolites, peptides, and proteins in specific cell types

We applied the multimodal imaging workflow to study the distribution of molecules in pancreatic tissue sections of C57BL/6 mice. Images obtained from each imaging modality are shown in Fig. 2. Fig. 2A shows the brightfield optical image of the analyzed region. Fig. 2B shows the overlay of the IF images of insulin (green), glucagon (pink), and cell nuclei (DAPI stain, blue) obtained for the serial tissue section. The IF images of insulin (Fig. 2C) and glucagon (Fig. 2D) indicate the localization of β-cells throughout the islet and α-cells on the periphery of the islets, which is consistent with literature reports.^42^ Selected ion images of lipids and metabolites highlighting representative distinct distribution patterns observed in nano-DESI MSI with MeOH/H_2_O as the extraction solvent are shown in Figs. 2E-I. Meanwhile, representative ion images of proteins and peptides obtained using nano-DESI MSI with ACN/H_2_O/HCOOH as the extraction solvent are shown in Figs. 2J-Q. The spatial distributions of some proteins delineate the localization of specific cell types. As expected, insulin signal observed using nano-DESI MSI (Fig. 2J) is co-localized with β-cells. The excellent correspondence between insulin localization obtained using IF (Fig. 2c) and nano-DESI MSI (Fig. 2J) confirms the accuracy of protein maps obtained using nano-DESI MSI. We used the ion image of insulin in Fig. 2J to determine the outline of the islets, which is shown as a light blue trace in each image in Fig. 2. The ion image of glucagon (Fig. 2K) represents the localization of α-cells, which are observed at the periphery of each islet, which is in good agreement with the results obtained using IF microscopy. We observe a 9490 Da protein (Fig. 2L) co-localizing with insulin, indicating that it is secreted by or/and stored in β-cells.

**Fig. 2.**
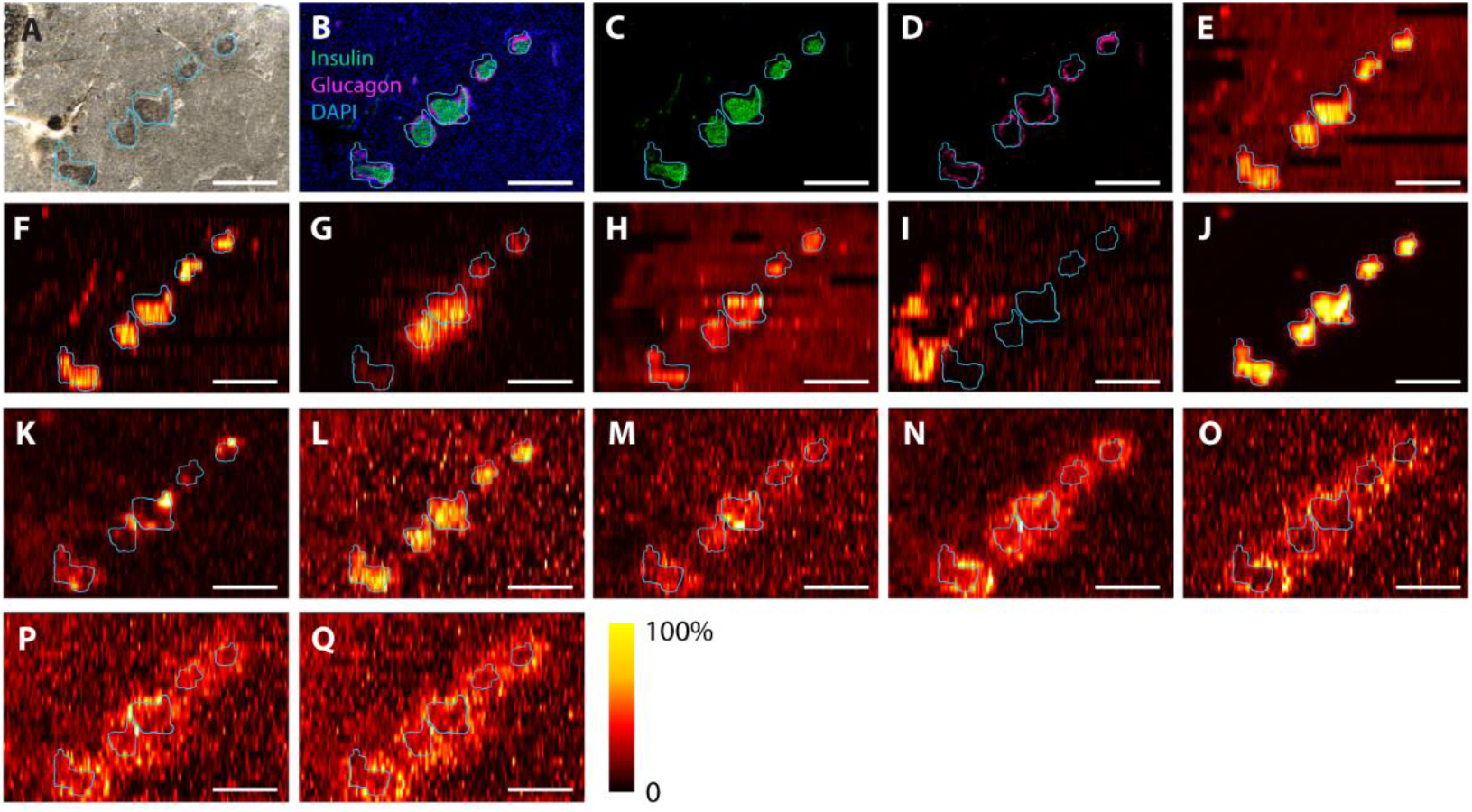
Ion images of lipids, metabolites, peptides and proteins obtained for the same mouse pancreatic tissue section using the multimodal imaging approach. (A) Brightfield optical image of the analyzed region of the pancreatic tissue section. (B) Overlay IF image of insulin (green), glucagon (pink) and DAPI (blue) obtained from the adjacent tissue section. Individual IF image of (C) insulin and (D) glucagon. Ion images of lipids and metabolites normalized to TIC: (E) *m/z* 810.5986^+^, [PC 36:1 + Na]^+^, (F) *m/z* 858.589^+^, [PC 40:5 + Na]^+^, (G) *m/z* 306.0771^-^, [glutathione – H]^-^, (H) *m/z* 810.5311^-^, [PS 38:4 – H]^-^, (I) *m/z* 949.5327^-^. Ion images of intact proteoforms normalized to TIC: (J) *m/z* 1160.241^5+^, 5796 Da, insulin-2, (K) *m/z* 871.583^4+^, 3482 Da, glucagon, (L) *m/z* 1582.659^6+^, 9490 Da, (M) *m/z* 636.343^5+^, 3177 Da, serpinin-RR, (N) *m/z* 1048.775^3+^, 3143 Da, (O) *m/z* 775.947^3+^, 2325 Da, (P) *m/z* 1223.490^2+^, 2445 Da, (Q) *m/z* 1067.422^3+^, 3199 Da. Scale bar: 500 µm. Light blue trace delineates the islets based on the insulin ion image shown in panel (I).

Another interesting species with a unique spatial distribution shown in Fig. 2M is a 3177 Da peptide, which has been identified as serpinin-RR (Fig. S6). Serpinins are a family of peptides that are derived from proteolytic cleavage of the C-terminus of chromogranin-A which is secreted by endocrine cells and various tumor cells.^43,44^ Serpinin peptides have unique biological functions in various tissues and cells. Serpinin-RR was reported recently in mouse brain, pancreas, and ileum tissue extracts.^45^ However, its biological function has not been investigated. The ion image of serpinin-RR in Fig. 2M indicates that it is more abundant in the endocrine cells of the mouse pancreatic tissue, indicating that serpinin-RR may play an active role in the function of pancreatic islets.

Several peptides including the 3143 Da (Fig. 2N), 2325 Da (Fig. 2O), 2445 Da (Fig. 2P), and 3199 Da (Fig. 2Q) species show a distinct halo-like distribution around the pancreatic islets. These peptides are enhanced in the acinar cells in close proximity to β-cells. It has been reported that pancreatic hormones produced by endocrine cells diffuse from the islets through insulo-acinar portal system to acinar cells and preferentially supply the peri-insular region.^46–49^ These islet hormones play important roles in regulating acinar cell functions.^47,50^ While the molecular identification of these peptides will require further study, their unique localization to acini in the peri-insular regions indicates that they may be involved in the endocrine-exocrine intracellular signaling and metabolism pathways.

We have also observed several insulin proteoforms (Fig. S1) including insulin-2 (Fig. S1D) and insulin-1 (Fig. S1E).^51^ The light blue trace overlay shows that all insulin proteoforms have the same spatial distribution and are localized to β-cells. MS/MS spectra of the individual insulin proteoforms obtained using high-energy collisional dissociation (HCD) (Figs. S9-S14) do not provide sufficient structure-specific information to identify their post-translational modifications (PTMs). This is probably due to the presence of the three interchain and intrachain disulfide bonds, which prevent fragmentation along the amino acid sequence in both A and B chain of insulin upon collisional activation. Interestingly, the six insulin proteoforms (Figs. S1F-K) have a +22 Da (Figs. S1F-G), +38 Da (Figs. S1H-I) and +61 Da (Figs. S1J-K) mass shifts as compared to the unmodified insulin-2 (Fig. S1D) and insulin-1 (Fig. S1E), respectively, indicating that insulin-2 and insulin-1 have the same modifications. MS/MS spectra along with annotated fragments and fragmentation maps of all the identified proteins/peptides are shown in Figs. S2-S6. Although several peptides/proteins could not be identified using database searching, we provide their MS/MS spectra in Figs. S7-S14, which can be re-evaluated as databases become more extensive.

Using protein cell type markers, we classify lipid and metabolite localization to specific cell types. For example, phosphatidylcholine (PC) (36:1) (Fig. 2E), PC (40:5) (Fig. 2F), and phosphatidylserine (PS) (38:4) (Fig. 2H) colocalize with insulin signals, indicating that these lipids are either produced or stored in endocrine cells. Interestingly, glutathione (Fig. 2G) shows an enhanced abundance in two islets and lower signal in the other three islets, which could be due to islet heterogeneity. This result underscores the power of corelative nano-DESI MSI in profiling lipid, metabolite and peptide signatures at the cellular level, by co-registering ion images of each molecule with either IF images or ion images of cell markers.

To further validate that proteins do not delocalize during nano-DESI MSI of lipids or in the washing steps, we stained both a tissue section analyzed for lipids and metabolites using nano-DESI MSI and its adjacent fresh tissue section with insulin antibodies. The co-registered insulin IF images acquired for both tissue sections are shown in Fig. S15B. We observe a good correspondence between the IF images obtained for the fresh tissue section and tissue section previously analyzed using nano-DESI MSI. In particular, the insulin fluorescence signal observed for both fresh tissue section and the section analyzed by nano-DESI MSI is tightly localized to the islets outlined by the light blue and yellow traces in both optical and IF images. Furthermore, the overlay image indicates a good overlap between the IF images obtained for both sections with no measurable smearing of the insulin signal in the section analyzed by nano-DESI MSI. This result indicates that there is no noticeable protein delocalization in the section analyzed for lipids and metabolites using nano-DESI MSI. This result validates our correlative imaging workflow and confirms the reliability of protein images obtained using this approach.

### Discovery of truncated forms of pancreatic peptides and visualization of their spatial distributions

An interesting finding reported in this study is that some multiply charged peptides are observed in negative ionization mode nano-DESI MSI of lipids and metabolites performed using MeOH/H_2_O (9/1, *v/v*) as the extraction solvent. Because of the efficient ionization of lipids in nano-DESI and other MSI modalities, peptide signals are rarely observed when analyzing tissue sections that have not been delipidated. Ion images of deprotonated peptides observed in three biological replicates are shown in Figs. 4 and S18-S20. Because negative ionization mode is rarely used by the proteomics community, software that is capable of annotating peptide fragmentation spectra obtained in negative mode is largely non-existent. Meanwhile, manual annotation of peptide MS/MS data is challenging and tedious as it requires one to guess possible peptide sequences and consider numerous isoforms and PTMs. Among all the detected peptides, we were able to manually annotate MS/MS spectra and identify the two intact C-peptide isoforms (Figs. S16D, S16I) and their C-terminal truncated forms several of which have not been previously reported (Figs. S16E-H, S16J-S).

**Fig. 3.**
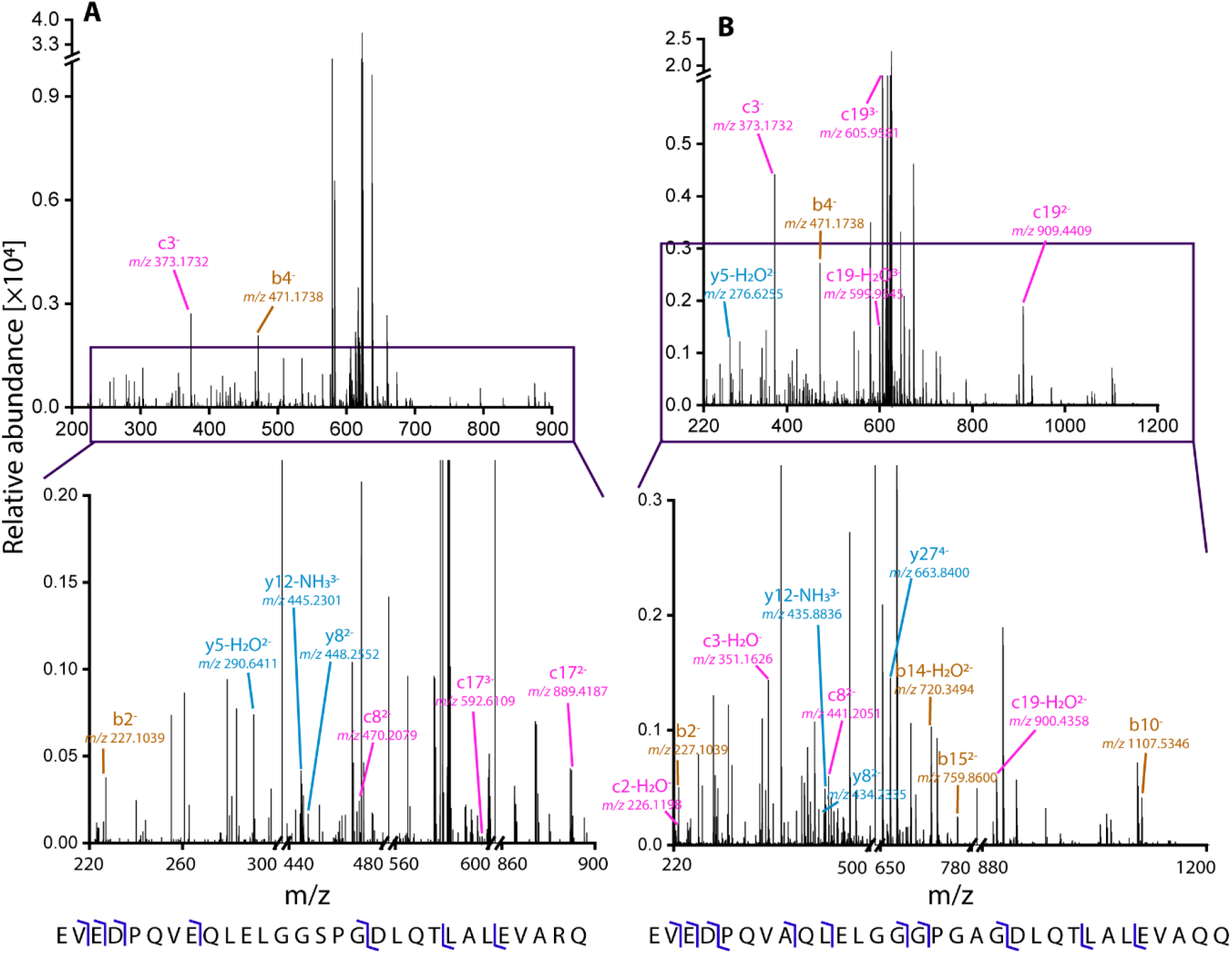
On-tissue identification of the C-peptides by MS/MS. MS/MS spectrum (top), zoomed-in MS/MS spectrum (middle), and the fragmentation map (bottom) of (A) -5 charge state of the 3121 Da C-peptide 1 and (B) -5 charge state of the 3133 Da C-peptide 2.

**Fig. 4.**
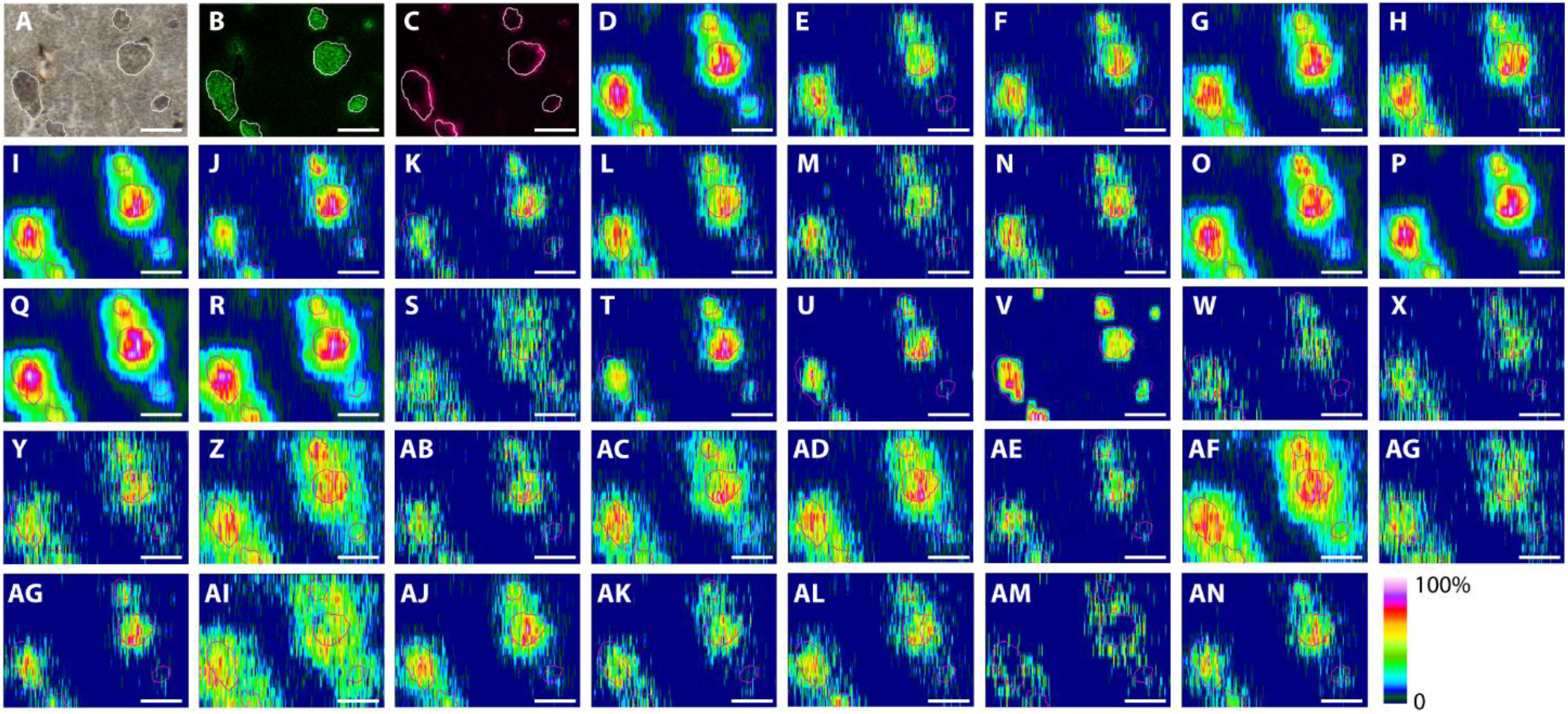
Ion images of pancreatic peptides. (A) Brightfield optical image of the analyzed region of the pancreatic tissue section. IF image of (B) insulin and (C) glucagon on the adjacent section. Ion images of peptides normalized to TIC: (D) *m/z* 623.1063^5-^, 3121 Da, C-peptide 1, (E) *m/z* 559.0068^4-^, 2240 Da, des-(22-29)-C-peptide 1, (F) *m/z* 469.6200^5-^, 2353 Da, des-(23-29)-C-peptide 1, (G) *m/z* 483.8282^5-^, 2424 Da, des-(24-29)-C-peptide 1, (H) *m/z* 633.3064^4-^, 2537 Da, des-(25-29)-C-peptide 1, (I) *m/z* 625.5064^5-^, 3133 Da, C-peptide 2, (J) *m/z* 374.5096^3-^, 1126 Da, des-(11-31)-C-peptide 2, (K) *m/z* 341.1621^4-^, 1369 Da, des-(13-31)-C-peptide 2, (L) *m/z* 606.2909^3-^, 1822 Da, des-(20-31)-C-peptide 2, (M) *m/z* 511.4924^4-^, 2050 Da, des-(22-31)-C-peptide 2, (N) *m/z* 543.7549^4-^, 2179 Da, des-(23-31)-C-peptide 2, (O) *m/z* 569.0170^4-^, 2280 Da, des-(24-31)-C-peptide 2, (P) *m/z* 597.2876^4-^, 2393 Da, des-(25-31)-C-peptide 2, (Q) *m/z* 615.0473^4-^, 2464 Da, des-(26-31)-C-peptide 2, (R) *m/z* 643.3176^4-^, 2577 Da, des-(27-31)-C-peptide 2, (S) *m/z* 599.6972^5-^, 3004 Da, des-(31)-C-peptide 2, (T) *m/z* 640.8114^2-^, 1284 Da, (U) *m/z* 676.3299^2-^, 1355 Da, (V) *m/z* 720.8988^2-^, 1444 Da, (W) *m/z* 436.7023^4-^, 1751 Da, (X) *m/z* 639.6412^3-^, 1922 Da, (Y) *m/z* 526.7459^4-^, 2111 Da, (Z) *m/z* 536.7561^4-^, 2151 Da, (AB) *m/z* 555.0167^4-^, 2224 Da, (AC) *m/z* 565.0266^4-^, 2264 Da, (AD) *m/z* 572.7758^4-^, 2295 Da, (AE) *m/z* 574.5125^4-^, 2302 Da, (AF) *m/z* 582.7870^4-^, 2335 Da, (AG) *m/z* 601.0472^4-^, 2408 Da, (AH) *m/z* 602.7839^4-^, 2415 Da, (AI) *m/z* 611.0578^4-^, 2448 Da, (AJ) *m/z* 620.5431^4-^, 2486 Da, (AK) *m/z* 624.5357^4-^, 2502 Da, (AL) *m/z* 648.8137^4-^, 2599 Da, (AM) *m/z* 750.8808^4-^, 3007 Da, (AN) *m/z* 629.9023^5-^, 3154 Da. Scale bar: 250 µm. White and pink traces delineate the islets based on the IF image shown in panel (B).

C-peptide is part of the proinsulin molecule, which connects insulin A and B chain and promotes the correct folding of proinsulin. When proinsulin is cleaved, it releases C-peptide and bioactive hormone insulin.^52^ C-peptide 1 and C-peptide 2 are the two C-peptide isoforms present in mouse pancreas. The intact forms of the C-peptides are known as biologically active hormones and play important biochemical roles that are different from those of insulin. Specifically, C-peptides interact with various types of cell membrane receptors and participate in intracellular signaling.^52,53^ Meanwhile, little is known about the role of truncated C-peptides. Two C-peptides truncated at the C-terminus, des-(25-29)-C-peptide 1 and des-(27-31)-C-peptide 2 which lack five C-terminal residues, have been observed in the extracts of mouse pancreatic tissue.^54^ A few other truncated forms of C-peptides have been observed in rat or mouse pancreatic islets.^55–57^ However, the biological function of the truncated peptides is not fully understood.^58^ It has been demonstrated that the C-terminal pentapeptide of C-peptide acts as an active site that efficiently stimulates Na^+^/K^+^ ATPase activity of renal tubule 36-80% of the intact C-peptide effect and binds to the same cell membranes as the intact C-peptide.^52,59,60^ It is reasonable to assume that the loss of the C-terminal pentapeptide has a pronounced effect on their biological function. In this study, we observed four truncated peptides derived from the loss of C-terminal residues from C-peptide 1 (Figs. S16E-H) and ten peptides derived from C-peptide 2 (Figs. S16J-S). Using higher energy collisional dissociation (HCD), we mainly detected negatively charged b, c, y ions and their analogs produced by H_2_O and NH_3_ losses in their MS/MS spectra.^61–64^ The annotated MS/MS spectra and the fragmentation maps of the two intact C-peptide isoforms are shown in Fig. 3; the corresponding data obtained for truncated C-peptides are shown in Figs. S20-S33. We note that only confidently assigned terminal fragments and their analogs produced by one H_2_O or NH_3_ loss are annotated in MS/MS spectra. Internal fragments and fragments corresponding to losses of other neutral molecules or multiple neutral losses (CO_2_, CH_2_O, H_2_S, NH=C=NH, C_2_H_4_O, CH_3_, C_2_H_4_, CH_4_S, 2H_2_O, 2NH_3_, etc.) are not labeled in the spectra.

Ion images of all the peptides observed in this study are shown in Fig. 4. The outlines of the islets were determined based on the IF image of insulin (Fig. 4B) and are shown in each image in Fig. 4. As shown in the IF image of glucagon (Fig. 4C), α-cells are localized at the periphery of each islet. A more heterogenous colormap is used in Fig. 4 to better visualize chemical gradients of different peptides in and around the islets. As shown in Figs. 4D-S, both intact C-peptide isoforms and their truncated forms are enhanced in endocrine cells but the gradients extend into acinar cells. Interestingly, we observe differences in chemical gradients of different C-peptides in acinar cells in the peri-insular regions. For example, some peptide signals (Figs. 4E, 4F, 4J, 4K, 4M, and 4N) exhibit a steep decline (steep chemical gradient) at islet boundaries indicating their tight localization to endocrine cells. Meanwhile, several other peptides (Fig. 4D, 4G-I, 4L, 4O-S) show a relatively slow decrease in signal (shallower chemical gradients) that extends farther into the acinar tissue. Line profiles of several C-peptides extracted from ion images are compared with line profiles of insulin and glucagon extracted from IF images in Fig. S19.

Aside from C-peptides and their truncated forms, we have mapped several other peptides that show unique distributions in and near the islets (Figs. 4T-AN). A 2302 Da peptide (Fig. 4AE) matches the intact mass of islet amyloid polypeptide 16-37, while a 2502 Da peptide (Fig. 4AK) matches the intact mass of urocortin-3 16-38. These two peptides have been observed using bulk LC-MS/MS of pancreatic tissues.^45^ They are known to be co-secreted with insulin in β-cells.^65^ Ion images indicate that these two peptides are relatively tightly localized to endocrine cells. Some other unidentified peptides also show a tight localization to endocrine cells. These include a 1284 Da (Fig. 4T), 1355 Da (Fig. 4U), 1444 Da (Fig. 4V), 1751 Da (Fig. 4W), 1922 Da (Fig. 4X), 2224 Da (Fig. 4AB), 2415 Da (Fig. 4AH), and 3154 Da (Fig. 4AN) peptides. Meanwhile, chemical gradients of several other peptides extend into acinar cells to a different extent. The 2151 Da (Fig. 4Z), 2335 Da (Fig. 4AF), and 2448 Da (Fig. 4AI) peptides show the highest extent of signal spreading from endocrine cells to more distant acinar cells as compared to other detected peptides. In the meantime, we observed one 3007 Da peptide (Fig. 4AM) localized only to acinar cells in the peri-insular regions and is absent in endocrine cells. The chemical gradient of these peptides from one cell type to another cell type indicates that they may be involved in the intracellular processes and function as signaling agents. Although we were unable to identify these peptides, their MS/MS spectra are provided in Figs. S34-S53 for reference.

### Chemical gradients of lipids in specific cell types

High-resolution ion images of lipids obtained from three biological replicates using both positive and negative ionization shown in Fig. 5 and Figs. S54-S56 show a much tighter localization than peptides. Representative high-resolution ion images of lipids obtained using both positive and negative ionization modes are shown in Fig. 5 and Figs. S54-S56. Lipids observed in the islets (Fig. 5D) show a much tighter localization to endocrine cells than most of the peptides shown in Fig. 4. Similarly, chemical gradients of lipids abundant in the exocrine tissue (Fig. 5E) do not extend into the interior of the islets. A more comprehensive visualization of lipid maps in mouse pancreatic tissues can be found in our previous publication.^40^ The annotated fragmentation spectra of these lipids can be found in our previous publication.^40^ The outline of β-cells based on the IF image of insulin (Fig. 5B) and that of α-cells based on the IF image of glucagon (Fig. 5C) are overlaid on top of ion images as a white and light blue trace, respectively. Line profiles of several lipids extracted from ion images are compared with line profiles of insulin and glucagon extracted from IF images in Fig. S58. Based on these results, we have identified phospholipids observed in enhanced abundance in endocrine cells. These include LPC (18:0), sphingomyelin (SM) (d34:1), PC (36:2), PC (36:1), PC (38:4), SM (d42:2), and PS (38:4) shown in Fig. 5D. Line profiles provided in Fig. S58 indicate that the signals of PC (36:1), SM (d34:1), and PS (38:4) in the islets are colocalized with insulin. We observe that SM (d34:1) has a steeper chemical gradient at the edge of the islet as compared to PC (36:1) and PS (38:4). We also observe several lipids that show a low signal in the islets and are localized to acinar cells. These include LPC (18:3), LPC (18:2), LPC (20:4), LPC (22:6), fatty acid (FA) (20:5), FA (22:5), lyso-phosphatidic acid (LPA) (18:2), LPA (20:4), lyso-phosphatidylethanolamine (LPE) (18:2), LPE (20:4), LPE (20:3) and LPI (20:4) as shown in Fig. 5E. Line profiles of some of these species shown in Fig. S58 indicate a steep decline of LPC (18:2) and LPE (18:2) signals at the edge of the islets and relatively shallow chemical gradients of LPC (22:6) and FA (20:5). Interestingly, we observed several lipids and metabolites enhanced in the cell clusters at the islet periphery as shown in Fig. S57, including LPA (18:0), LPS (18:0), and LPC (18:1). Their spatial distribution only partially colocalizes with α-cells, indicating there might be a subpopulation of cells enriched in these lipids. This highlights the potential of nano-DESI MSI in resolving cell subtypes within the tissue.

**Fig. 5.**
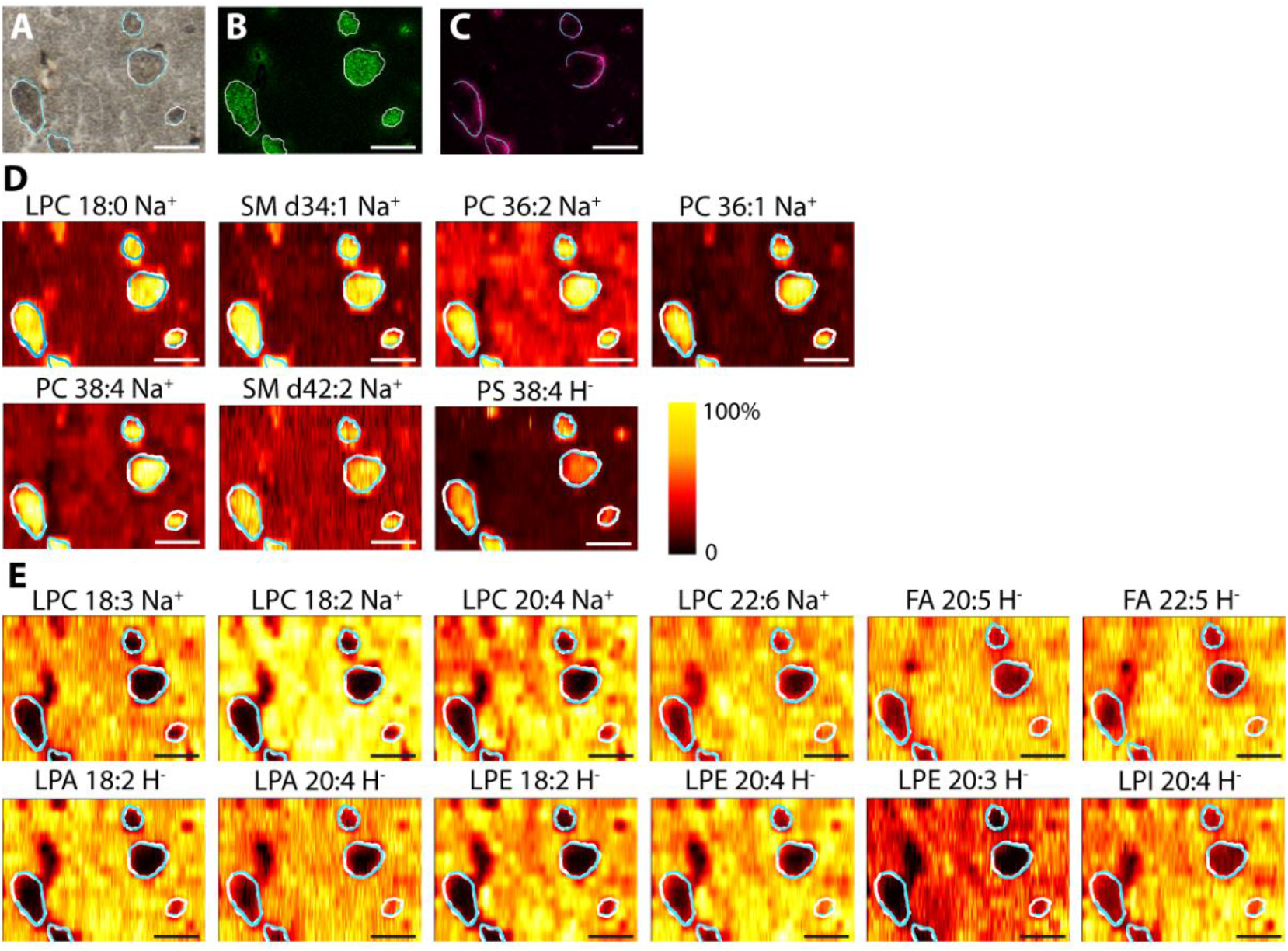
Ion images of lipids and metabolites on mouse pancreas. (A) Brightfield optical image of the analyzed region of the pancreatic tissue section. IF image of (B) insulin and (C) glucagon on the adjacent section. Ion images of lipids and metabolites normalized to TIC with (D) species showing enhanced abundance in endocrine cells and (E) species showing diminished abundance in islets of Langerhans. Scale bar: 250 µm. White trace delineates the location of endocrine cells based on the IF image of insulin in panel (B); light blue trace delineates the location of α-cells based on the IF image of glucagon in panel (C).

Collectively, our results demonstrate that the powerful combination of IF microscopy with high-resolution nano-DESI MSI provides unique insights into the distribution of lipids, metabolites, and peptides in different cell types in biological tissues. By highlighting cell types using IF microscopy, we resolved molecular features of specific cell types in both endocrine and exocrine regions of pancreatic tissues and identified species that may be used to understand biochemical pathways in this complex system.

## Discussion

The emergence of spatial multi-omics is critical to advancing our understanding of biological systems. Given the intricate connectivity and complexity inherent in these systems, no singular technique can provide a comprehensive description of key biochemical pathways. Consequently, the development of multimodal methodologies, such as the integration of corelative nano-DESI MSI imaging with IF microscopy presented in this study, is a burgeoning frontier in research. The approach developed in this study provides spatially-resolved cell-type-specific maps of a broad range of biomolecules, including lipids, metabolites, peptides, and proteins in the same tissue sample, which substantially expands the molecular coverage of MSI approaches. Using this approach, we discovered a series of peptides likely involved in endocrine-exocrine communication and measured chemical gradients of hundreds of biomolecules across different cell types.

A majority of lipid species observed in this study are tightly localized to specific cell types (Fig. 5). We hypothesize that these lipids are components of cell membranes. Although lipids typically suppress signals of other biomolecules in MSI, we observed many peptides in negative ionization mode in the same imaging experiment alongside lipids (Fig. 4). On-tissue MS/MS capabilities of nano-DESI MSI allowed us to identify some peptides that have not been previously reported in literature. Specifically, we identified four truncated analogs of C-peptide 1 and ten truncated analogs of C-peptide 2 along with the two intact isoforms. Sequence similarity between these truncated peptides that differ only by several C-terminal residues makes it difficult to explore their localization using antibody-based imaging techniques. Our results highlight the power of nano-DESI MSI for mapping and top-down identification of peptides and protein products that cannot be examined using other techniques.^66^

Truncated peptides have been observed in both healthy and diseased states of biological systems. For example, elevated levels of ubiquitin and β-thymosins truncated at the C-terminus in cancerous tissues have been used as markers of tumor proliferation.^66,67^ Insulin and C-peptide are produced in equimolar amounts during proinsulin conversion in β-cells. However, an elevated ratio of insulin to C-peptide was observed in rat β-cell extracts, which was attributed to either degradation of C-peptides or impaired cleavage of proinsulin.^68^ Truncated C-peptides can be produced either from proinsulin or as degradation products generated by β-cells.^58,69^ Truncation of C-peptides in β-cells of rat pancreas was first reported in 1973.^55^ The discovered truncated form, des-(23-31)-C-peptide 2, was suggested to be cleaved prior to and liberated during proinsulin conversion.^55^ A later study reported another truncated form, des-(27-31)-C-peptide 2, in rat β-cells and suggested that it could be either produced through the degradation of free C-peptide 2 or generated during proinsulin conversion.^58^ Their different mode of generation indicates that they may be involved in different pathways and serve as functional units in different biological processes.^52,58^ It has been demonstrated that C-peptide truncation varies in different species. For example, des-(27-31)-C-peptide 2 is an abundant truncated form comprising ∼10%-37% of the total C-peptide in rat β-cells,^58^ but only ∼1.5% or less in healthy human samples.^70^ Although the role of truncated C-peptides in pancreatic biology is not fully understood, there is a growing evidence of their involvement in both diabetes and pancreatic cancers.^71^ In clinical studies, C-peptide test has been used to distinguish between type 1 and type 2 diabetes.^72^ In fact, several truncated C-peptides proteolytically cleaved at positions 21, 24, 26, and 30 have been identified in human insulinoma cells.^71^ Understanding the roles of truncated C-peptides in diseased tissues could provide valuable insights into the disease mechanism and related metabolism pathways.

It is remarkable that the multimodal nano-DESI MSI approach allowed us to both identify 16 forms of C-peptides in mouse pancreatic tissue and examine their spatial localization and chemical gradients across different cell types. A majority of C-peptides are expressed both in the endocrine tissue and acinar cells in the peri-insular regions (Fig. 4). Exocrine cells are located in close proximity to endocrine cells in pancreas and likely influence their functional state.^73,74^ Several studies have emphasized the influence of endocrine-exocrine crosstalk on the proliferation and health of β-cells.^51^ For example, pancreatic enzymes, elastase and carboxyl ester lipase, secreted by exocrine cells influence endocrine cell function, proliferation, and longevity.^75^ Conversely, peptides secreted by endocrine cells have been shown to support exocrine cell function through the insulo-acinar portal system.^47^ Several well-studied endocrine hormones play a pivotal role in promoting (pancreatic polypeptide), inhibiting (glucagon, somatostatin, and amylin), and regulating (insulin, ghrelin and cholecystokinin) exocrine enzyme secretion.^47^ Moreover, pathological conditions in the exocrine tissue can cause dysfunction in endocrine tissue and vice versa. For example, exocrine cell loss and dysfunction was observed in type 1 diabetic tissue and attributed to endocrine β-cell insulin deficiency. Meanwhile, endocrine/β-cell dysfunction can be induced by exocrine pancreas damage in type 3c diabetes.^73,75^ C-peptides are originally produced from proinsulin in β-cells. The observed shallow chemical gradients of C-peptides extending from the islets to exocrine cells in the peri-insular regions suggest that C-peptides are transported from β-cells to exocrine cells. This indicates that they may be involved in intracellular processes related to endocrine-exocrine communication. It has been reported that endocrine cells communicate with peripheral lymphoid tissue through exocytosis of β-cell peptides.^76^ Several proinsulin peptides, including B chain segments, truncated C-peptide segments, and hybrid insulin peptides (HIP) comprised of C-peptide fragments fused to other β-cell granule peptides have been identified as autoantigens presented by class II major histocompatibility complex (MHC-II) molecules on antigen-presenting cells (APCs). Autoimmune responses in type 1 diabetes can be activated when these peptides are recognized by islet-infiltrating helper T cells.^76–80^ Additionally, it has been proposed that MHC-II species can present proinsulin-derived islet antigens to regulatory T cells, providing protection against type I diabetes and other autoimmune diseases.^81^ In fact, a short peptide epitope C19-A3, which is a proinsulin C-A chain junction containing the C-terminal half of the C-peptide, has been tested for immunotherapy of type 1 diabetes to restore immune tolerance and modify autoimmune response, which efficiently preserved β-cell mass.^82,83^ The concentration gradients of autoangiogenic C-peptides in healthy pancreatic tissues reported in this study may be indicative of such protection. We hypothesize that the observed differences in chemical gradients of truncated C-peptides may be indicative of differences in their functions. Future research will use the experimental platform described in this study to identify additional peptides and investigate their abundance and localization in both healthy and diseased pancreatic tissues. These efforts will facilitate understanding of the role of peptides in endocrine-exocrine intracellular signaling pathways in the pancreas of both healthy and diseased states.

The multimodal imaging approach described in this study can be further improved to enhance the molecular coverage, sensitivity, chemical specificity, and spatial resolution. Several strategies may be used for increasing the molecular coverage and sensitivity of nano-DESI MSI. These include further optimization of the composition of the extraction solvent to enhance signals of low-abundance species,^25,84^ coupling of nano-DESI MSI with ion mobility separation to improve molecular coverage and facilitate identification,^30,85,86^ online derivatization to improve the ionization efficiency of poorly-ionizable analytes,^87^ and individual ion detection to improve sensitivity towards protein detection.^28^ Chemical specificity can be improved by integrating on-line chemical reactions with MSI to resolve lipid isomers.^88,89^ Although the currently achievable spatial resolution of nano-DESI MSI is sufficient to visualize molecular features in cell clusters of a specific type, further improvement is required to enable imaging at a subcellular level necessary to understand cellular heterogeneity and intracellular interactions. Higher spatial resolution may be achieved by employing computational methods, such as image fusion for high spatial resolution prediction.^17,18,20,22^ Furthermore, coupling nano-DESI MSI to other imaging modalities such as Raman microscopy or histology staining may provide additional biochemical information about the system.^22,90^

## Conclusions

The experimental platform reported in this study combines correlative nano-DESI MSI of small molecules and proteins with IF imaging for identifying different cell types and their molecular signatures in biological tissues. This platform enables spatial multi-omics analysis of biological tissue sections with cellular-level spatial resolution. A comparison of nano-DESI MSI and IF data indicates that lipids and some proteins are excellent markers of different cell types. Meanwhile, chemical gradients observed for peptides provide insights into endocrine-exocrine communication. Although the combination of IF and MSI is a powerful tool for studying cell-specific molecular signatures in biological tissues, it is often difficult to perform both analyses on the same sample. Our results indicate that lipids can be used as cell markers in mouse pancreas, which will simplify the experimental workflow and enable cell-specific multi-omics analysis of the same tissue section. Further improvement of this platform and its application to diseased tissues, such as diabetic pancreas, could provide insights into the molecular phenotype, disease mechanism and progression, and help identify preclinical diagnostic biomarkers and therapeutic targets. The correlative nano-DESI MSI approach and the experimental findings reported in this study open several research directions both in pancreatic biology and chemical imaging.

## Experimental

### Chemicals and materials

HPLC grade water, methanol (MeOH), acetonitrile (ACN) and optima LC/MS Grade formic acid (HCOOH) were purchased from Fisher chemical (Hampton, NH). Ethanol 200 proof was obtained from Decon Laboratories, INC (King of Prussia, PA). 99.8+% chloroform was purchased from Alfa Aesar (Tewksbury, MA). Polymicro flexible fused silica capillary tubing (OD 150 μm, ID 50 μm; OD 360 μm, ID 100 μm; and OD 800 μm, ID 200 μm) was ordered from Molex (Thief River Falls, MN). CoraLite^®^ plus 488 INS mouse monoclonal antibody (Cat# CL488-66198) and CoraLite^®^ plus 647 glucagon rabbit polyclonal antibody (Cat# CL647-15954) were purchased from Proteintech (Rosemont, IL). DAPI (4’,6-Diamidino-2-Phenylindole, Dihydrochloride) (Cat# Invitrogen™ D1306) was purchased from Thermo Fisher Scientific. Fluoromount™ was purchased from Diagnostic BioSystems Inc. (Pleasanton, CA).

### Animal tissue preparation

Pancreas of wild type male C57BL/6 mice (RRID: IMSR_JAC: 000664) were used in this study. The mice were housed and euthanized following protocol #1511001324 as approved by the Purdue institutional Animal Care and Use committee (IACUC). Adult, 3−4-month-old mice were euthanized by CO_2_ inhalation followed by cervical dislocation, after which their pancreas were collected and snap frozen in isopentane kept over dry ice.^91,92^ Tissues were sectioned at −21 °C using a CM1860 Cryostat (Leica Microsystems, Wetzlar, Germany) to 18 μm and thaw-mounted onto glass microscope slides (IMEB, Inc. Tek-Select Gold Series Microscope Slides, Clear Glass, Positive Charged). The sections were stored in a −80 °C freezer until use.

### Immunofluorescence imaging

Path Scan Enabler IV (Meyer Instruments, Inc., Houston, TX) was used to acquire wild-field optical images of the pancreatic tissue sections. Subsequently, the sections were stained for IF imaging of insulin and glucagon. Sections were fixed in 4% paraformaldehyde (in PBS) for 10 mins, quenched in 100 mM glycine for 10 mins and incubated in blocking buffer (5% goat serum, 2% bovine serum albumin, 0.1% Triton X-100 and 0.1% sodium azide in PBS) for 1 h at room temperature. Tissue sections were then incubated in CoraLite^®^ plus 488 conjugated insulin primary antibody with CoraLite^®^ plus 647 conjugated glucagon primary antibody (1:500 for both antibodies in blocking buffer) overnight at 4 °C. Next, tissue sections were incubated with DAPI (1:1000 in PBS) for 10 mins at room temperature. Each slide was prepared by adding 1-2 drops of fluorophore mounting media, followed by the precise sealing of a coverslip using nail polish. IF images were acquired using a Nikon A1Rsi confocal microscope (Nikon, Tokyo, Japan). Insulin, glucagon and nuclei were visualized using green (excitation: 488 nm, emission: 500-550 nm), far-red (excitation: 640 nm, emission: 663-738 nm), and blue (excitation: 405 nm, emission: 425-475 nm) channels, respectively.

### Multimodal Nano-DESI MSI

Nano-DESI MSI of lipids and metabolites were performed on a Q-Exactive HF-X Orbitrap mass spectrometer (Thermo Fisher Scientific, Waltham, MA) equipped with a custom-designed nano-DESI source described in our previous publications.^23,27,31^ Briefly, a high-resolution glass capillary probe was assembled using two fused silica capillaries pulled down to ∼10 µm in diameter. The probe was positioned at a ∼90-degree angle in front of the mass spectrometer inlet. Tissue samples were placed on a motorized XYZ stage controlled by a custom-designed LabVIEW program^23^ and scanned under the nano-DESI probe at a scan rate of 15 µm/s.

The instrument was tuned and calibrated using the Pierce™ LTQ Velos ESI positive ion calibration solution and Pierce™ ESI negative ion calibration solution. MeOH/H_2_O (9/1, *v/v*) was used as the extraction solvent for lipids and metabolites and the solvent flow rate was 0.5 µL/min. The capillary temperature was 300 °C; mass range was *m/z* 133-1700; mass resolving power was 60,000 (m/Δm at *m/z* 200); the AGC target was 1E6; the maximum injection time was 200 ms. Line-scans were acquired in alternate polarity, with all odd lines in positive ionization mode and all even lines in negative ionization mode. The high voltage was 4.5 kV for positive and -3.5 kV for negative ionization mode. The step between the lines was 30 µm.

After nano-DESI MSI of lipids and metabolites, tissue sections were immersed into serial ethanol solutions (70, 90, and 100%) for 20 s each, followed by washing in 99.8% chloroform for 25 s.^27^ Nano-DESI MSI of proteins was then performed on the same region of the tissue section using an Agilent 6560 ion mobility (IM) quadrupole time-of-flight (Q-TOF) instrument.^30^ The system tune and mass calibration were performed using Agilent ESI-L low concentration tune mix/0.1 mM HP-0321/ACN/H_2_O (10 mL/3 µL/85.5 mL/4.5 mL). Protein imaging data were acquired in positive ionization mode with a mass range of *m/z* 300-1800 and ACN/H_2_O/FA (80/20/0.07, *v/v/v*) as extraction solvent delivered at a flow rate of 0.5 µL/min. The gas temperature was 300 °C; capillary voltage was 4 kV. The Quad AMU was 200 amu. High pressure funnel RF, trap funnel RF and rear funnel RF were 180 V, 200 V and 180 V, respectively. The acquisition rate was 1 Hz. The step between the lines was 60 µm.

### Tandem mass spectrometry (MS/MS) data acquisition and analysis

On-tissue MS/MS experiments were performed on the Q-Exactive HF-X Orbitrap mass spectrometer using higher-energy collisional dissociation (HCD). MS/MS spectra were acquired by scanning the nano-DESI probe across a region of interest (e.g. endocrine, exocrine, and blood vessel regions) in tissue samples.^27^ Peptides detected in negative ionization mode were fragmented using normalized collisional energies of 20, 25, and 30 arbitrary units. The mass resolution was 60,000 and mass isolation window was 3.0 Da. Proteins and peptides detected in positive ionization mode were fragmented using normalized collisional energies of 15, 20, and 25 arbitrary units. The mass resolution was 60,000 and mass isolation window was 2.0 Da. The AGC target was 1E5 and the mass range of *m/z* 133-1999 was used for both modes.

Deprotonated peptide fragmentation spectra were manually annotated by matching accurate *m/z* of the precursor ions and *m/z* of fragments to the predicted *m/z* of b, c, y ions and the corresponding neutral loss fragments of candidate sequences. Matched fragments with mass error<0.005 Da are annotated in the fragmentation spectra and fragmentation maps. For the identification of positively charged proteins/peptides, the .RAW files were processed using ProSightPD 4.0 for Proteome Discoverer v2.5 (PSPD, Thermo Scientific). The PSPD Comprehensive Discovery Proteomics with FDR analysis template was used. The workflow contains three search nodes: an annotated proteoform search with a 2.2 Da precursor tolerance, an annotated proteoform search with a 100 Da precursor tolerance, and a subsequence search with a 10 ppm precursor tolerance. All three search nodes used a Mus musculus database created from UniProt release 2020_04 (https://www.uniprot.org/proteomes/UP000000589). The workflow also included a false discovery rate (FDR) calculation. Results that passed the 1% FDR cutoff were annotated using PFR Accessions and included in the output files in .pdResult and .tdReport formats. TDViewer (http://tdviewer2.northwestern.edu) was used to review the results in the .tdReport file. Proteoform identifications were manually validated by examining the diagnostic fragments in the MS/MS spectra using TDValidator (Proteinaceous), including the isotopic profile, mass error<10 ppm and S/N>3. We note that *m/z* of proteins and peptides reported in this study were calculated using the most abundant isotopic peak.

### Image generation and registration

Ion images obtained from Q-Exactive HF-X Orbitrap mass spectrometer were generated using a custom Python script (https://github.com/LabLaskin/MSIGen). This script uses the pymsfilereader package to extract data from .RAW files. Mass spectral data were split using scan filters and aligned according to scan times. A mass window of 10 ppm was used to extract signals of selected m/z values.^27^ Ion images obtained from Q-TOF instrument were generated from .d files using a previously described workflow.^93^ In this workflow, Skyline software is used to extract signal intensities of the most abundant isotopic peak of each charge state of a protein in each pixel into a .txt file. Ion images are then generated from the .txt file using the custom Python script. All the signals were normalized to the corresponding total ion current (TIC) to compensate for signal variations during the MSI experiment.^31^

Image registration was performed using a custom Python script that prompts the user to select at least three anchor points for each IF image. The affine transform that best aligned these points was used to align the image coordinate planes. Images were overlayed representing each image with different RGB channels after manually adjusting brightness of each constituent image.

## Author contributions

Conceptualization: MY, JL

Methodology: MY, FNM, ZQ, PS, JL

Funding acquisition: JL, ZYZ

Investigation: JL

Visualization: MY, MI, ELH

Supervision: JL

Writing—original draft: MY, MI, FNM, ELH, ZQ

Writing—review & editing: MY, JL, PS

## Data availability

All data needed to evaluate the conclusions in the paper are present in the paper and/or the Supplementary Materials.

## Acknowledgments

This work was partially supported by grants from the National Science Foundation (2108729 and IIP-1916691) and National Institutes of Health (RF1MH128866 and RO1CA069202). The authors acknowledge the use of the facilities of the Bindley Bioscience Center, a core facility of the NIH-funded Indiana Clinical and Translational Sciences Institute. The authors gratefully acknowledge Dr. Jiamin Qiu and Prof. Shihuan Kuang from Purdue University for providing reagents, resources, and help with immunofluorescence staining; Dr. Xiaoguang Zhu for training and help with confocal microscopes; Dr. Yunpeng Bai for providing mouse brain and kidney tissues for the initial testing of the methodology; Dr. Joseph B. Greer, Dr. Ryan T. Fellers, Dr. Bryan P. Early, and Prof. Neil L. Kelleher for help with the top-down proteomics database searching. Part of Figure 1 was created with BioRender.com under the agreement number SD26QWVRTS.

