## Supplemental Materials for "Correlative Imaging for Comprehensive Molecular Mapping of Individual Cell Types in Biological Tissues"

Figure S1: Ion images of insulin proteoforms.
Figure S2-S6: MS/MS spectra with annotated fragments and fragmentation maps of the identified proteins/peptides.
Figure S7-S14: MS/MS spectra of the unidentified proteins/peptides.
Figure S15: Examination of protein delocalization after nano-DESI MSI.
Figure S16-S18: Ion images of pancreatic peptides obtained from three biological replicates.
Figure S19: Extracted line profiles of peptides from ion images and extracted line profiles of insulin and glucagon from IF images.
Figure S20-S33: MS/MS spectra with annotated fragments and fragmentation maps of the truncated C-peptides.
Figure S34-S53: MS/MS spectra of the unidentified deprotonated peptides.
Figure S54-S56: Ion images of lipids and metabolites on mouse pancreatic tissue section obtained from three biological replicates.
Figure S57: Ion images of species showing signal enhancement in cell clusters at the periphery of each islet.
Figure S58: Extracted line profiles of lipids and metabolites from ion images and extracted line profiles of insulin and glucagon from IF images.

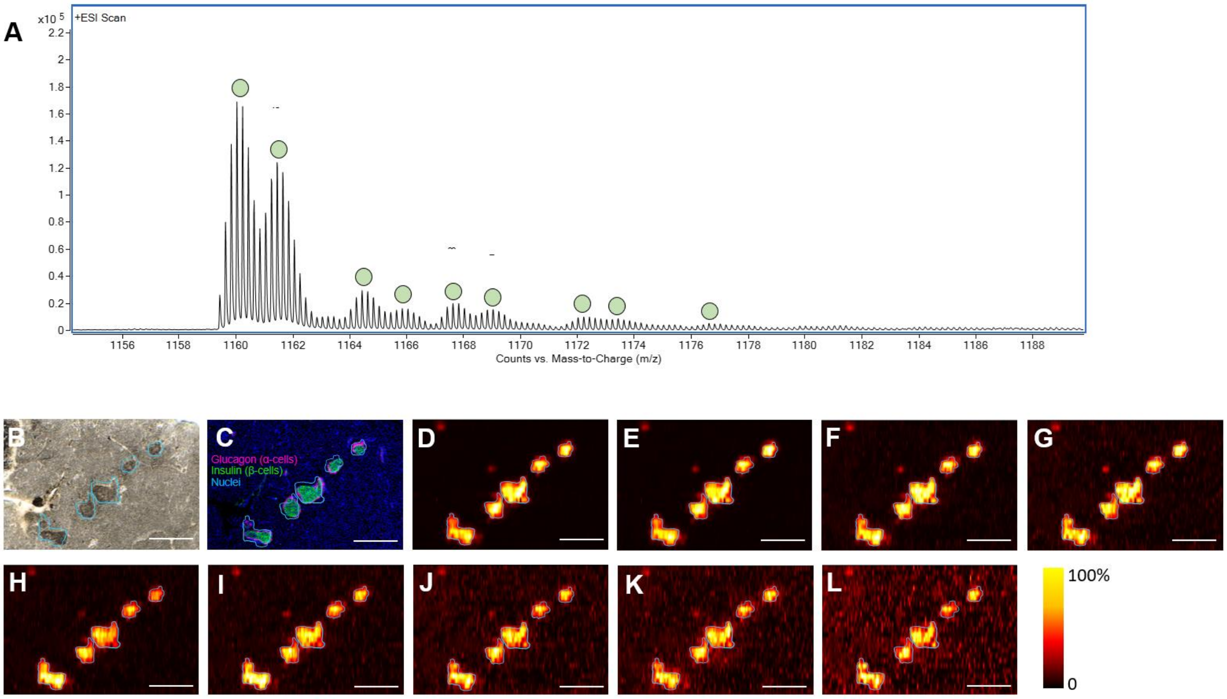


Fig. S1. Ion images of insulin proteoforms.

(A) Mass spectrum of the 5+ charge state of insulin proteoforms. Each proteoform is denoted by a green circular marker. (B) Brightfield optical image of the analyzed region of the pancreatic tissue section. (C) IF image of insulin (green), glucagon (pink), and DAPI (blue) obtained from the adjacent tissue section. Ion images of the 5+ charge state of the insulin proteoforms normalized to TIC: (D) *m/z* $1160.241^{5+}$, 5796 Da, insulin-2, (E) *m/z* $1161.656^{5+}$, 5803 Da, insulin-1, (F) *m/z* $1164.636^{5+}$, 5818 Da, (G) *m/z* $1166.049^{5+}$, 5825 Da, (H) *m/z* $1167.831^{5+}$, 5834 Da, (I) *m/z* $1169.130^{5+}$, 5841 Da, (J) *m/z* $1172.426^{5+}$, 5857 Da, (K) *m/z* $1173.635^{5+}$, 5863 Da, (L) *m/z* $1176.825^{5+}$, 5879 Da. Scale bar: 500 µm.

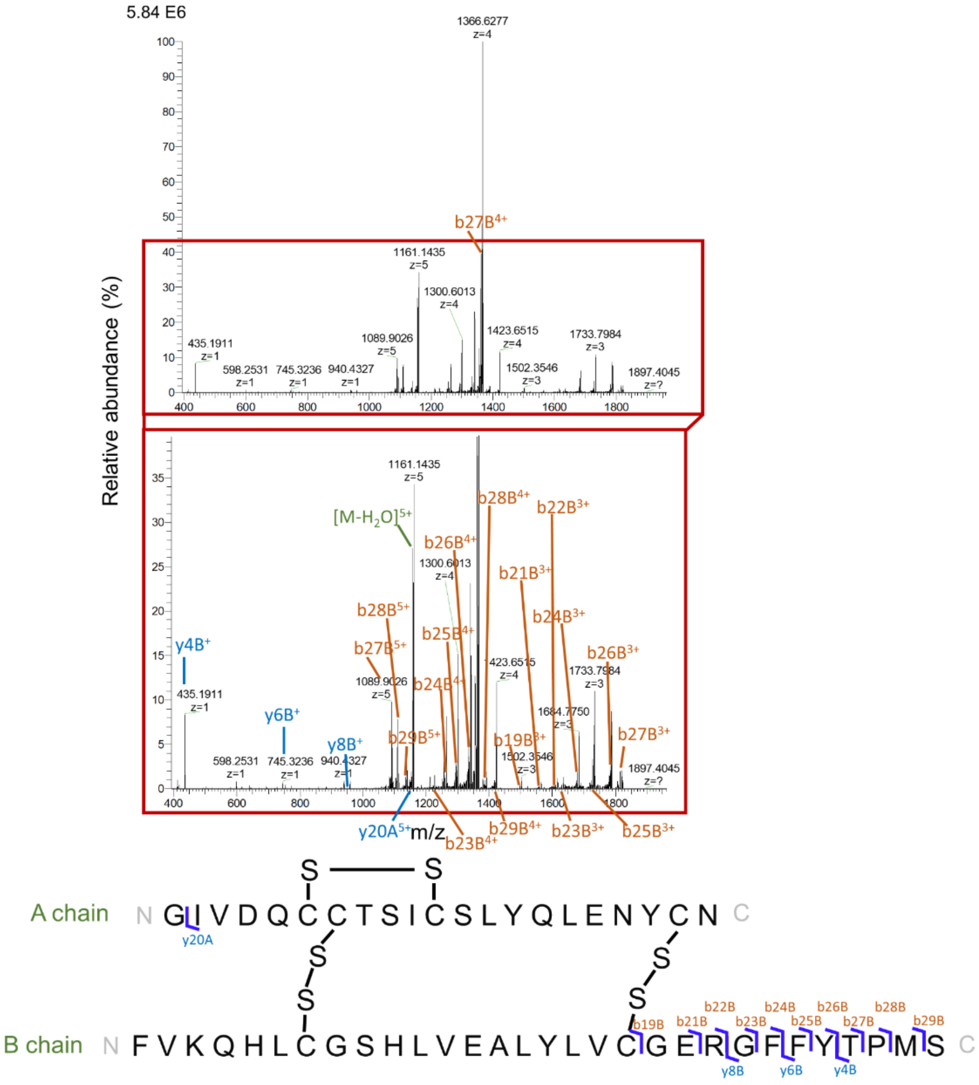

Fig. S2. MS/MS spectrum (top), zoomed-in MS/MS spectrum (middle) and the fragmentation map (bottom) of the +5 charge state of the 5796 Da insulin-2.

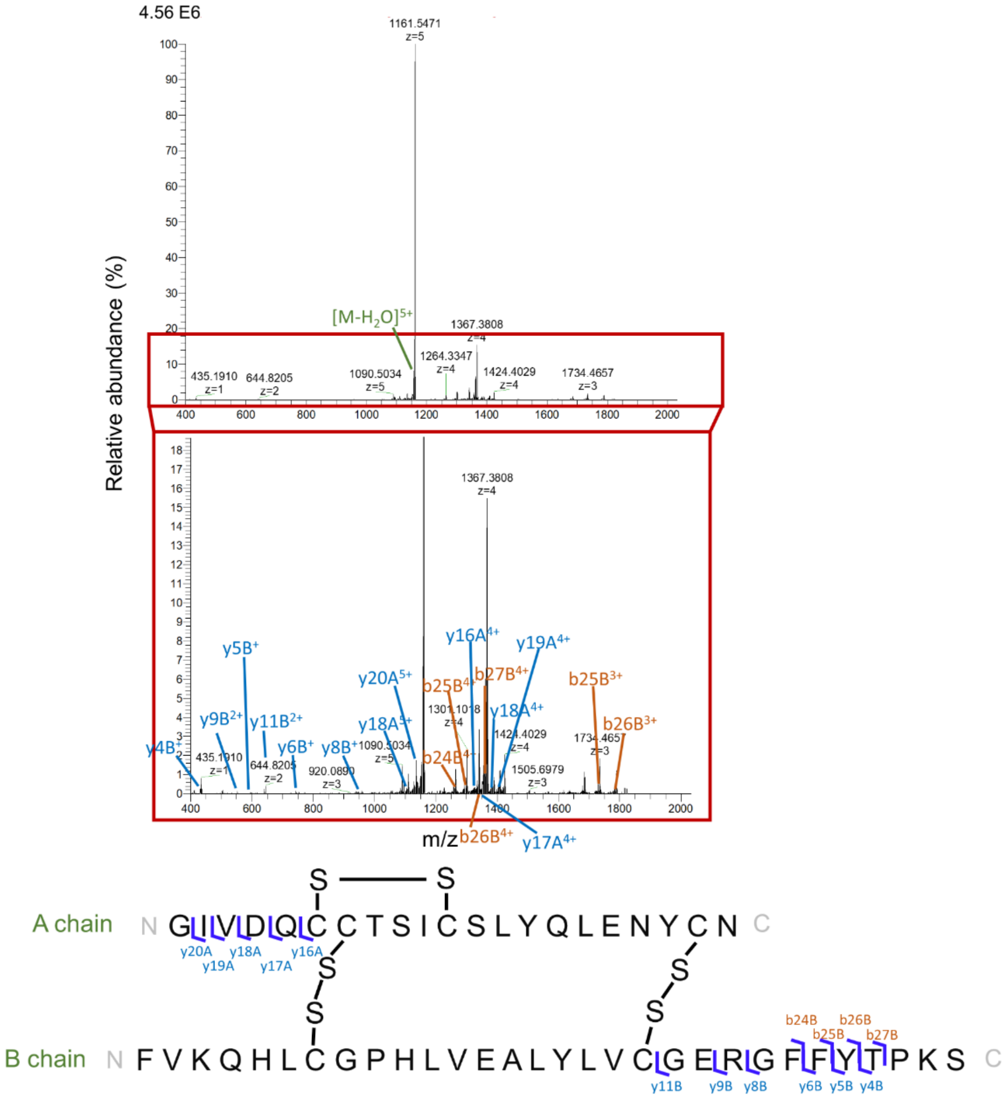


Fig. S3. MS/MS spectrum (top), zoomed-in MS/MS spectrum (middle) and the fragmentation map (bottom) of the +5 charge state of the 5803 Da insulin-1.

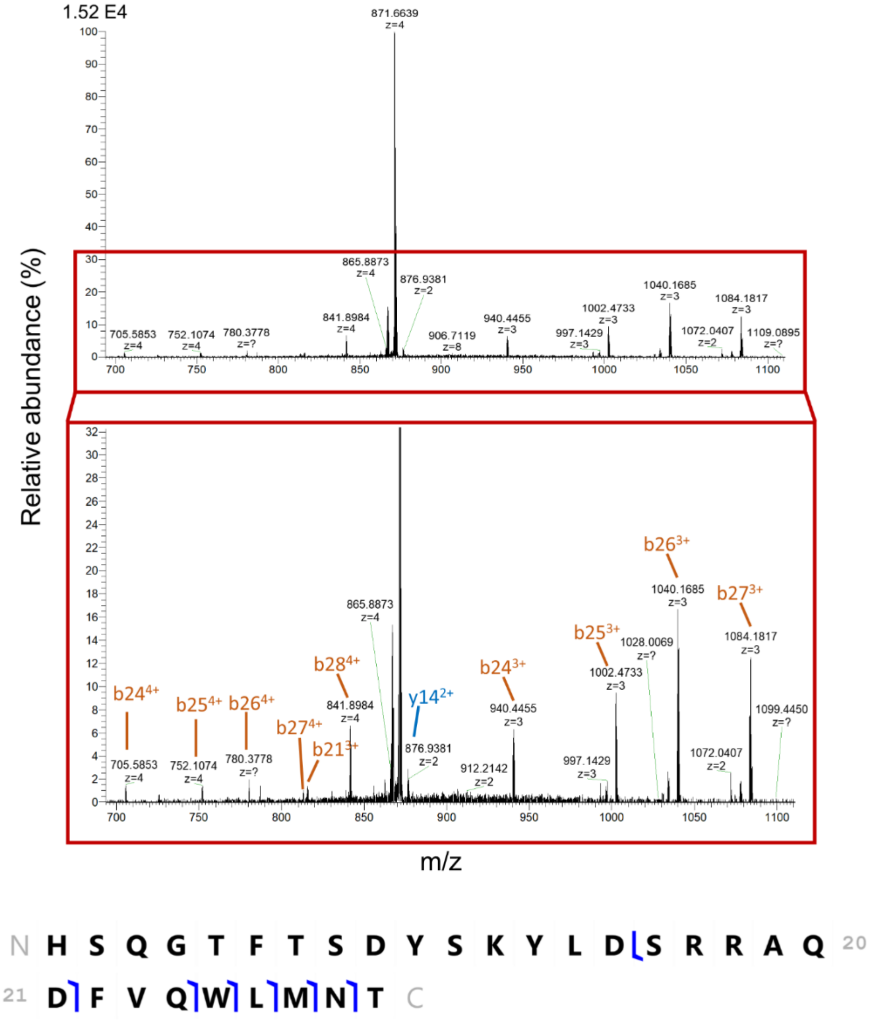


Fig. S4. MS/MS spectrum (top), zoomed-in MS/MS spectrum (middle) and the fragmentation map (bottom) of the +4 charge state of the 3482 Da glucagon.

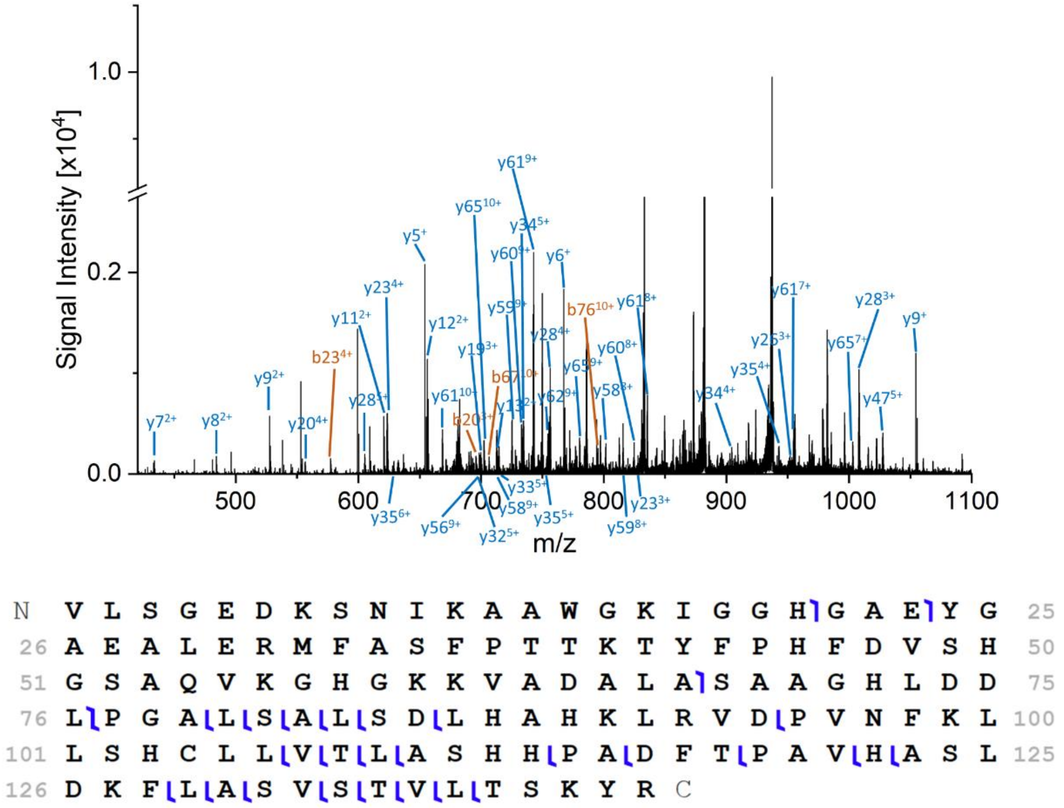


Fig. S5. The overlay MS/MS spectrum (top) and the fragmentation map (bottom) of the +16, +17, +18, +20, +22 charge state of the 14981 Da hemoglobin subunit alpha.

Note that the observed mass shifted +27 Da compared to the theoretical mass, this could be due to unresolved unknown post-translational modifications.

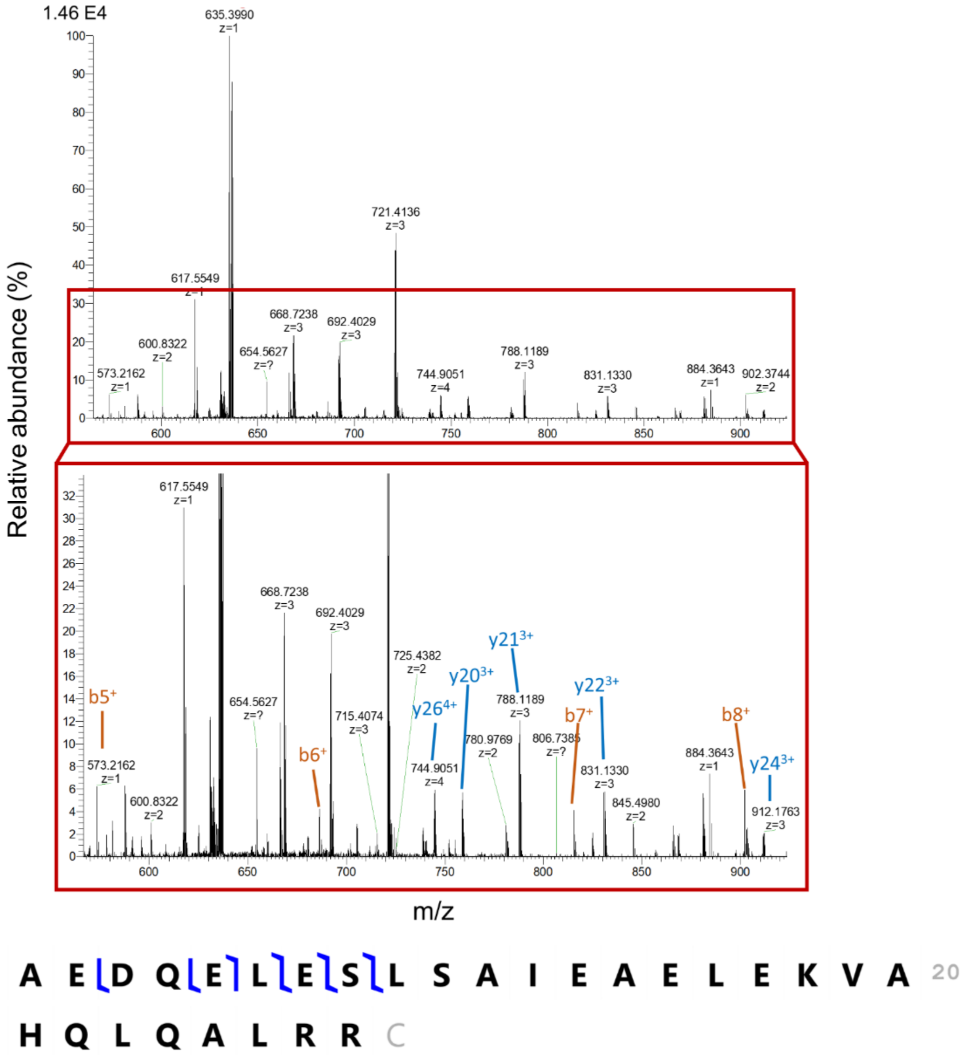


Fig. S6. MS/MS spectrum (top), zoomed-in MS/MS spectrum (middle) and the fragmentation map (bottom) of the +5 charge state of the 3177 Da serpinin-RR.

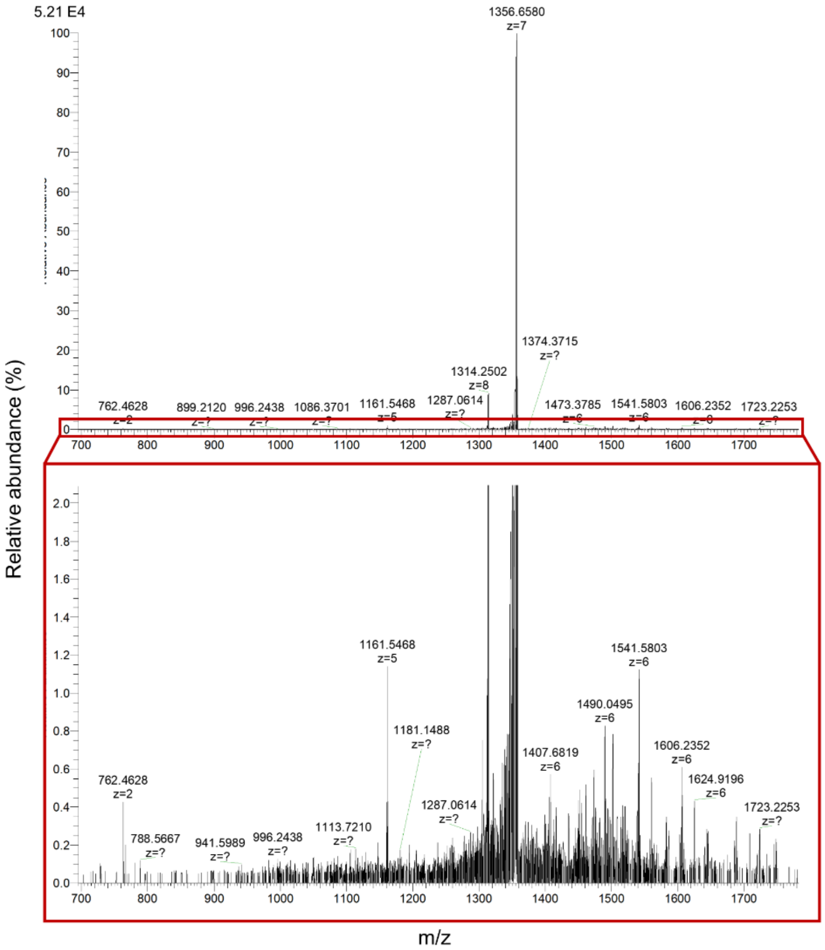


Fig. S7. MS/MS spectrum (top) and the zoomed-in MS/MS spectrum (bottom) of the +7 charge state of the 9490 Da protein.

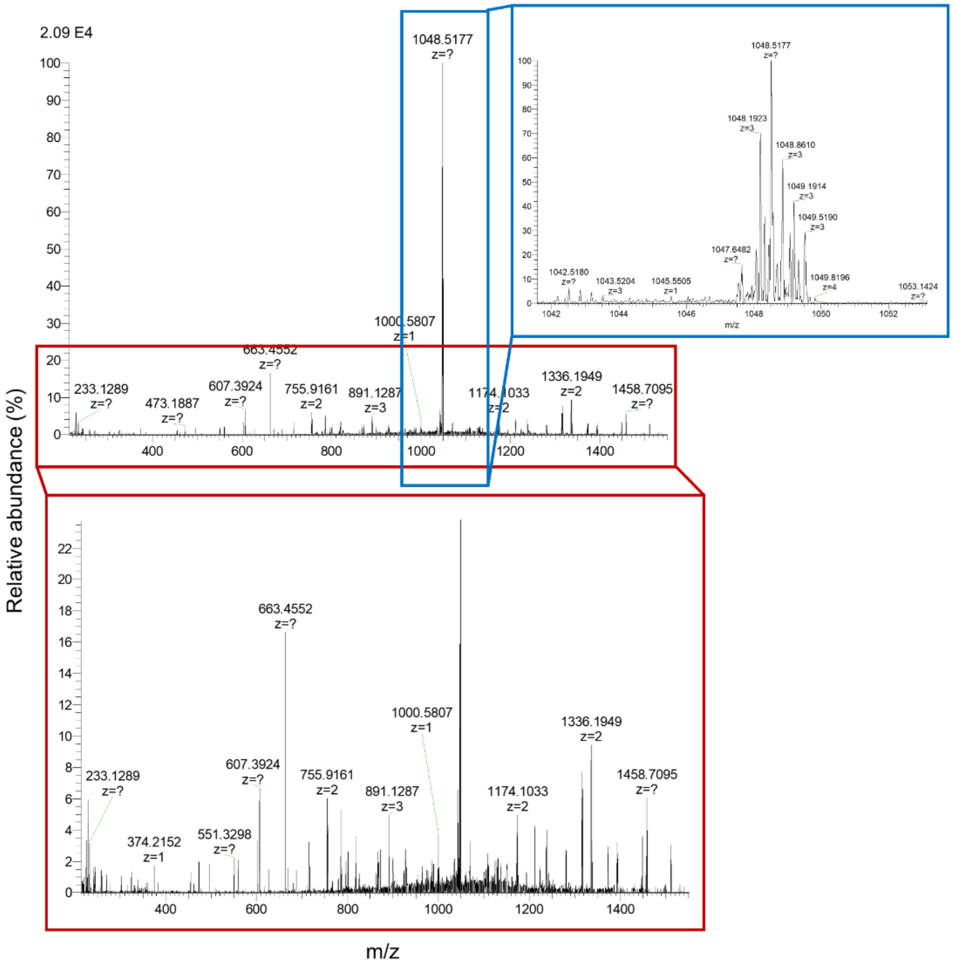


Fig. S8. MS/MS spectrum (top), the zoomed-in precursor peak (side) and the zoomed-in MS/MS spectrum (bottom) of the +3 charge state of the 3143 Da protein.

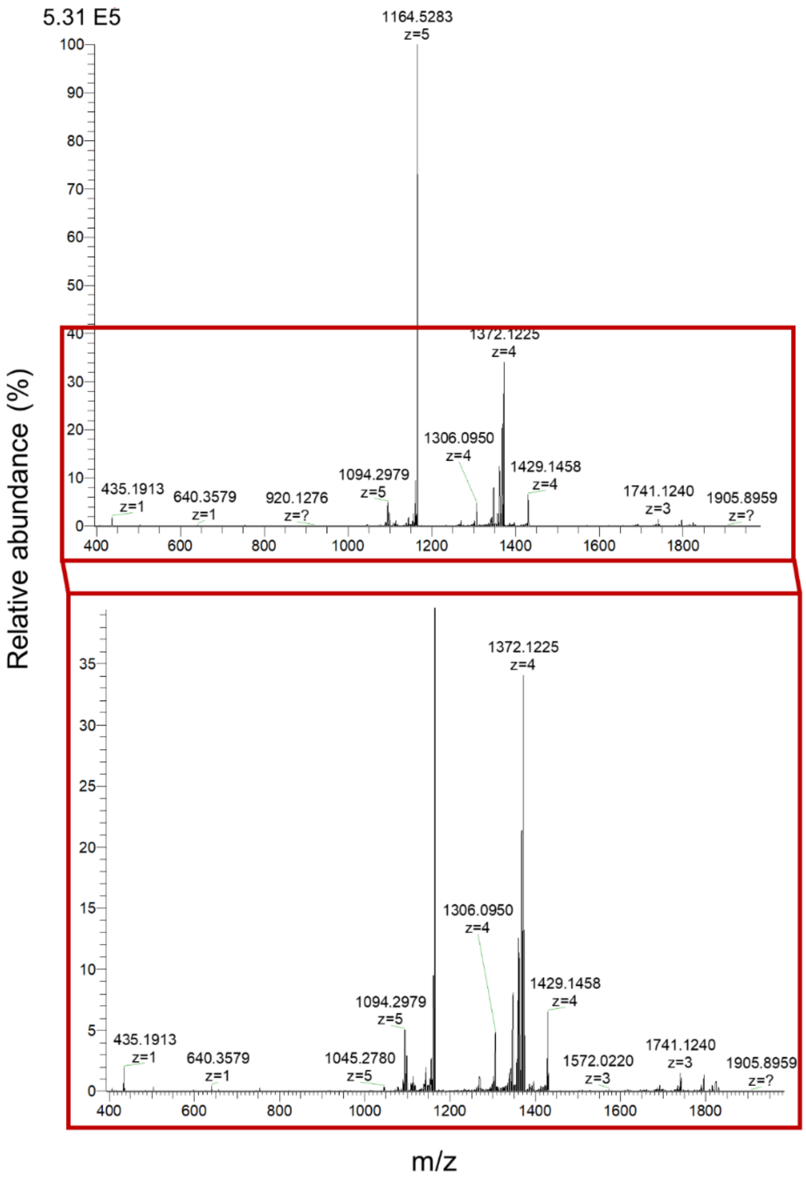


Fig. S9. MS/MS spectrum (top) and the zoomed-in MS/MS spectrum (bottom) of the +5 charge state of the 5818 Da insulin proteoform.

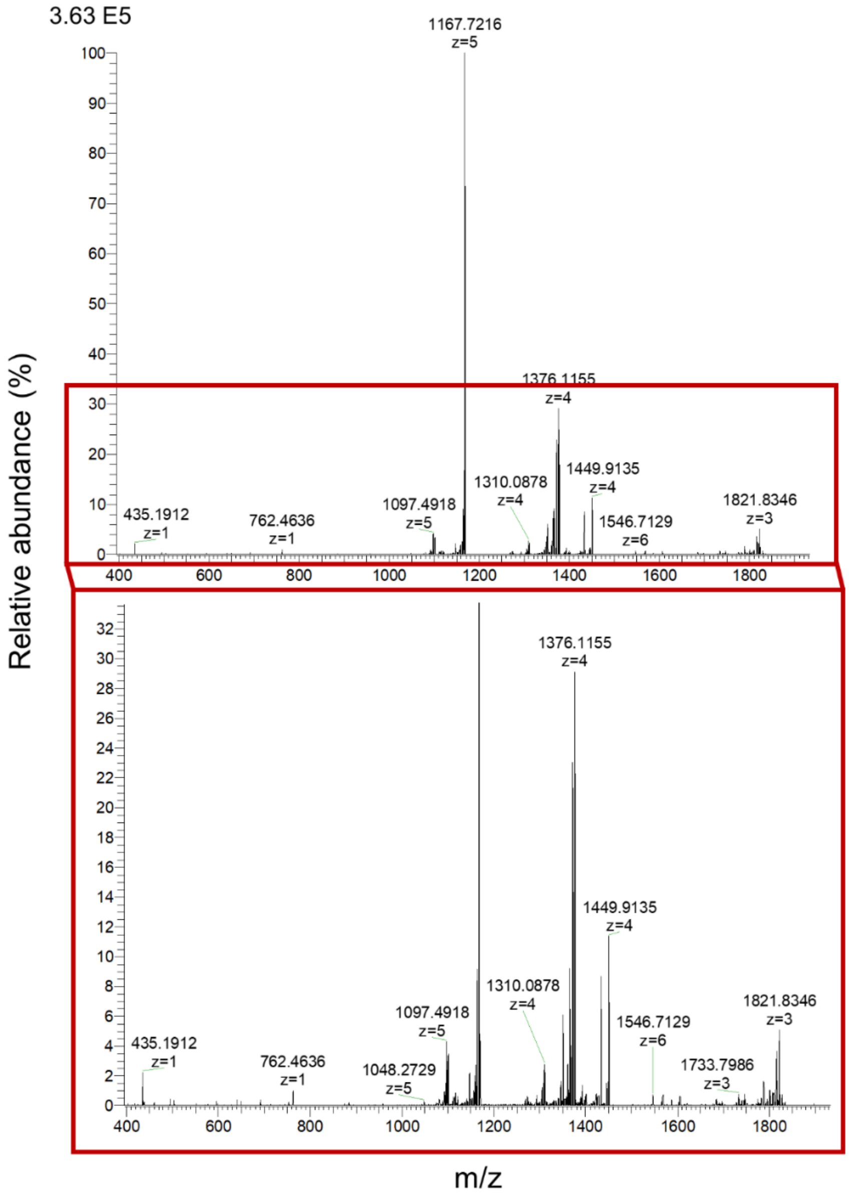


Fig. S10. MS/MS spectrum (top) and the zoomed-in MS/MS spectrum (bottom) of the +5 charge state of the 5834 Da insulin proteoform.

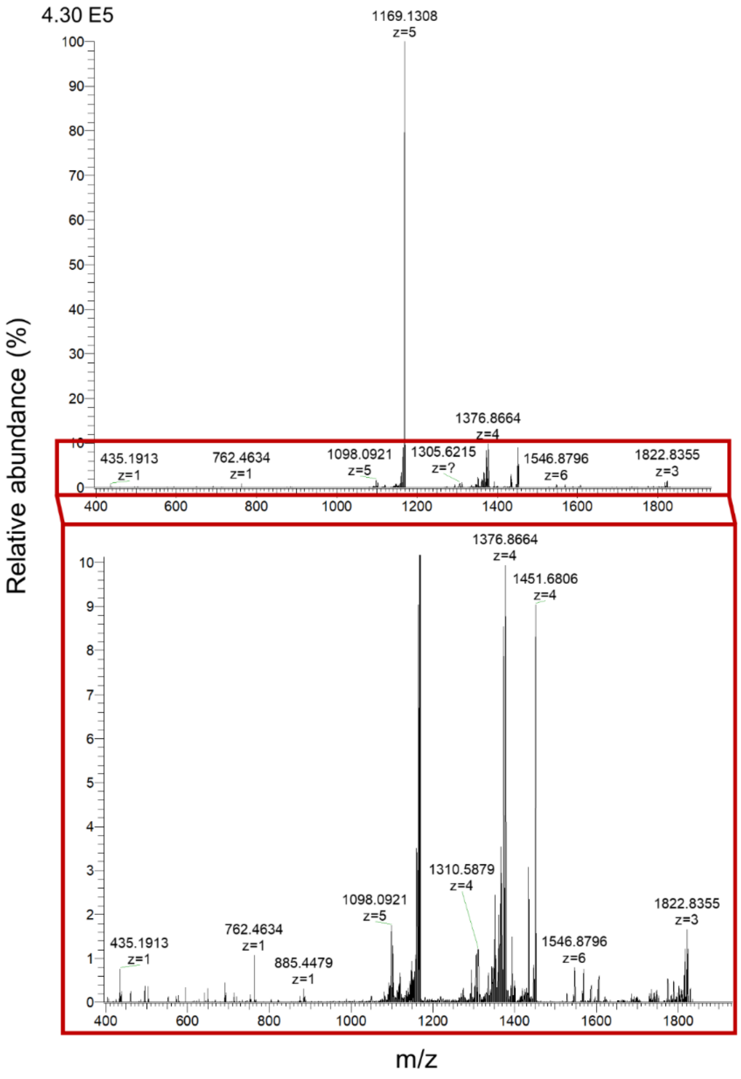


Fig. S11. MS/MS spectrum (top) and the zoomed-in MS/MS spectrum (bottom) of the +5 charge state of the 5841 Da insulin proteoform.

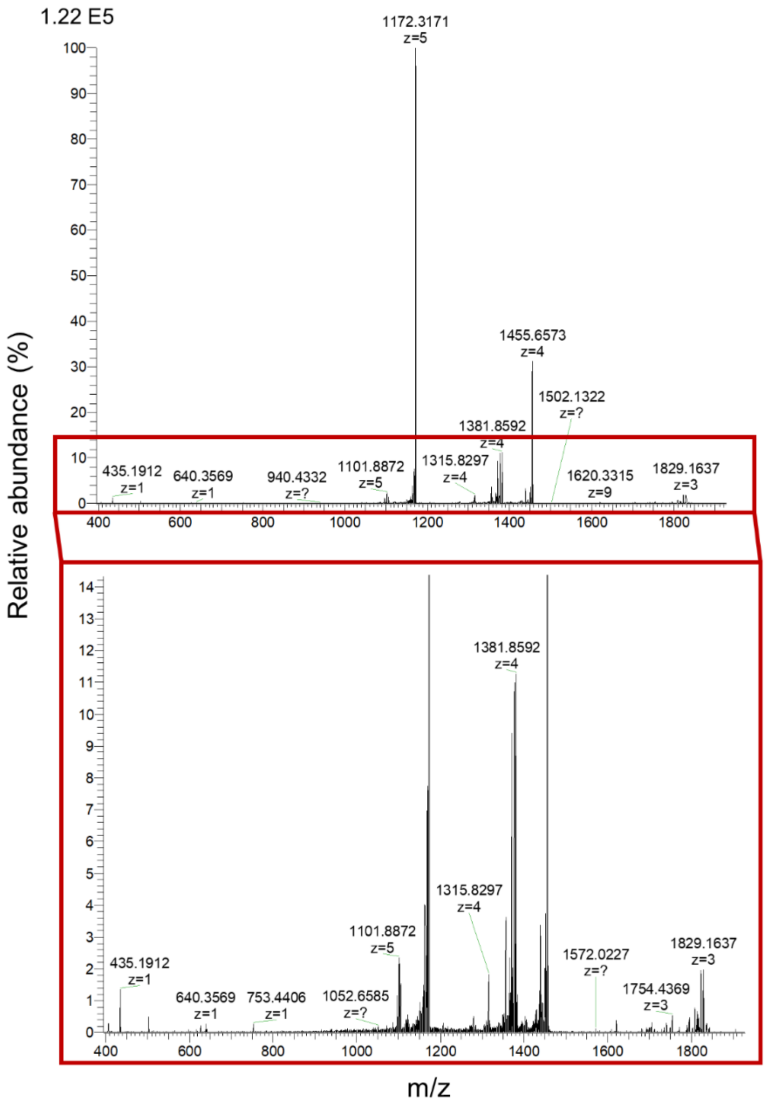


Fig. S12. MS/MS spectrum (top) and the zoomed-in MS/MS spectrum (bottom) of the +5 charge state of the 5857 Da insulin proteoform.

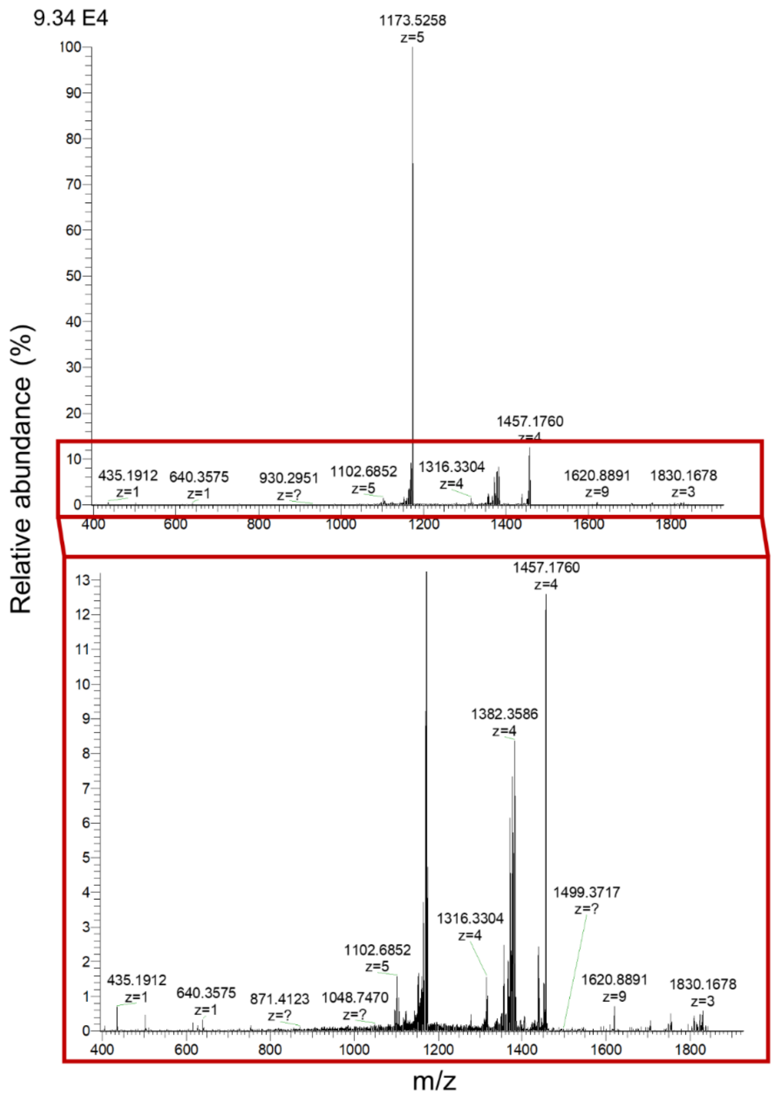


Fig. S13. MS/MS spectrum (top) and the zoomed-in MS/MS spectrum (bottom) of the +5 charge state of the 5863 Da insulin proteoform.

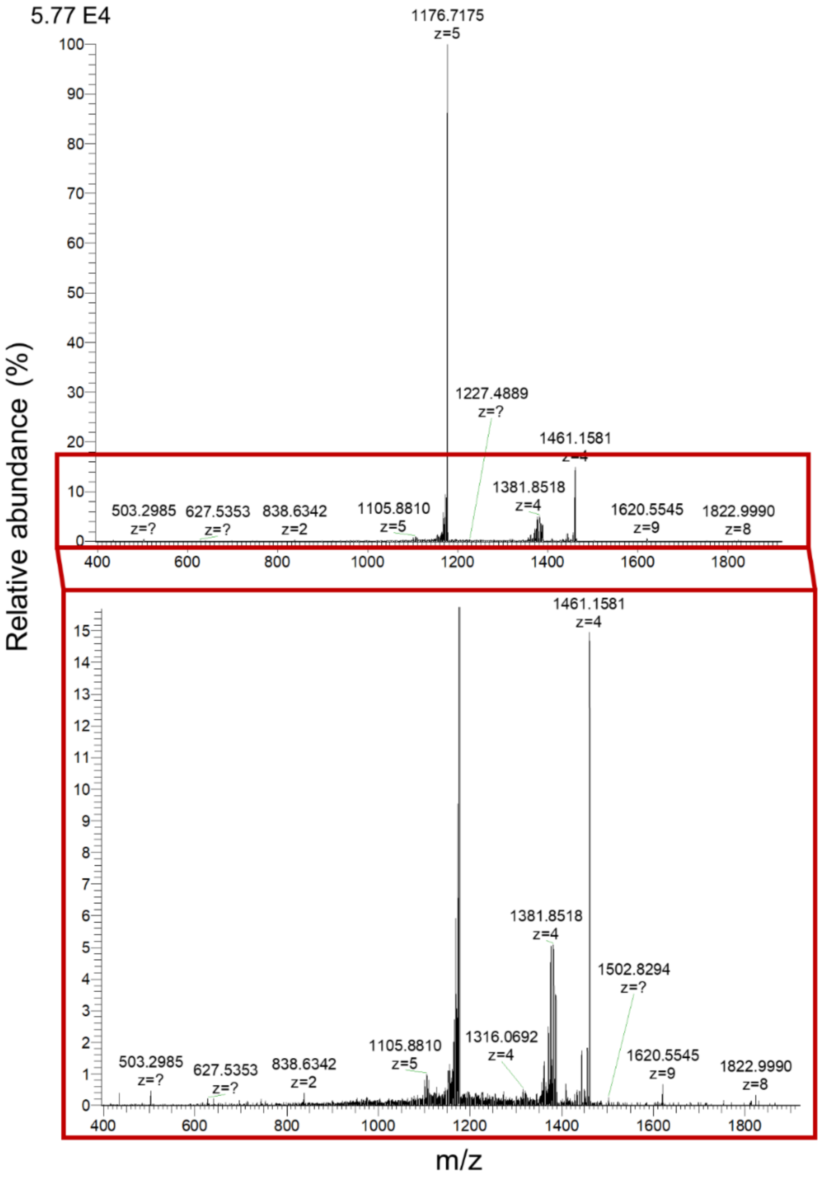


Fig. S14. MS/MS spectrum (top) and the zoomed-in MS/MS spectrum (bottom) of the +5 charge state of the 5879 Da insulin proteoform.

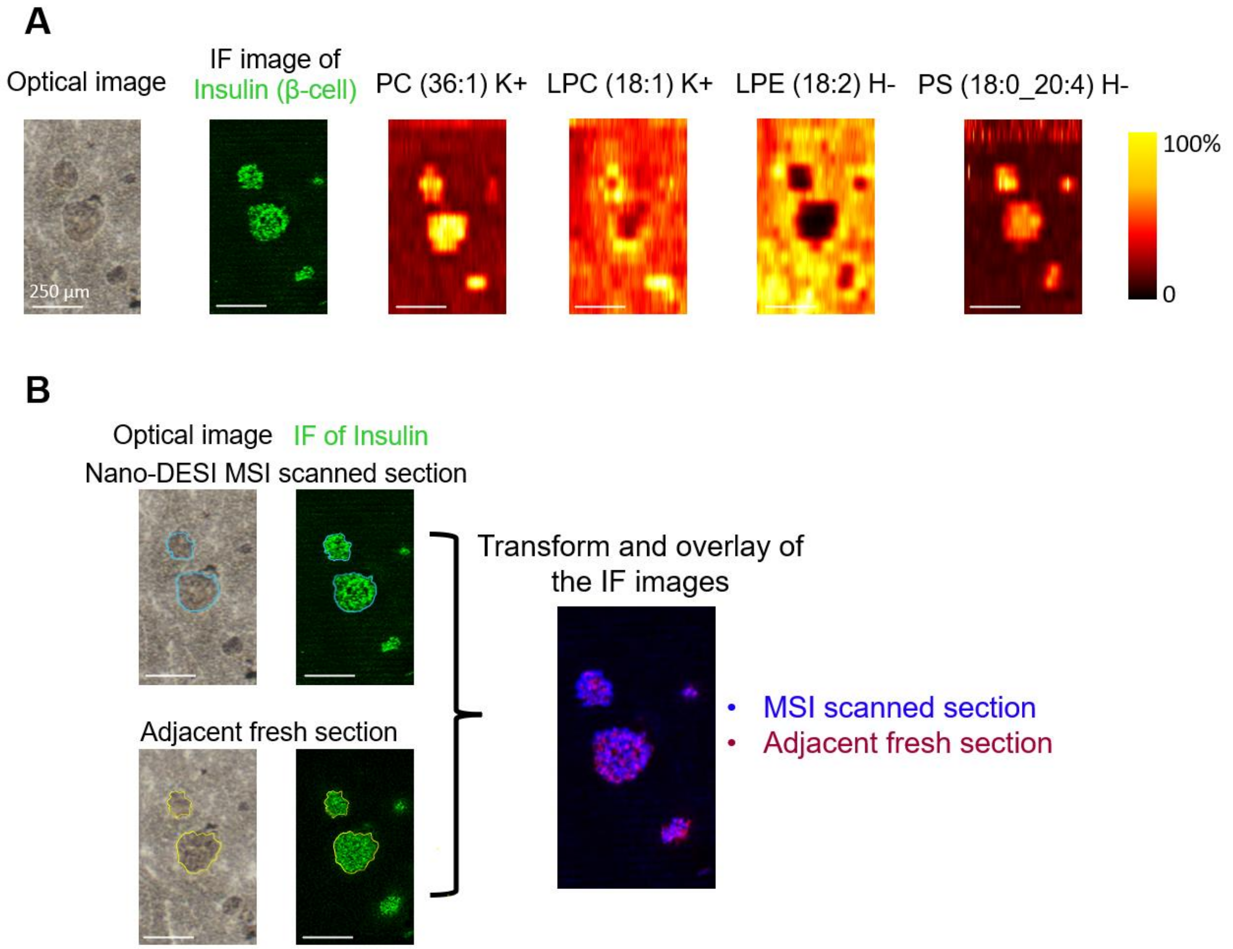


Fig. S15. Validation of protein localization after nano-DESI MSI.

(A) From left to right: brightfield optical image of the pancreatic tissue section (the first image), IF image of insulin obtained from the same section (the second image), example ion images of lipids and metabolites obtained using nano-DESI MSI with MeOH/$H_2O$ as extraction solvent (the third to sixth images). Abbreviations: PC: phosphatidylcholine, LPC: lyso-phosphatidylcholine, LPE: lyso-phosphatidylethanolamine, PS: phosphatidylserine. (B) The brightfield optical images and insulin IF images obtained from the nano-DESI MSI analyzed section (left top) and the adjacent fresh section (left bottom), and overlay of the two insulin IF images (right). Light blue trace delineates the islets outline of the nano-DESI MSI analyzed section and yellow trace delineates the islets outline of the adjacent fresh section. Scale bar: 250 µm.

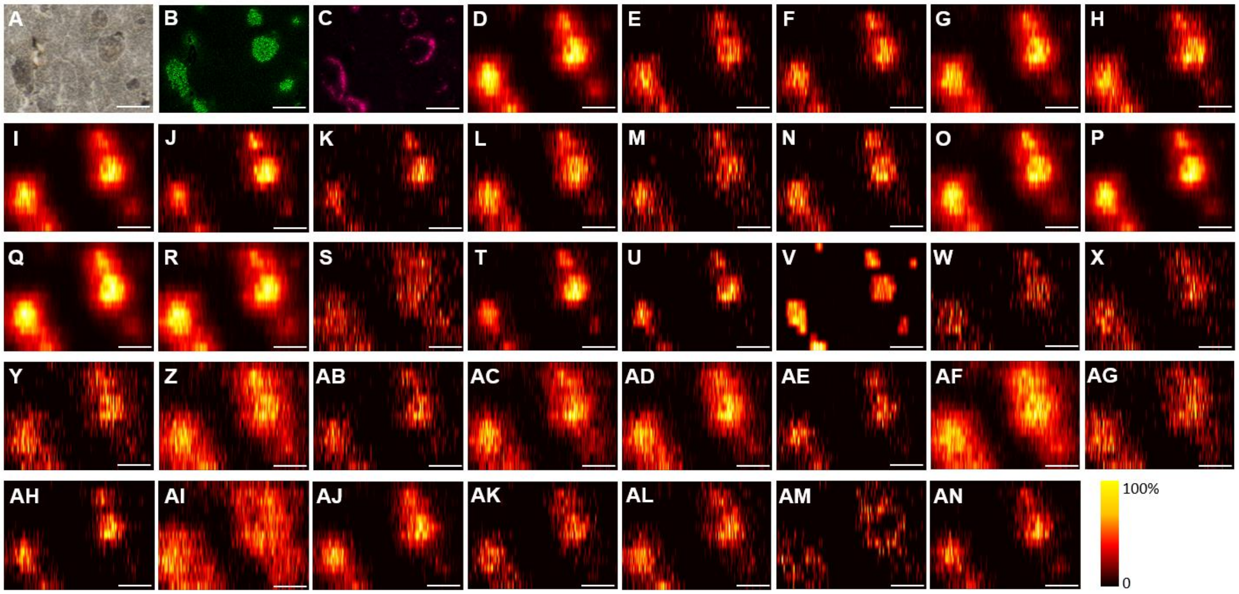


Fig. S16. Ion images of pancreatic peptides obtained from replicate 1.

(A) Brightfield optical image of the analyzed region of the pancreatic tissue section. IF image of (B) INS and (C) glucagon on the adjacent section. Ion images of peptides normalized to TIC: (D) *m/z* 623.1063$^{5-}$, 3121 Da, C-peptide 1, (E) *m/z* 559.0068$^{4-}$, 2240 Da, des-(22-29)-C-peptide 1, (F) *m/z* 469.6200$^{5-}$, 2353 Da, des-(23-29)-C-peptide 1, (G) *m/z* 483.8282$^{5-}$, 2424 Da, des-(24-29)-C-peptide 1, (H) *m/z* 633.3064$^{4-}$, 2537 Da, des-(25-29)-C-peptide 1, (I) *m/z* 625.5064$^{5-}$, 3133 Da, C-peptide 2, (J) *m/z* 374.5096$^{3-}$, 1126 Da, des-(11-31)-C-peptide 2, (K) *m/z* 341.1621$^{4-}$, 1369 Da, des-(13-31)-C-peptide 2, (L) *m/z* 606.2909$^{3-}$, 1822 Da, des-(20-31)-C-peptide 2, (M) *m/z* 511.4924$^{4-}$, 2050 Da, des-(22-31)-C-peptide 2, (N) *m/z* 543.7549$^{4-}$, 2179 Da, des-(23-31)-C-peptide 2, (O) *m/z* 569.0170$^{4-}$, 2280 Da, des-(24-31)-C-peptide 2, (P) *m/z* 597.2876$^{4-}$, 2393 Da, des-(25-31)-C-peptide 2, (Q) *m/z* 615.0473$^{4-}$, 2464 Da, des-(26-31)-C-peptide 2, (R) *m/z* 643.3176$^{4-}$, 2577 Da, des-(27-31)-C-peptide 2, (S) *m/z* 599.6972$^{5-}$, 3004 Da, des-(31)-C-peptide 2, (T) *m/z* 640.8114$^{2-}$, 1284 Da, (U) *m/z* 676.3299$^{2-}$, 1355 Da, (V) *m/z* 720.8988$^{2-}$, 1444 Da, (W) *m/z* 436.7023$^{4-}$, 1751 Da, (X) *m/z* 639.6412$^{3-}$, 1922 Da, (Y) *m/z* 526.7459$^{4-}$, 2111 Da, (Z) *m/z* 536.7561$^{4-}$, 2151 Da, (AB) *m/z* 555.0167$^{4-}$, 2224 Da, (AC) *m/z* 565.0266$^{4-}$, 2264 Da, (AD) *m/z* 572.7758$^{4-}$, 2295 Da, (AE) *m/z* 574.5125$^{4-}$, 2302 Da, (AF) *m/z* 582.7870$^{4-}$, 2335 Da, (AG) *m/z* 601.0472$^{4-}$, 2408 Da, (AH) *m/z* 602.7839$^{4-}$, 2415 Da, (AI) *m/z* 611.0578$^{4-}$, 2448 Da, (AJ) *m/z* 620.5431$^{4-}$, 2486 Da, (AK) *m/z* 624.5357$^{4-}$, 2502 Da, (AL) *m/z* 648.8137$^{4-}$, 2599 Da, (AM) *m/z* 750.8808$^{4-}$, 3007 Da, (AN) *m/z* 629.9023$^{5-}$, 3154 Da. Scale bar: 250 µm.

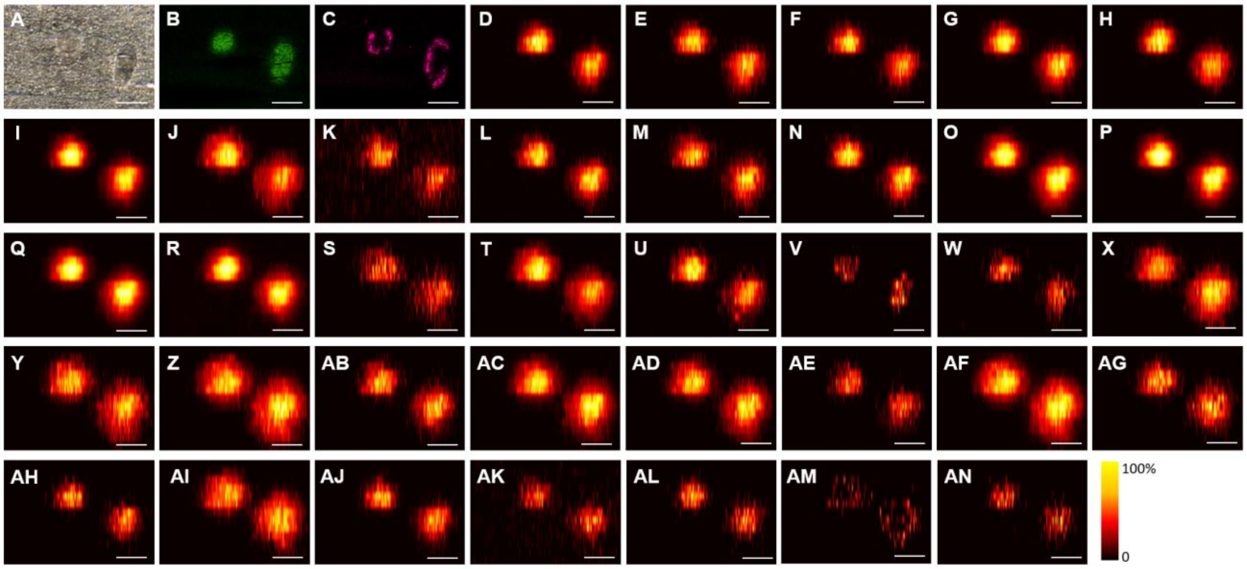


Fig. S17. Ion images of pancreatic peptides obtained from replicate 2.

(A) Brightfield optical image of the analyzed region of the pancreatic tissue section. IF image of (B) INS and (C) glucagon on the adjacent section. Ion images of peptides normalized to TIC: (D) *m/z* 623.1063$^{5-}$, 3121 Da, C-peptide 1, (E) *m/z* 559.0068$^{4-}$, 2240 Da, des-(22-29)-C-peptide 1, (F) *m/z* 469.6200$^{5-}$, 2353 Da, des-(23-29)-C-peptide 1, (G) *m/z* 483.8282$^{5-}$, 2424 Da, des-(24-29)-C-peptide 1, (H) *m/z* 633.3064$^{4-}$, 2537 Da, des-(25-29)-C-peptide 1, (I) *m/z* 625.5064$^{5-}$, 3133 Da, C-peptide 2, (J) *m/z* 374.5096$^{3-}$, 1126 Da, des-(11-31)-C-peptide 2, (K) *m/z* 341.1621$^{4-}$, 1369 Da, des-(13-31)-C-peptide 2, (L) *m/z* 606.2909$^{3-}$, 1822 Da, des-(20-31)-C-peptide 2, (M) *m/z* 511.4924$^{4-}$, 2050 Da, des-(22-31)-C-peptide 2, (N) *m/z* 543.7549$^{4-}$, 2179 Da, des-(23-31)-C-peptide 2, (O) *m/z* 569.0170$^{4-}$, 2280 Da, des-(24-31)-C-peptide 2, (P) *m/z* 597.2876$^{4-}$, 2393 Da, des-(25-31)-C-peptide 2, (Q) *m/z* 615.0473$^{4-}$, 2464 Da, des-(26-31)-C-peptide 2, (R) *m/z* 643.3176$^{4-}$, 2577 Da, des-(27-31)-C-peptide 2, (S) *m/z* 599.6972$^{5-}$, 3004 Da, des-(31)-C-peptide 2, (T) *m/z* 640.8114$^{2-}$, 1284 Da, (U) *m/z* 676.3299$^{2-}$, 1355 Da, (V) *m/z* 720.8988$^{2-}$, 1444 Da, (W) *m/z* 436.7023$^{4-}$, 1751 Da, (X) *m/z* 639.6412$^{3-}$, 1922 Da, (Y) *m/z* 526.7459$^{4-}$, 2111 Da, (Z) *m/z* 536.7561$^{4-}$, 2151 Da, (AB) *m/z* 555.0167$^{4-}$, 2224 Da, (AC) *m/z* 565.0266$^{4-}$, 2264 Da, (AD) *m/z* 572.7758$^{4-}$, 2295 Da, (AE) *m/z* 574.5125$^{4-}$, 2302 Da, (AF) *m/z* 582.7870$^{4-}$, 2335 Da, (AG) *m/z* 601.0472$^{4-}$, 2408 Da, (AH) *m/z* 602.7839$^{4-}$, 2415 Da, (AI) *m/z* 611.0578$^{4-}$, 2448 Da, (AJ) *m/z* 620.5431$^{4-}$, 2486 Da, (AK) *m/z* 624.5357$^{4-}$, 2502 Da, (AL) *m/z* 648.8137$^{4-}$, 2599 Da, (AM) *m/z* 750.8808$^{4-}$, 3007 Da, (AN) *m/z* 629.9023$^{5-}$, 3154 Da. Scale bar: 250 µm.

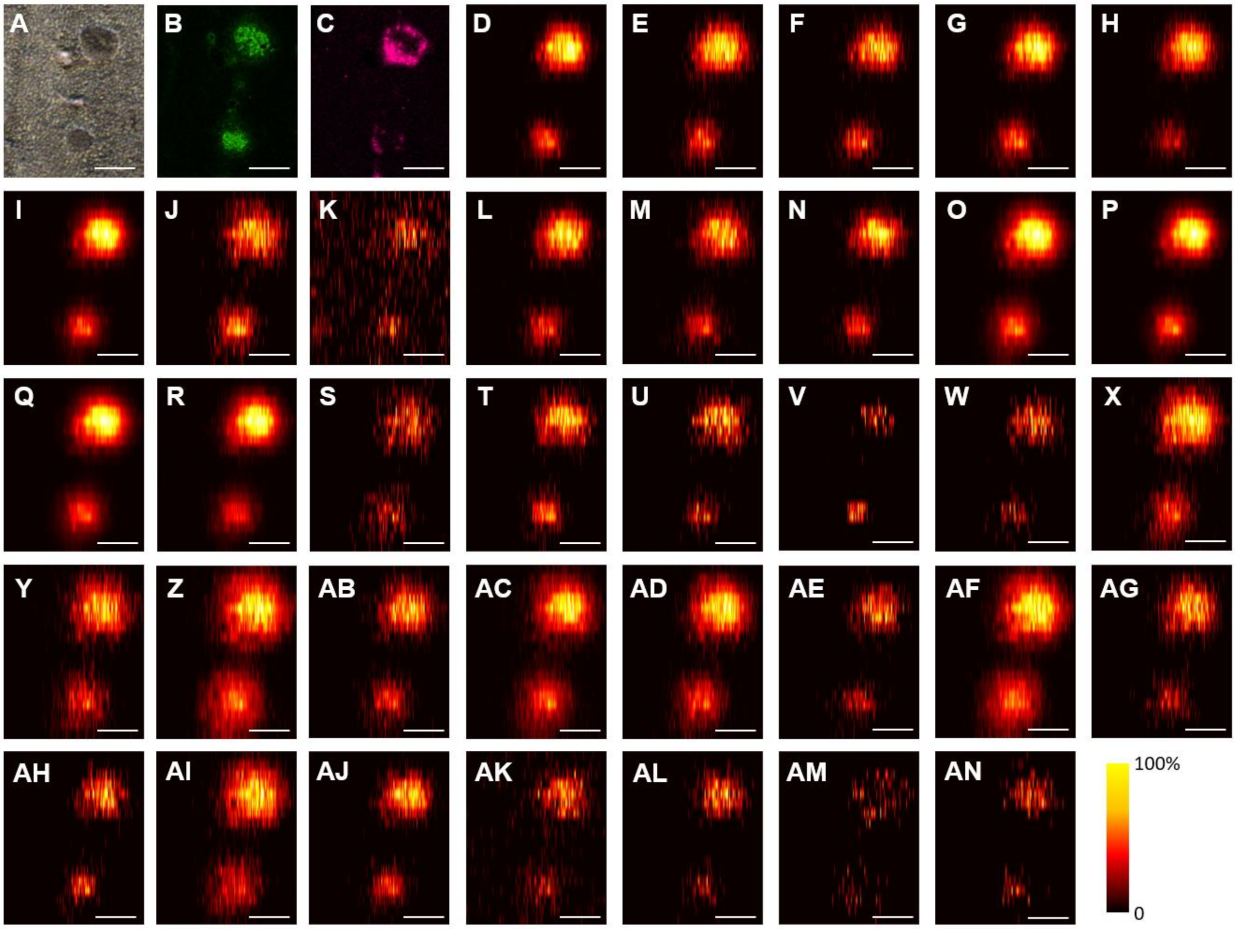


Fig. S18. Ion images of pancreatic peptides obtained from replicate 3.

(A) Brightfield optical image of the analyzed region of the pancreatic tissue section. IF image of (B) INS and (C) glucagon on the adjacent section. Ion images of peptides normalized to TIC: (D) *m/z* 623.1063$^{5-}$, 3121 Da, C-peptide 1, (E) *m/z* 559.0068$^{4-}$, 2240 Da, des-(22-29)-C-peptide 1, (F) *m/z* 469.6200$^{5-}$, 2353 Da, des-(23-29)-C-peptide 1, (G) *m/z* 483.8282$^{5-}$, 2424 Da, des-(24-29)-C-peptide 1, (H) *m/z* 633.3064$^{4-}$, 2537 Da, des-(25-29)-C-peptide 1, (I) *m/z* 625.5064$^{5-}$, 3133 Da, C-peptide 2, (J) *m/z* 374.5096$^{3-}$, 1126 Da, des-(11-31)-C-peptide 2, (K) *m/z* 341.1621$^{4-}$, 1369 Da, des-(13-31)-C-peptide 2, (L) *m/z* 606.2909$^{3-}$, 1822 Da, des-(20-31)-C-peptide 2, (M) *m/z* 511.4924$^{4-}$, 2050 Da, des-(22-31)-C-peptide 2, (N) *m/z* 543.7549$^{4-}$, 2179 Da, des-(23-31)-C-peptide 2, (O) *m/z* 569.0170$^{4-}$, 2280 Da, des-(24-31)-C-peptide 2, (P) *m/z* 597.2876$^{4-}$, 2393 Da, des-(25-31)-C-peptide 2, (Q) *m/z* 615.0473$^{4-}$, 2464 Da, des-(26-31)-C-peptide 2, (R) *m/z* 643.3176$^{4-}$, 2577 Da, des-(27-31)-C-peptide 2, (S) *m/z* 599.6972$^{5-}$, 3004 Da, des-(31)-C-peptide 2, (T) *m/z* 640.8114$^{2-}$, 1284 Da, (U) *m/z* 676.3299$^{2-}$, 1355 Da, (V) *m/z* 720.8988$^{2-}$, 1444 Da, (W) *m/z* 436.7023$^{4-}$, 1751 Da, (X) *m/z* 639.6412$^{3-}$, 1922 Da, (Y) *m/z* 526.7459$^{4-}$, 2111 Da, (Z) *m/z* 536.7561$^{4-}$, 2151 Da, (AB) *m/z* 555.0167$^{4-}$, 2224 Da, (AC) *m/z* 565.0266$^{4-}$, 2264 Da, (AD) *m/z* 572.7758$^{4-}$, 2295 Da, (AE) *m/z* 574.5125$^{4-}$, 2302 Da, (AF) *m/z* 582.7870$^{4-}$, 2335 Da, (AG) *m/z* 601.0472$^{4-}$, 2408 Da, (AH) *m/z* 602.7839$^{4-}$, 2415 Da, (AI) *m/z* 611.0578$^{4-}$, 2448 Da, (AJ) *m/z* 620.5431$^{4-}$, 2486 Da, (AK) *m/z* 624.5357$^{4-}$, 2502 Da, (AL) *m/z* 648.8137$^{4-}$, 2599 Da, (AM) *m/z* 750.8808$^{4-}$, 3007 Da, (AN) *m/z* 629.9023$^{5-}$, 3154 Da. Scale bar: 250 µm.

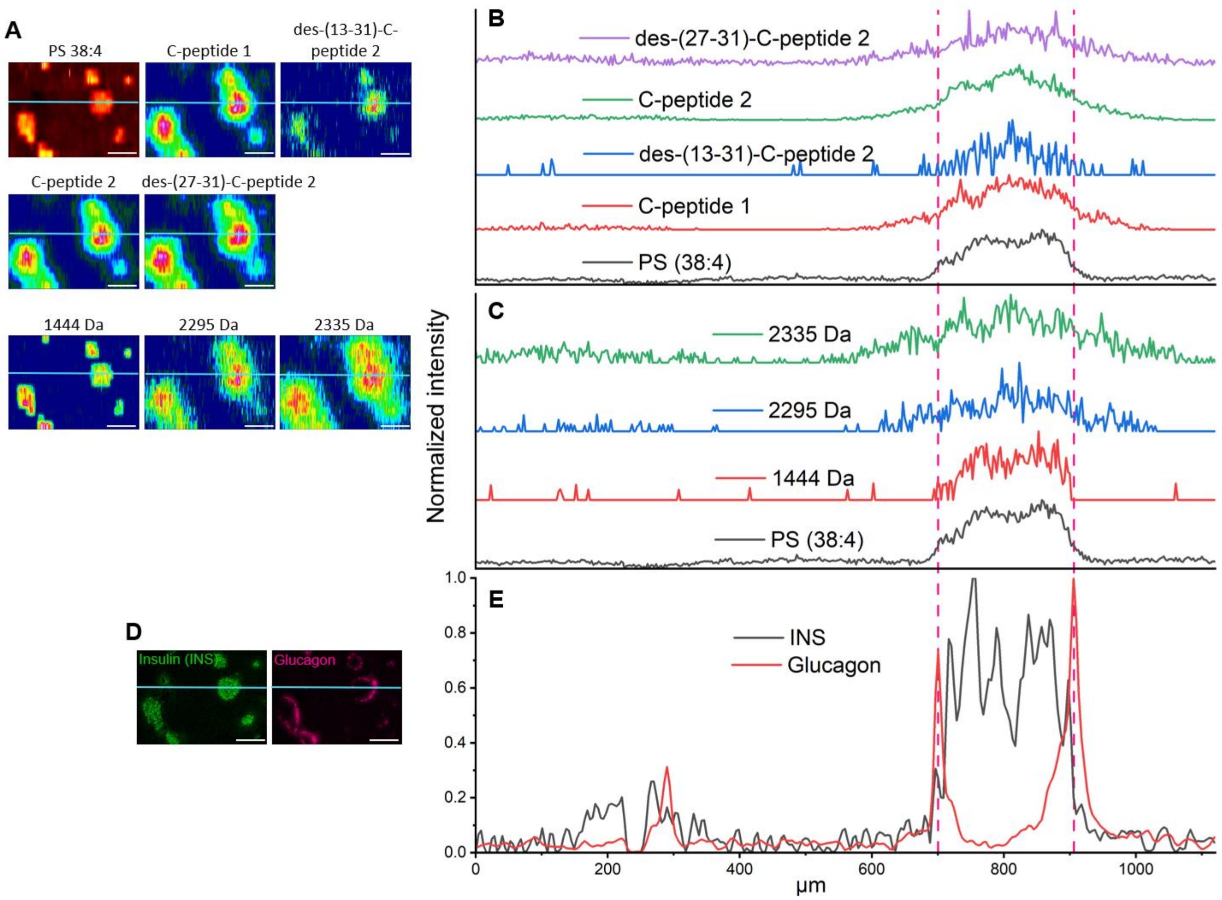


Fig. S19. Extracted line profiles of deprotonated peptides from ion images and extracted line profiles of insulin and glucagon from IF images.

(A) The ion images of peptides with the light blue trace marking the location of the extracted line profiles. Overlay of the line profiles of (B) C-peptides and (C) other peptides. (D) The insulin and glucagon IF images with the light blue trace marking the location of (E) the extracted line profiles. Scale bar: 250 μm. The signal intensities were self-normalized.

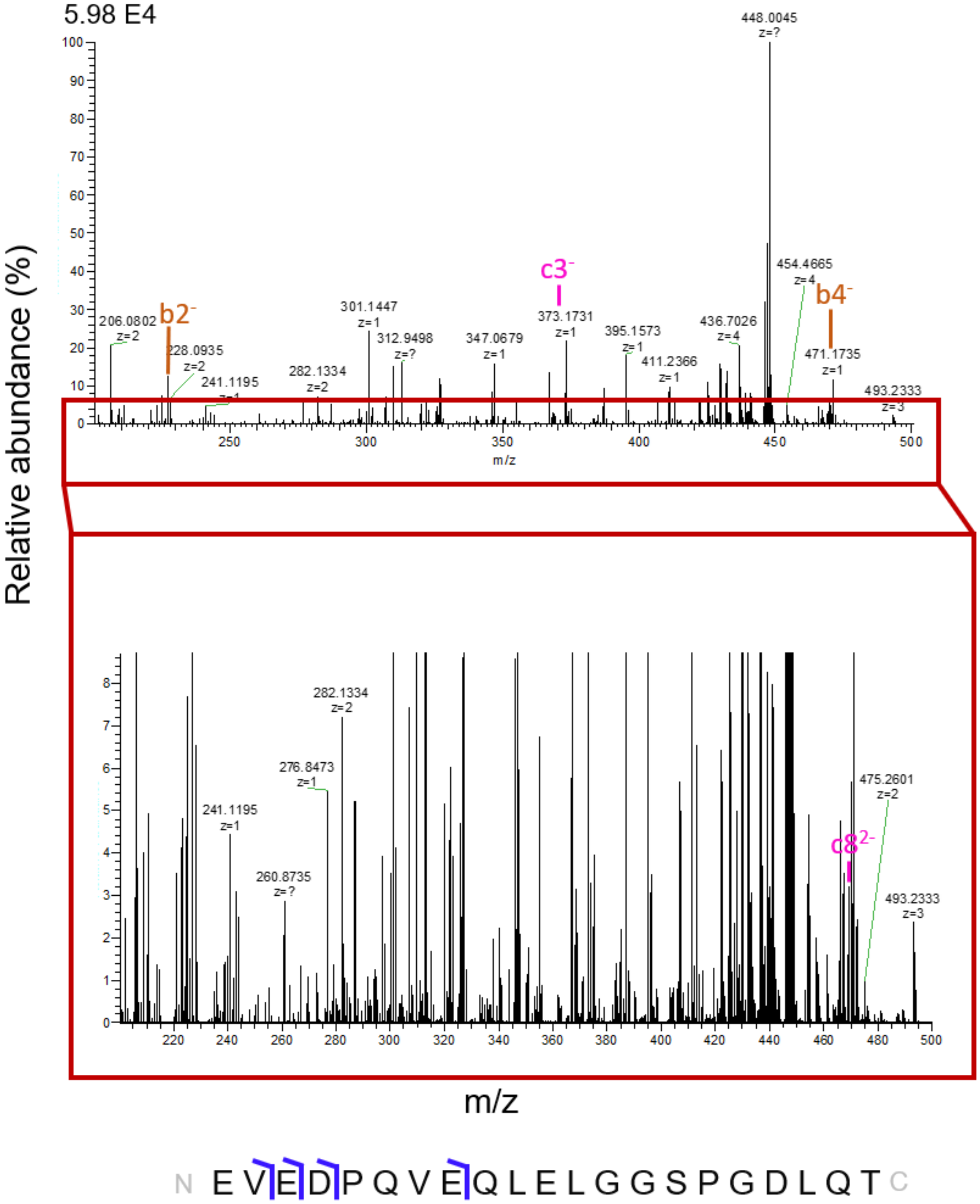


Fig. S20. MS/MS spectrum (top), zoomed-in MS/MS spectrum (middle) and the fragmentation map (bottom) of the -5 charge state of the 2240 Da des-(22-29)-C-peptide-1 at m/z 447.0035.

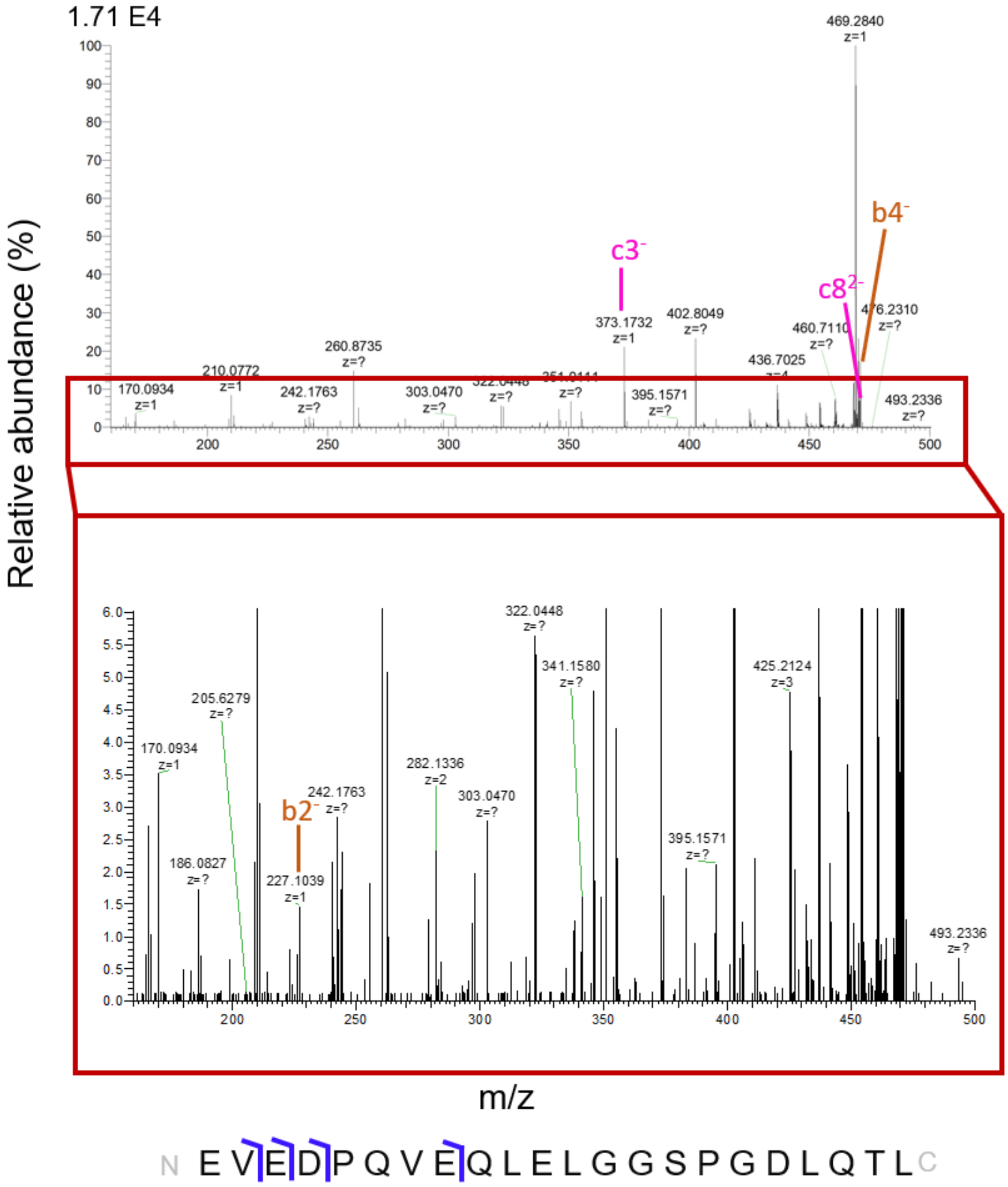


Fig. S21. MS/MS spectrum (top), zoomed-in MS/MS spectrum (middle) and the fragmentation map (bottom) of the -5 charge state of 2353 Da des-(23-29)-C-peptide-1 at m/z 469.62.

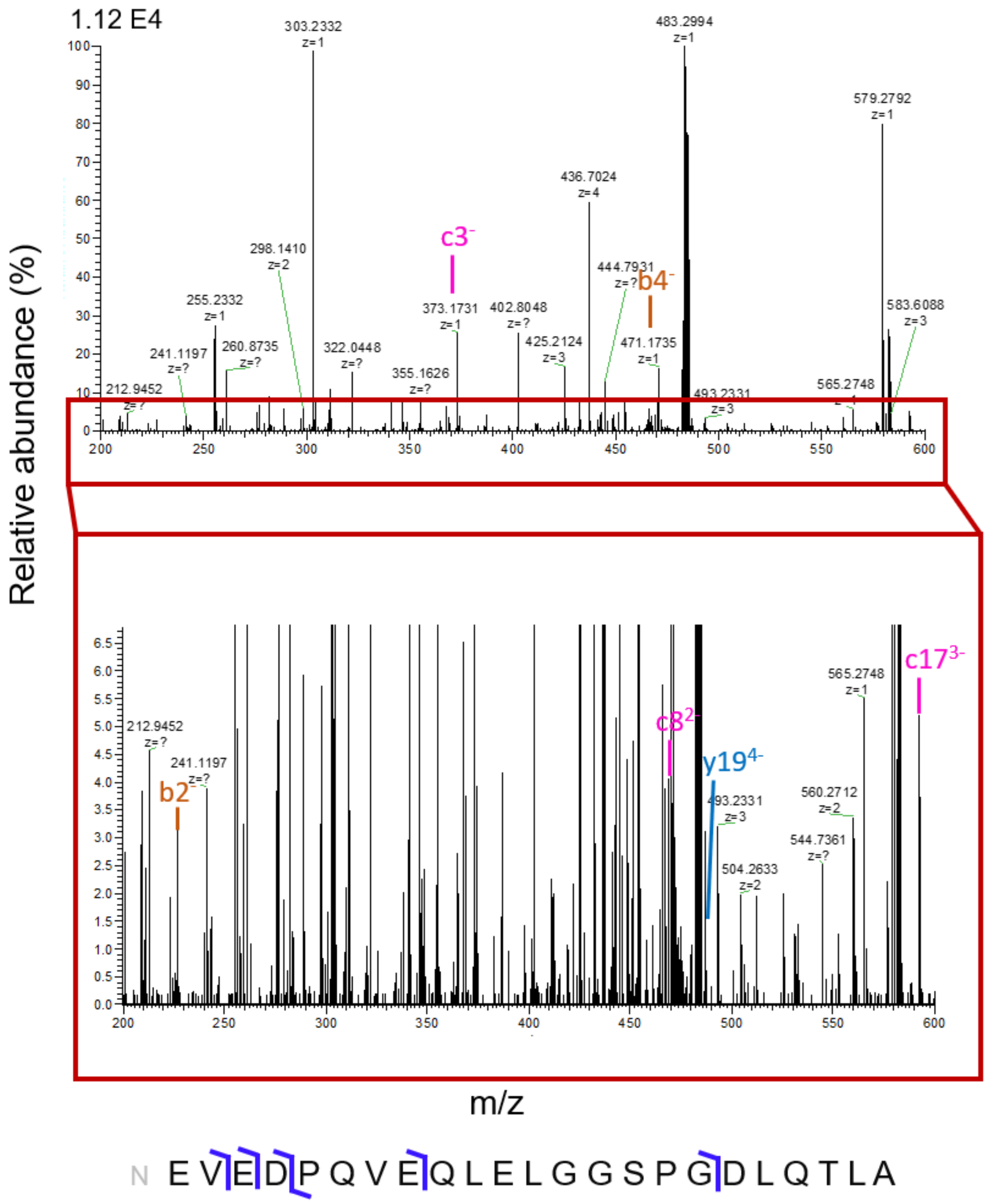


Fig. S22. MS/MS spectrum (top), zoomed-in MS/MS spectrum (middle) and the fragmentation map (bottom) of the -5 charge state of 2424 Da des-(24-29)-C-peptide-1 at m/z 483.8278.

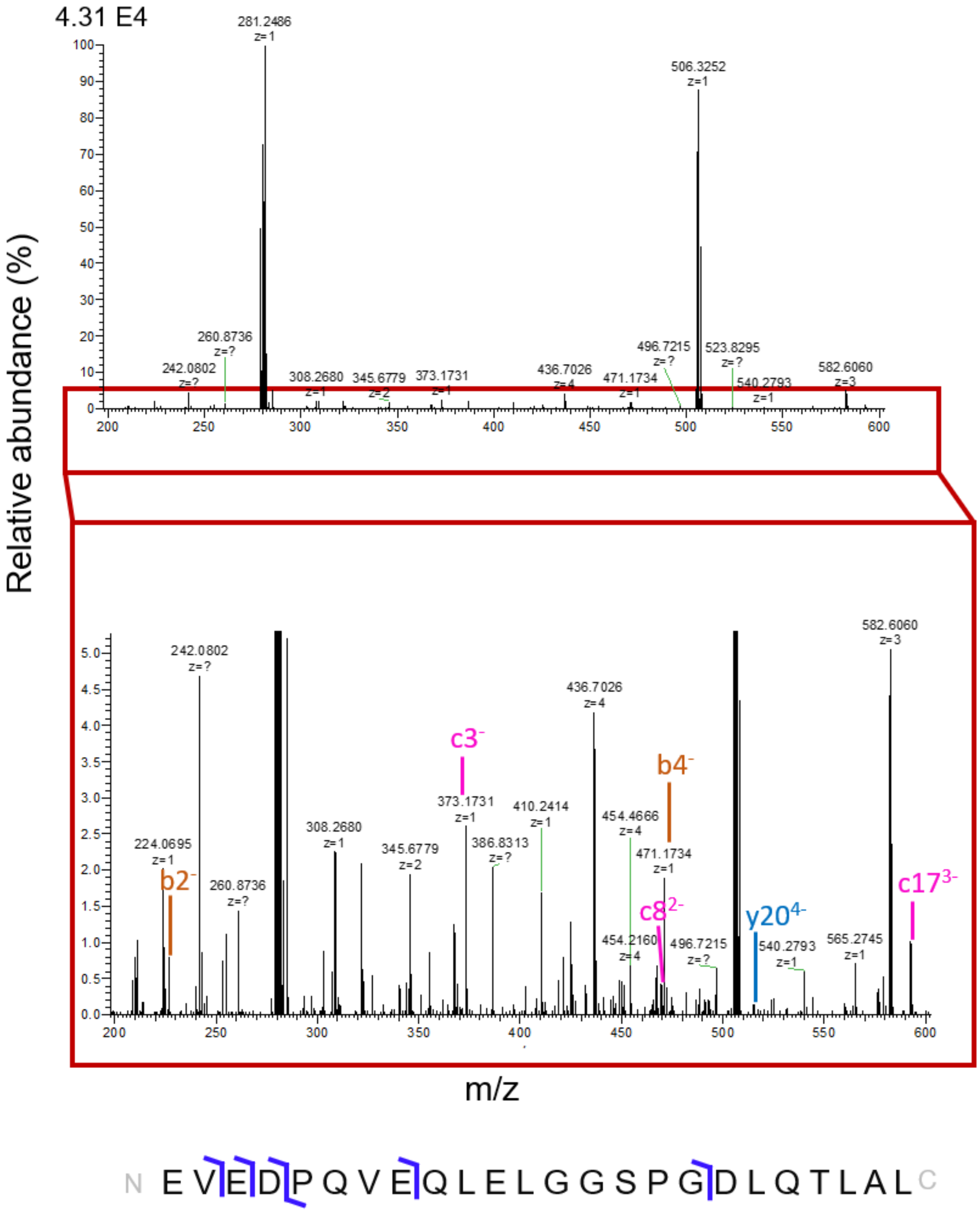


Fig. S23. MS/MS spectrum (top), zoomed-in MS/MS spectrum (middle) and the fragmentation map (bottom) of the -5 charge state of 2537 Da des-(25-29)-C-peptide-1 at m/z 506.4447.

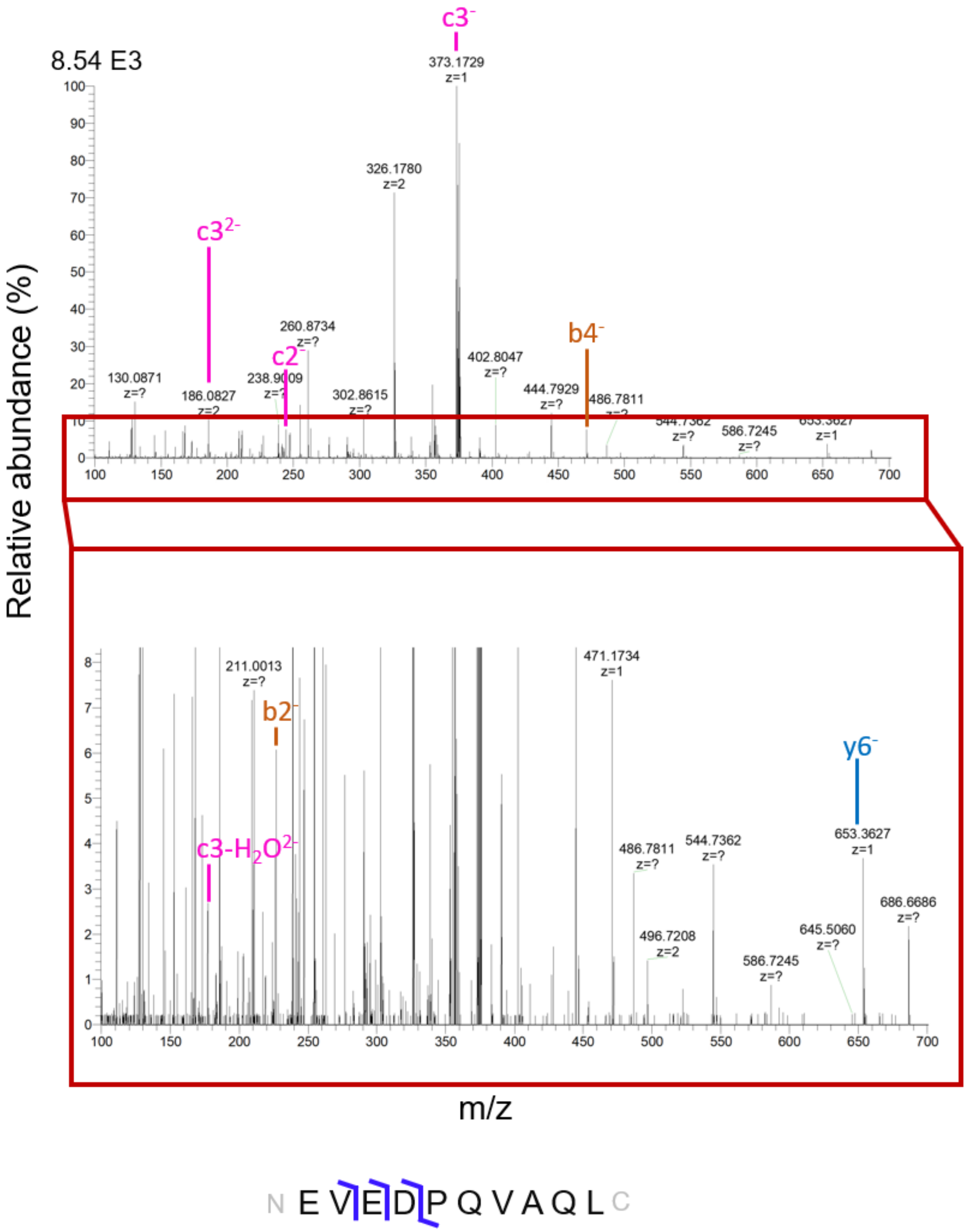


Fig. S24. MS/MS spectrum (top), zoomed-in MS/MS spectrum (middle) and the fragmentation map (bottom) of the -3 charge state of the 1126 Da des-(11-31)-C-peptide-2 at m/z 374.5096.

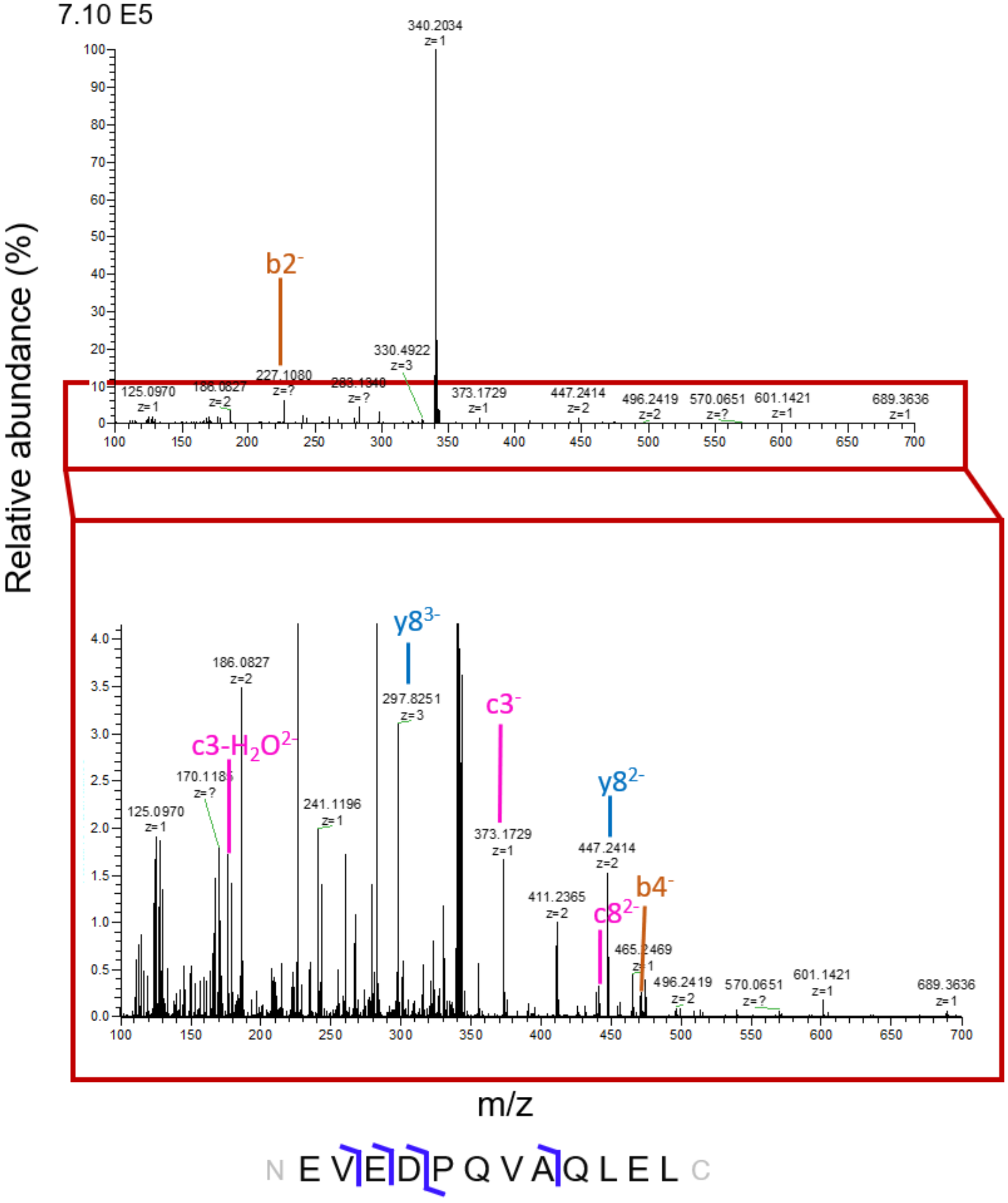


Fig. S25. MS/MS spectrum (top), zoomed-in MS/MS spectrum (middle) and the fragmentation map (bottom) of the -4 charge state of the 1368 Da des-(13-31)-C-peptide-2 at m/z 341.161.

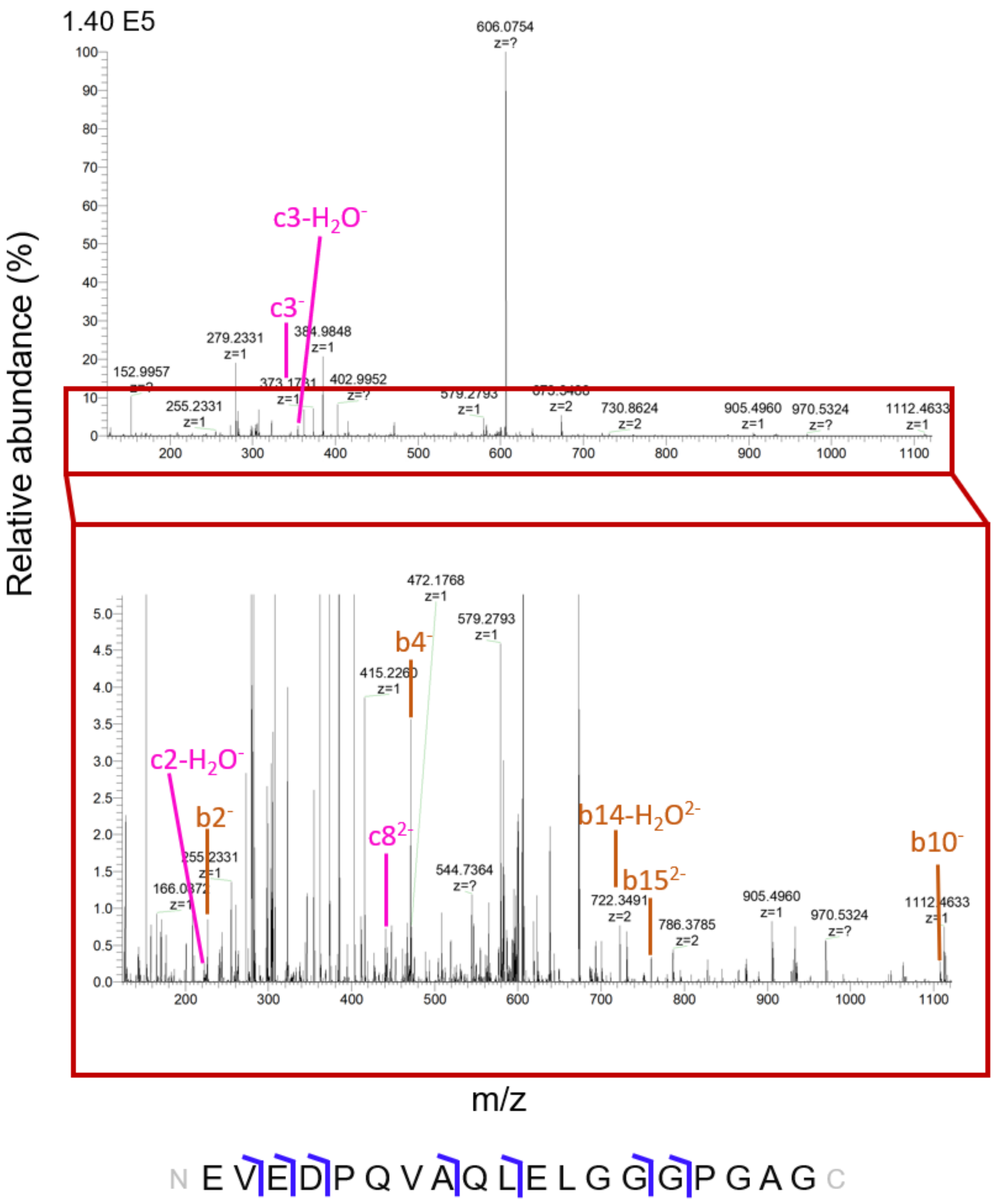


Fig. S26. MS/MS spectrum (top), zoomed-in MS/MS spectrum (middle) and the fragmentation map (bottom) of the -3 charge state of the 1821 Da des-(20-31)-C-peptide-2 at m/z 606.291.

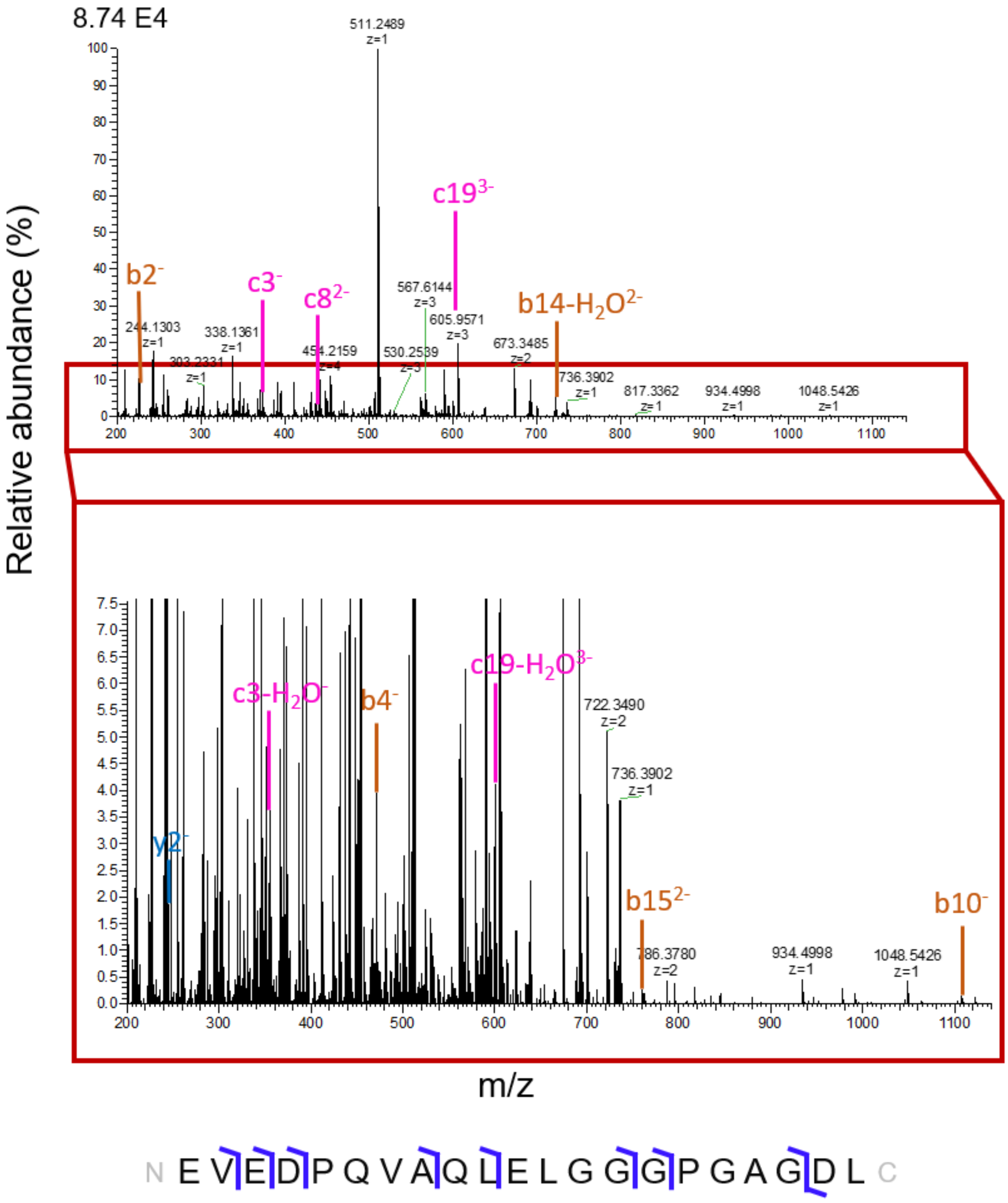


Fig. S27. MS/MS spectrum (top), zoomed-in MS/MS spectrum (middle) and the fragmentation map (bottom) of the -4 charge state of the 2050 Da des-(22-31)-C-peptide-2 at m/z 511.4920.

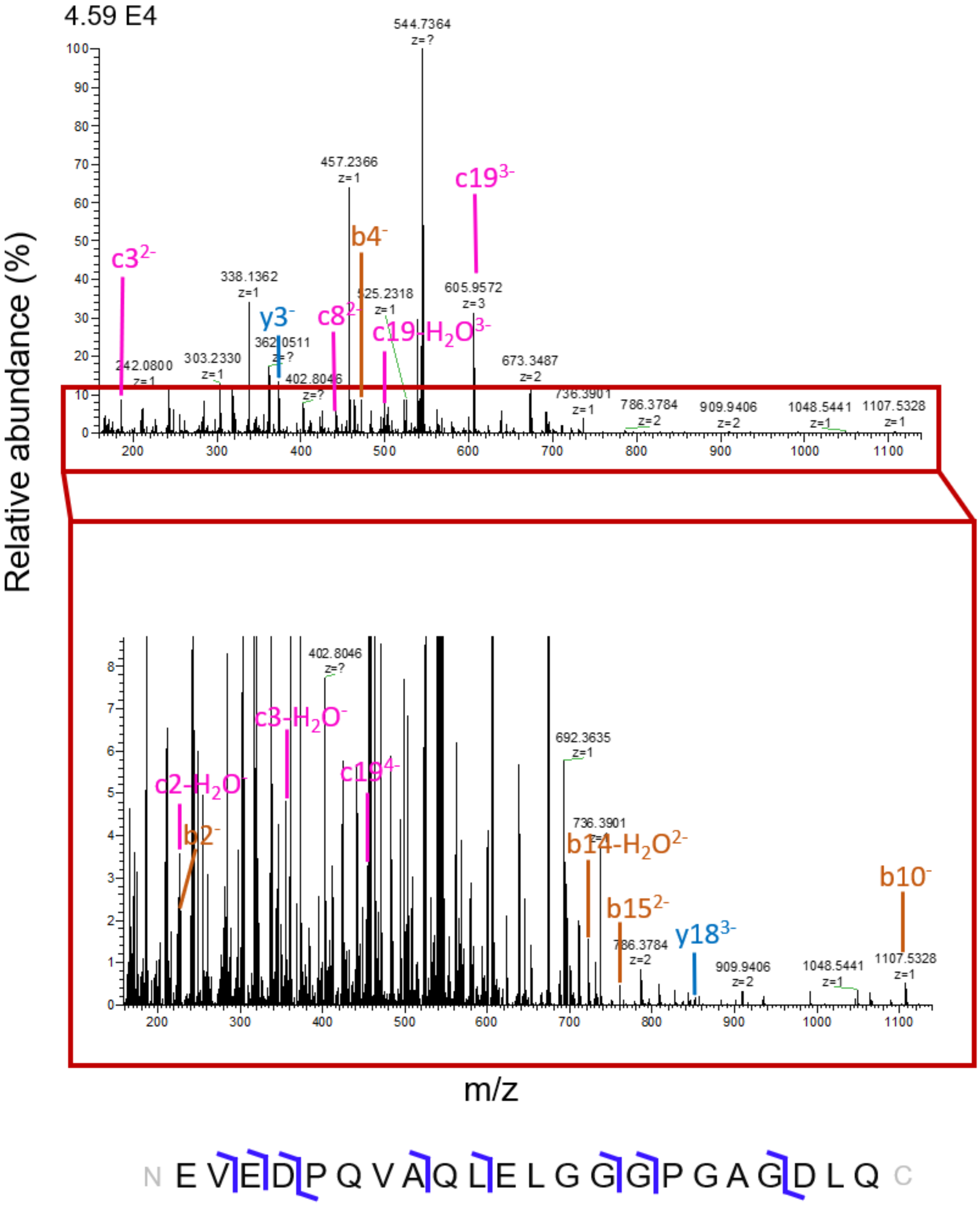


Fig. S28. MS/MS spectrum (top), zoomed-in MS/MS spectrum (middle) and the fragmentation map (bottom) of the -4 charge state of the 2179 Da des-(23-31)-C-peptide-2 at m/z 543.755.

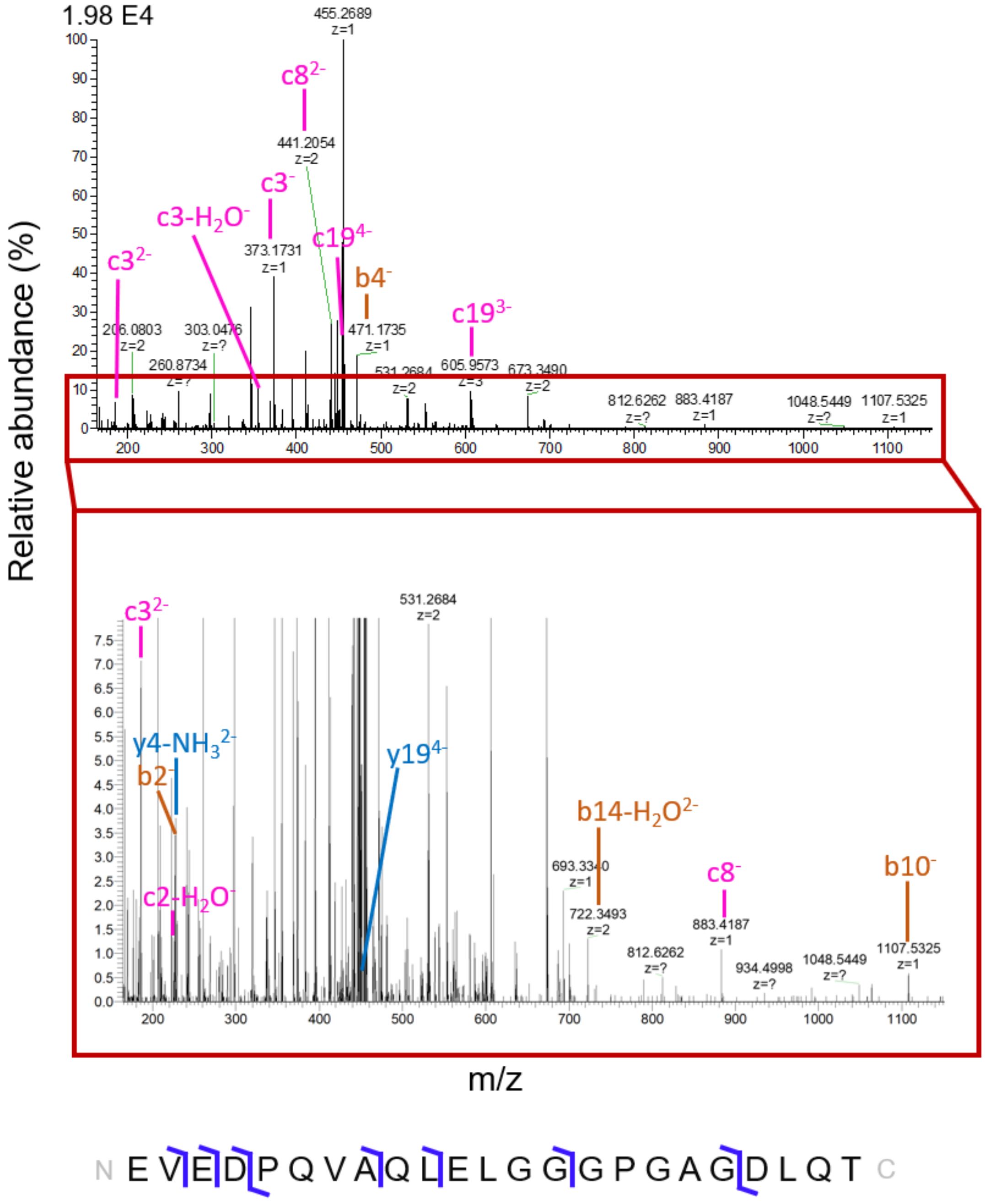


Fig. S29. MS/MS spectrum (top), zoomed-in MS/MS spectrum (middle) and the fragmentation map (bottom) of the -5 charge state of the 2280 Da des-(24-31)-C-peptide-2 at m/z 455.012.

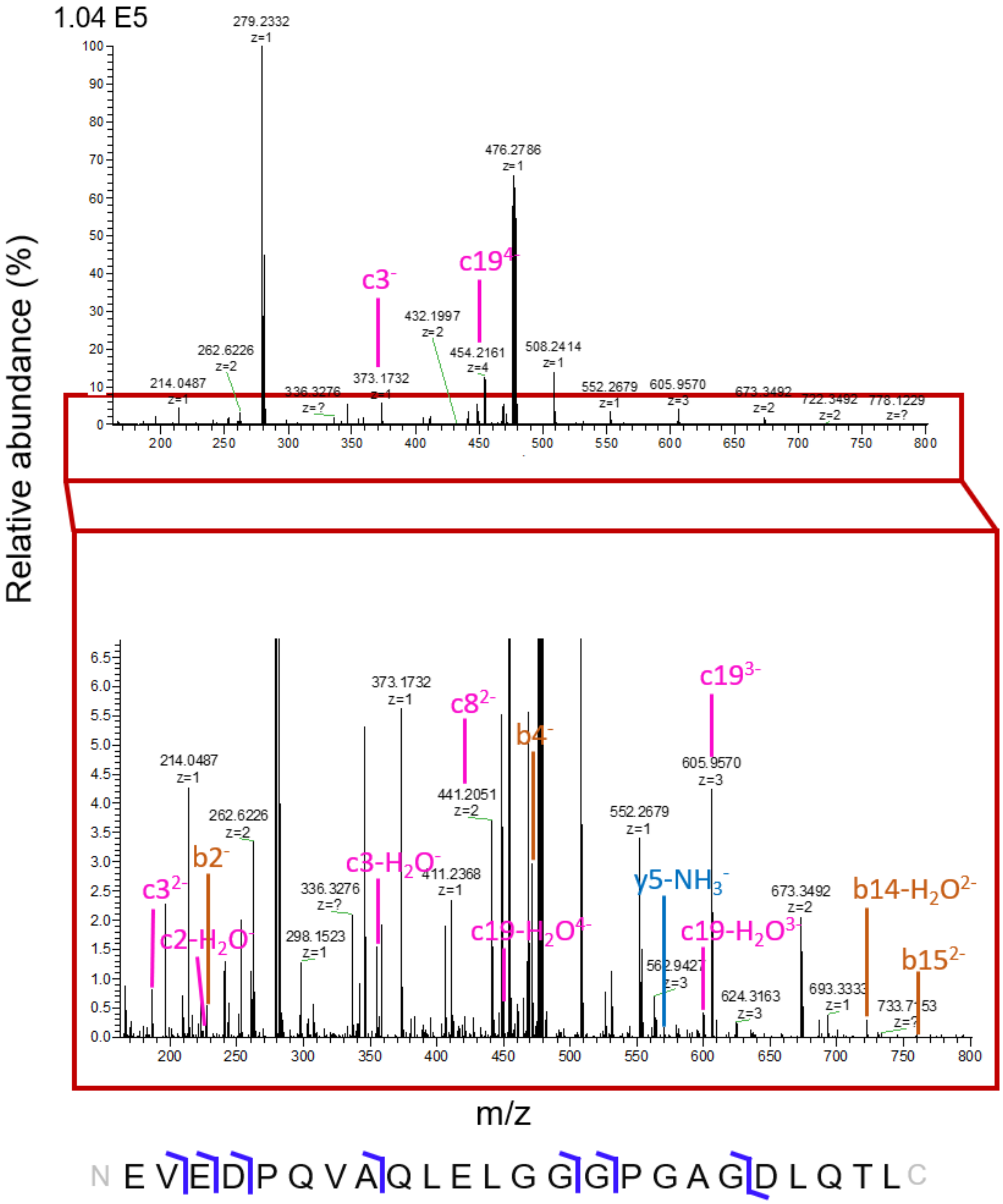


Fig. S30. MS/MS spectrum (top), zoomed-in MS/MS spectrum (middle) and the fragmentation map (bottom) of the -5 charge state of the 2393 Da des-(25-31)-C-peptide-2 at m/z 477.629.

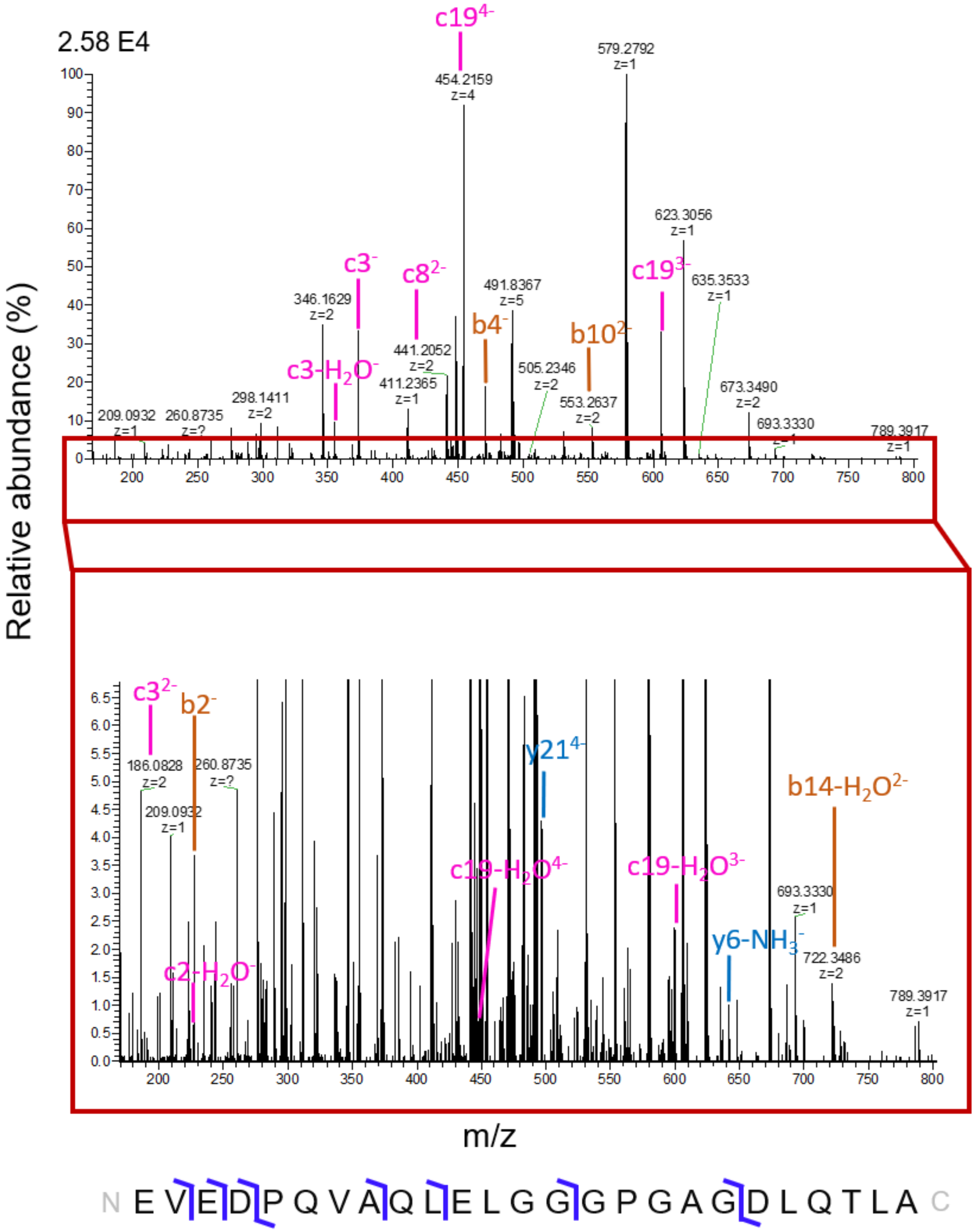


Fig. S31. MS/MS spectrum (top), zoomed-in MS/MS spectrum (middle) and the fragmentation map (bottom) of the -5 charge state of the 2464 Da des-(26-31)-C-peptide-2 at m/z 491.8364.

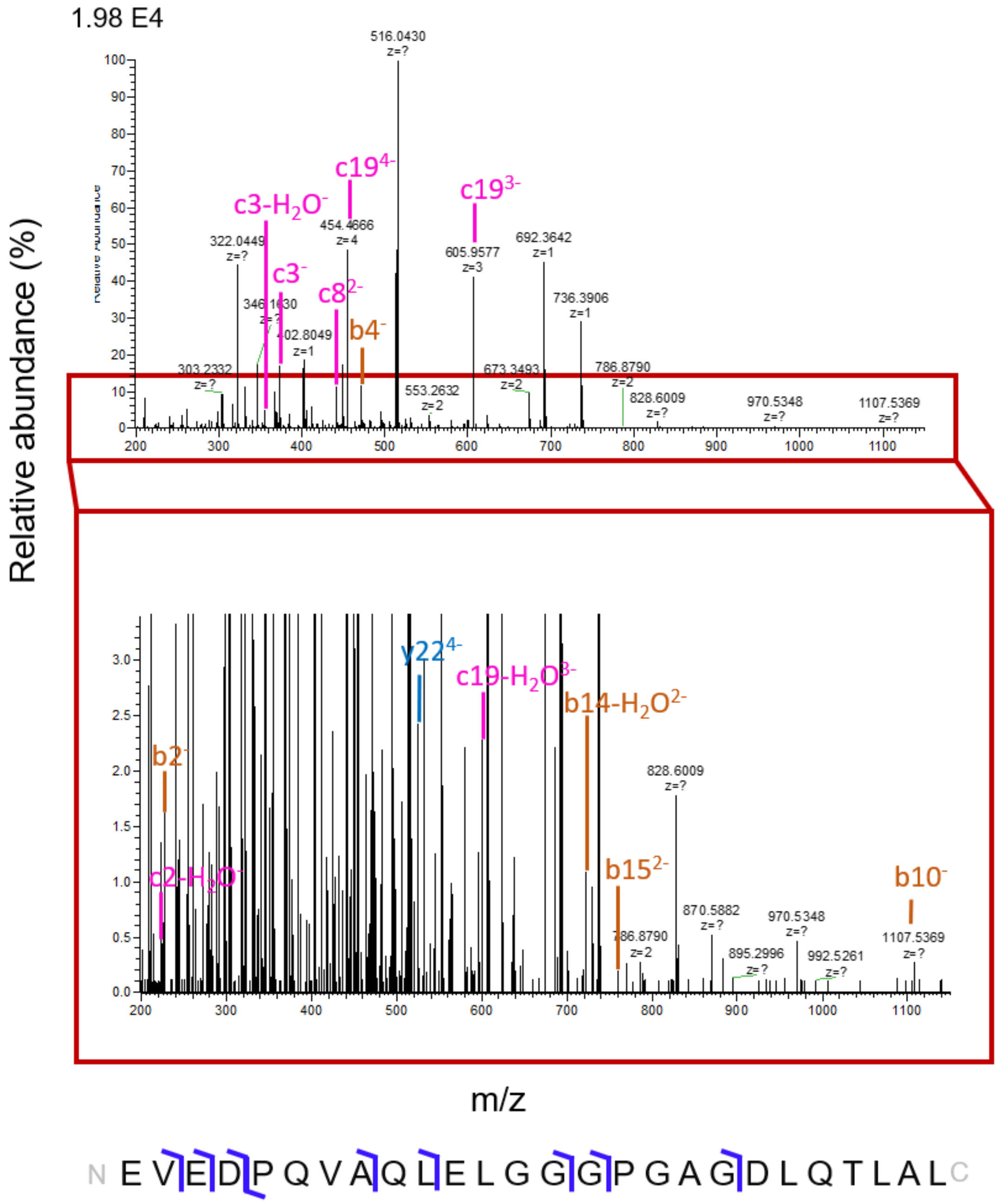


Fig. S32. MS/MS spectrum (top), zoomed-in MS/MS spectrum (middle) and the fragmentation map (bottom) of the -5 charge state of the 2577 Da des-(27-31)-C-peptide-2. at m/z 514.4532.

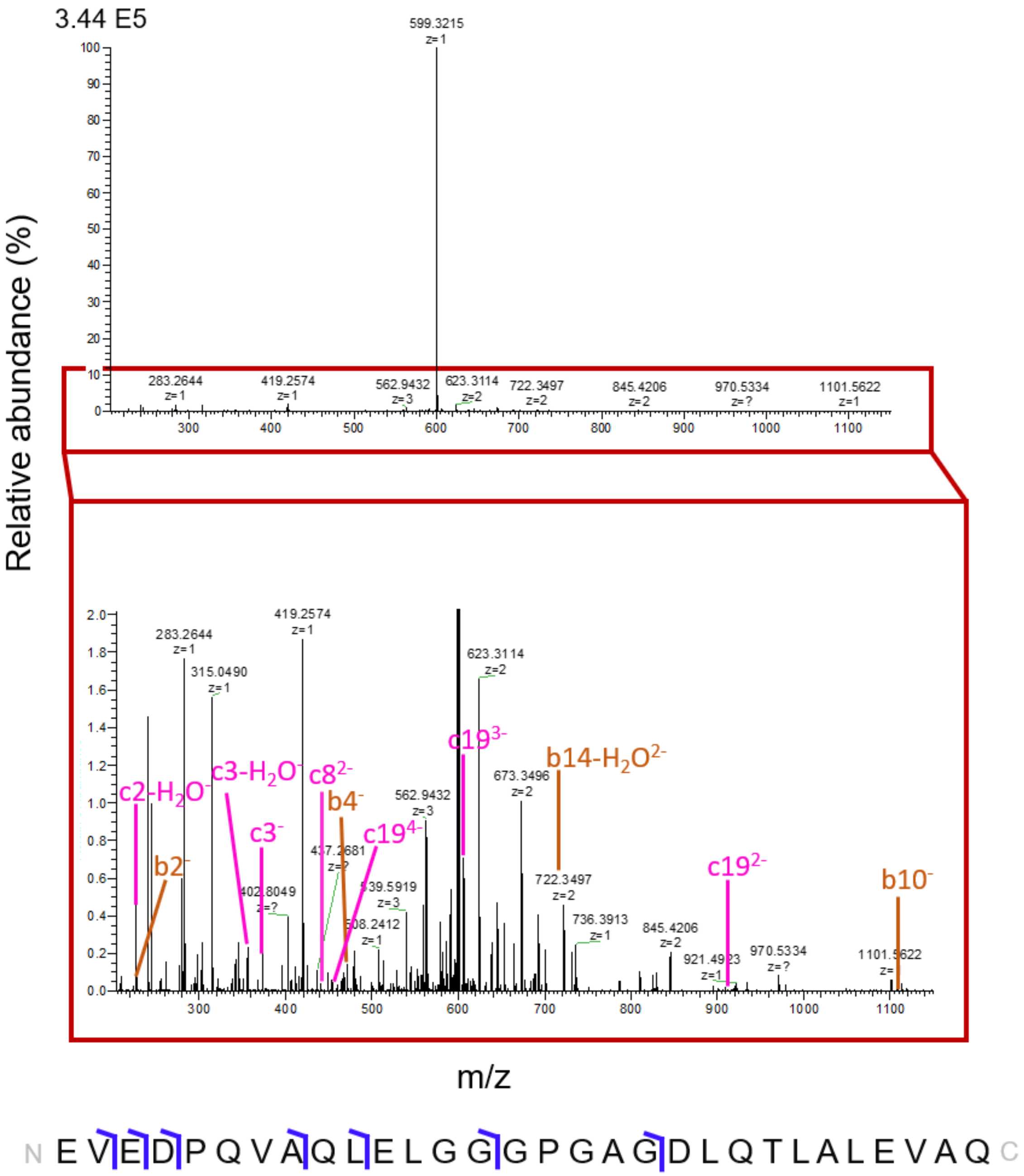


Fig. S33. MS/MS spectrum (top), zoomed-in MS/MS spectrum (middle) and the fragmentation map (bottom) of the -5 charge state of the 3003 Da des-(31)-C-peptide-2 at m/z 599.697.

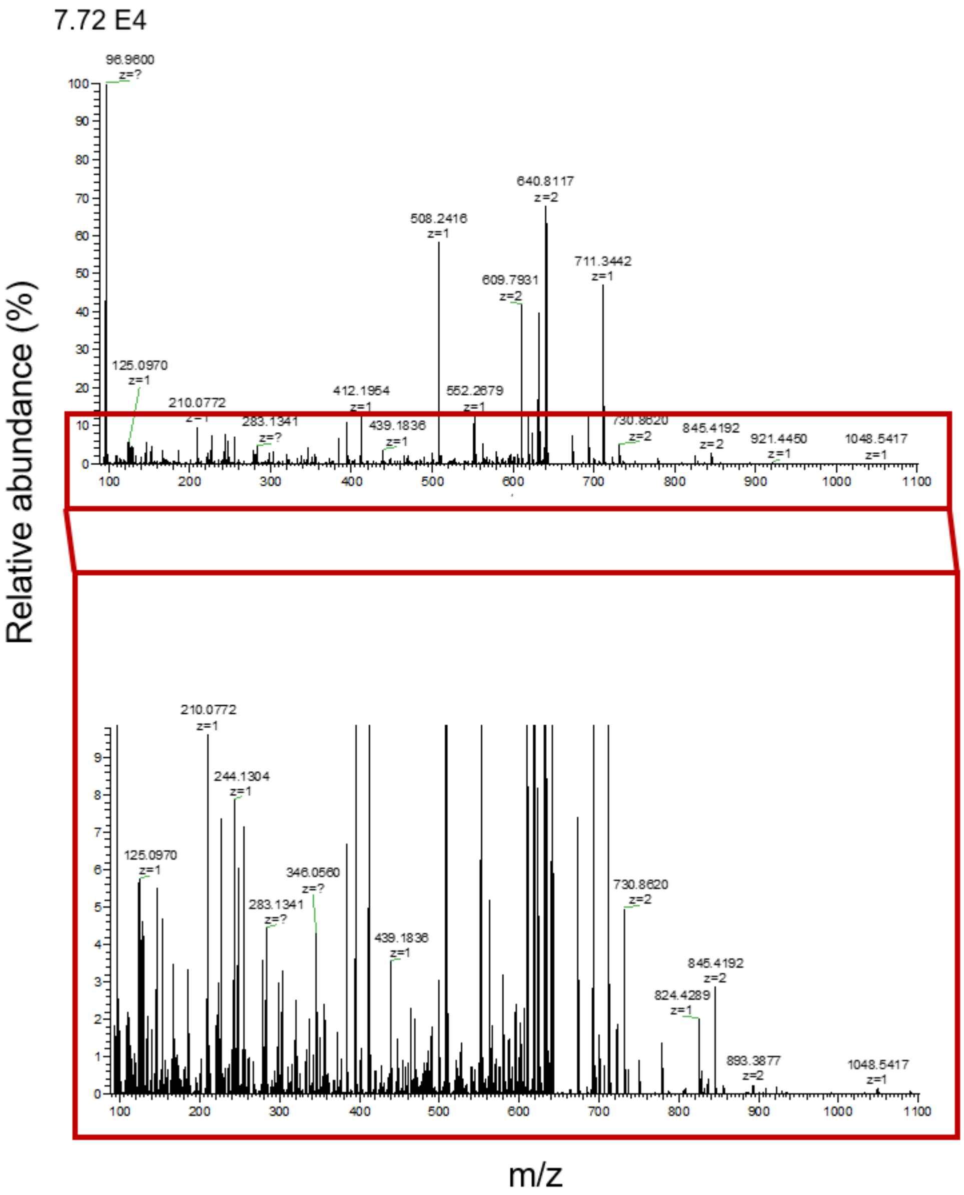


Fig. S34. MS/MS spectrum (top) and zoomed-in MS/MS spectrum (bottom) of the -2 charge state of the 1284 Da at m/z 640.8114.

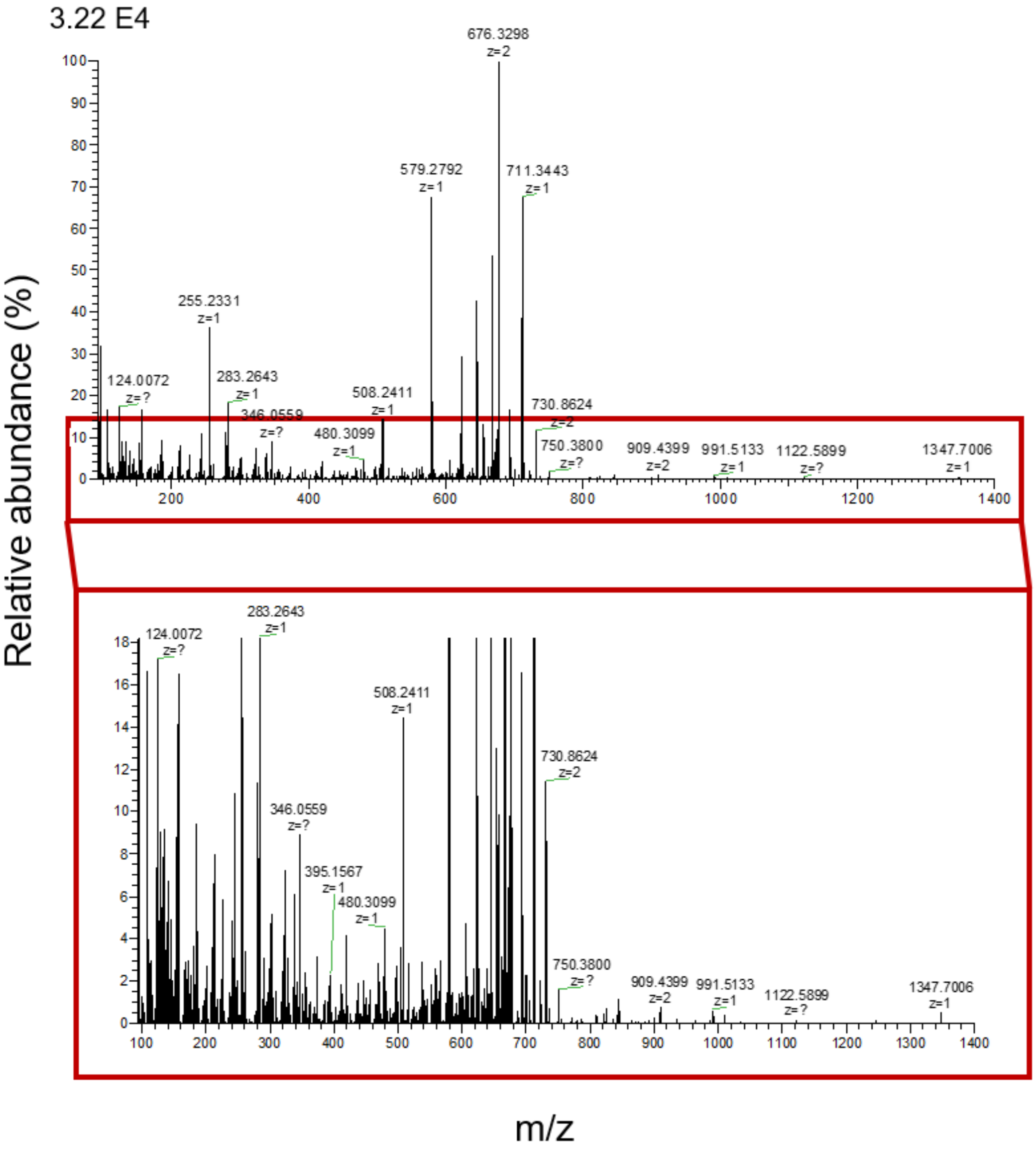


Fig. S35. MS/MS spectrum (top) and zoomed-in MS/MS spectrum (bottom) of the -2 charge state of the 1355 Da at m/z 676.3299.

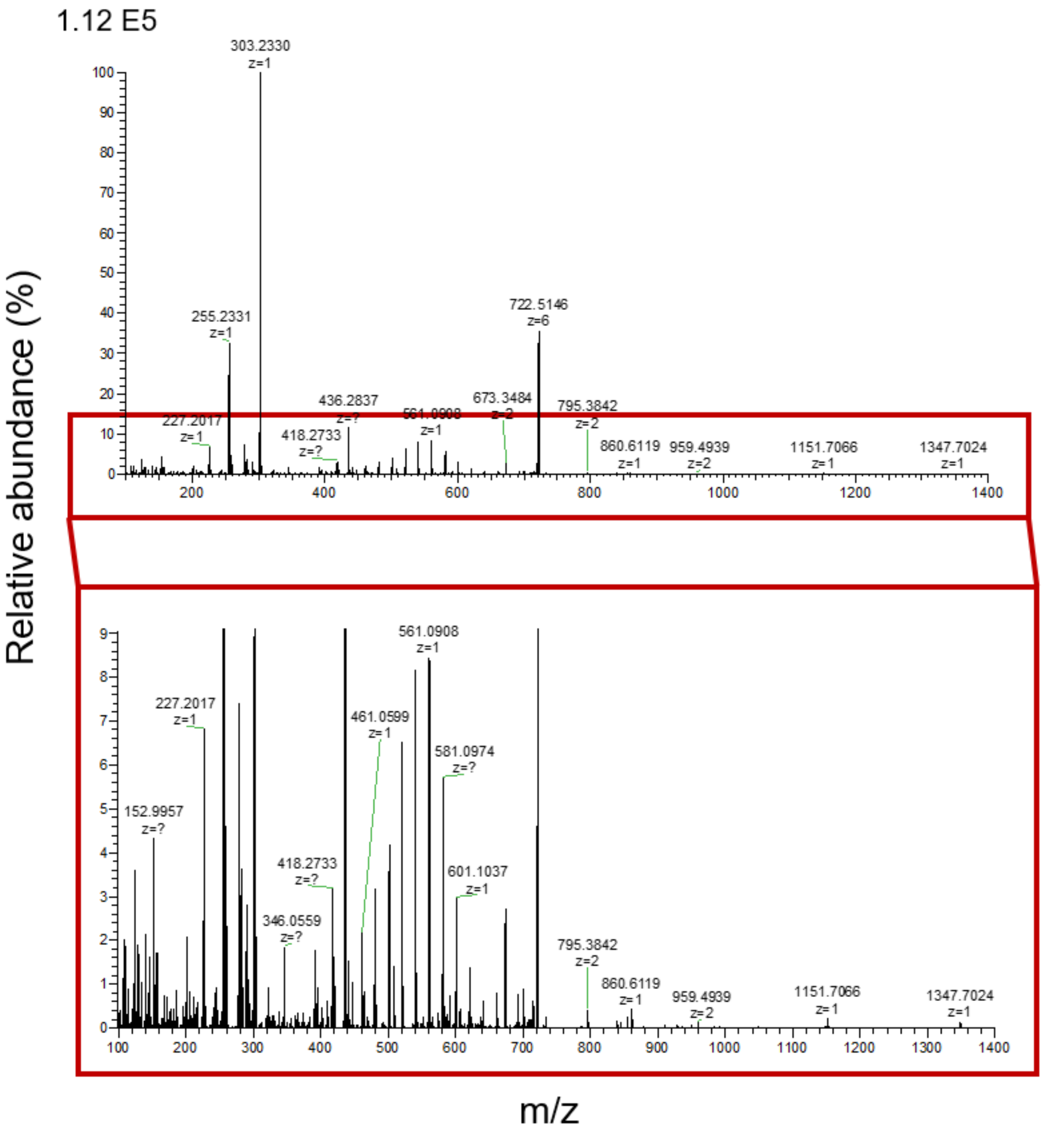


Fig. S36. MS/MS spectrum (top) and zoomed-in MS/MS spectrum (bottom) of the -2 charge state of the 1444 Da at m/z 720.8988.

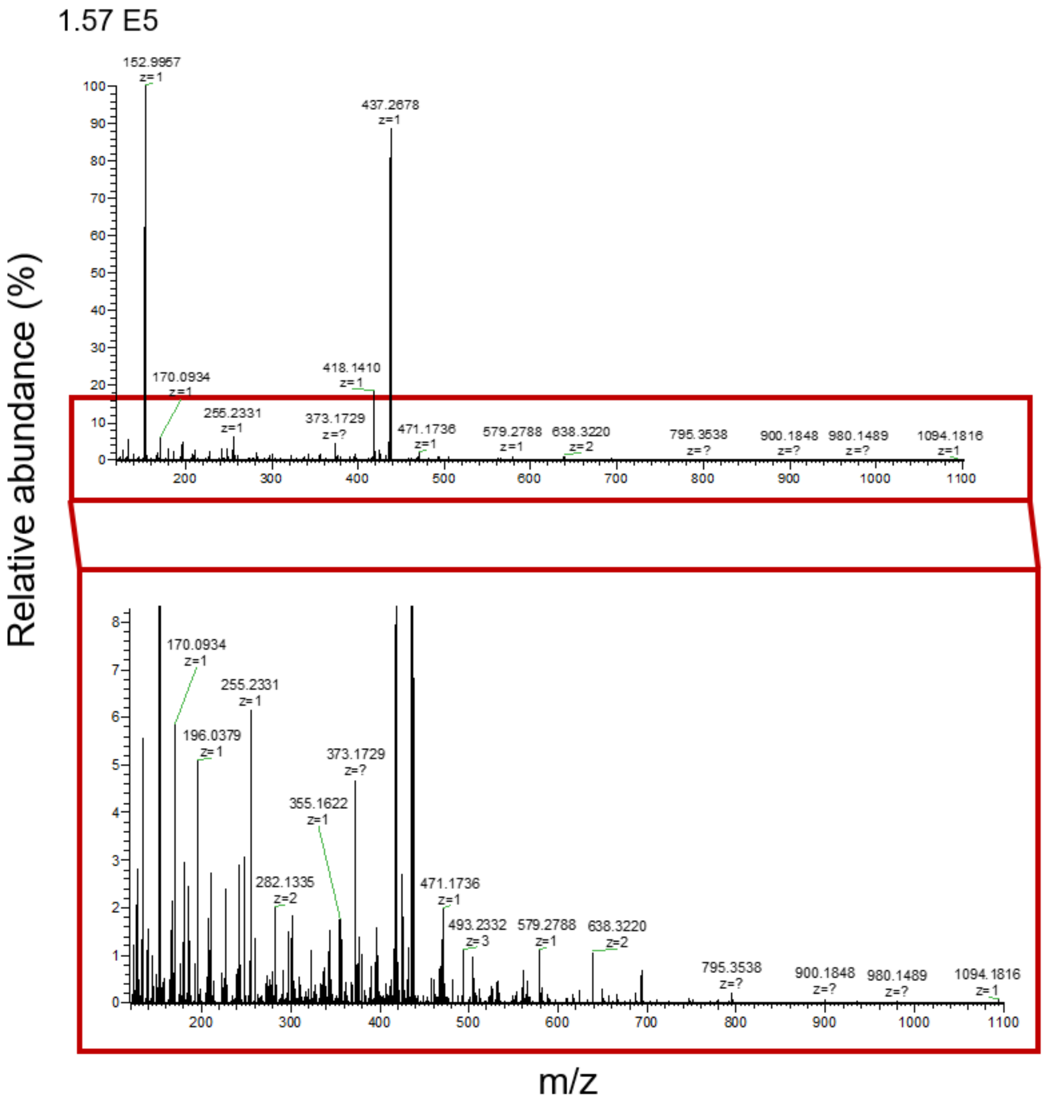


Fig. S37. MS/MS spectrum (top) and zoomed-in MS/MS spectrum (bottom) of the -4 charge state of the 1751 Da at m/z 436.7023.

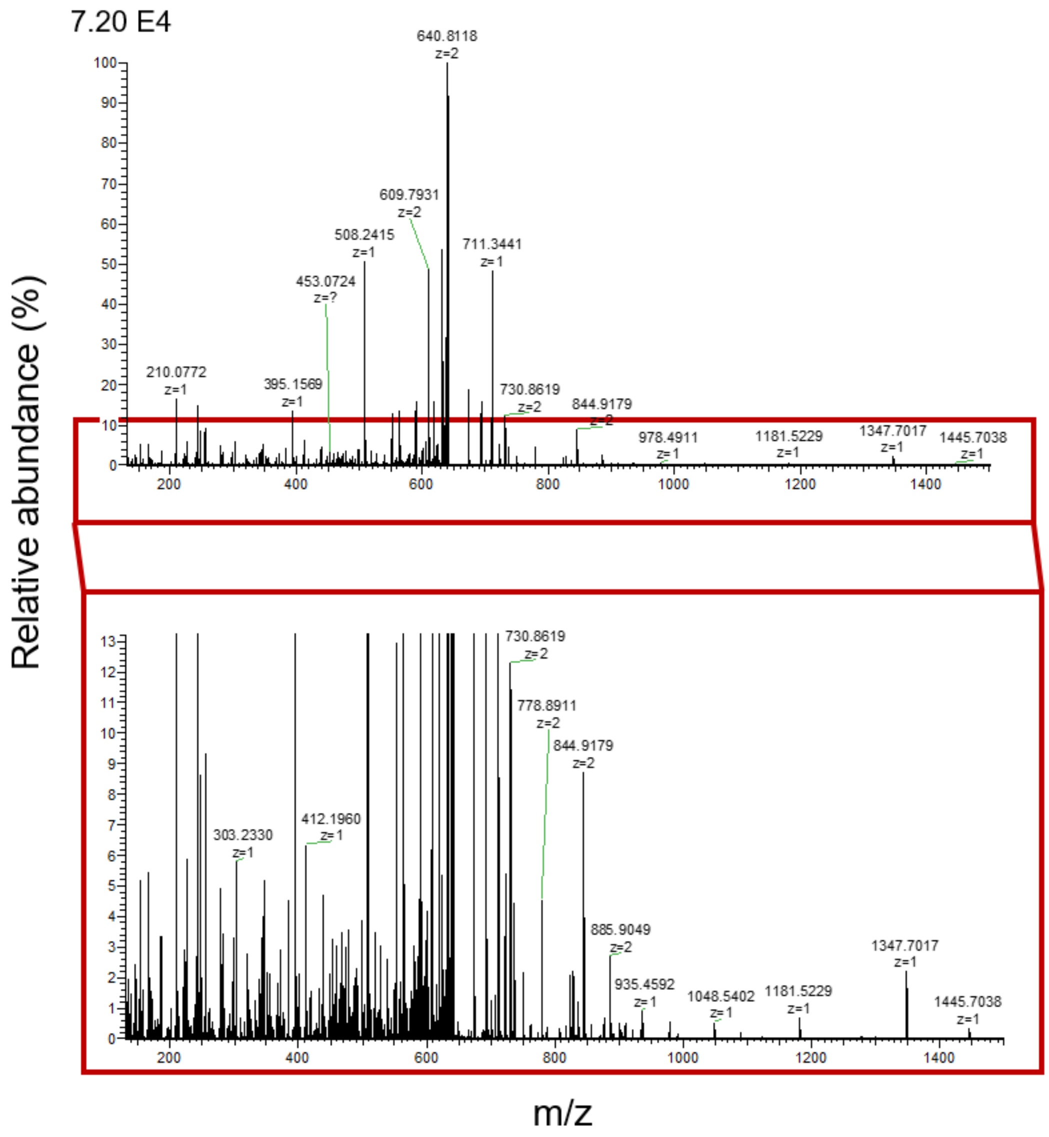


Fig. S38. MS/MS spectrum (top) and zoomed-in MS/MS spectrum (bottom) of the -3 charge state of the 1922 Da at m/z 639.6412.

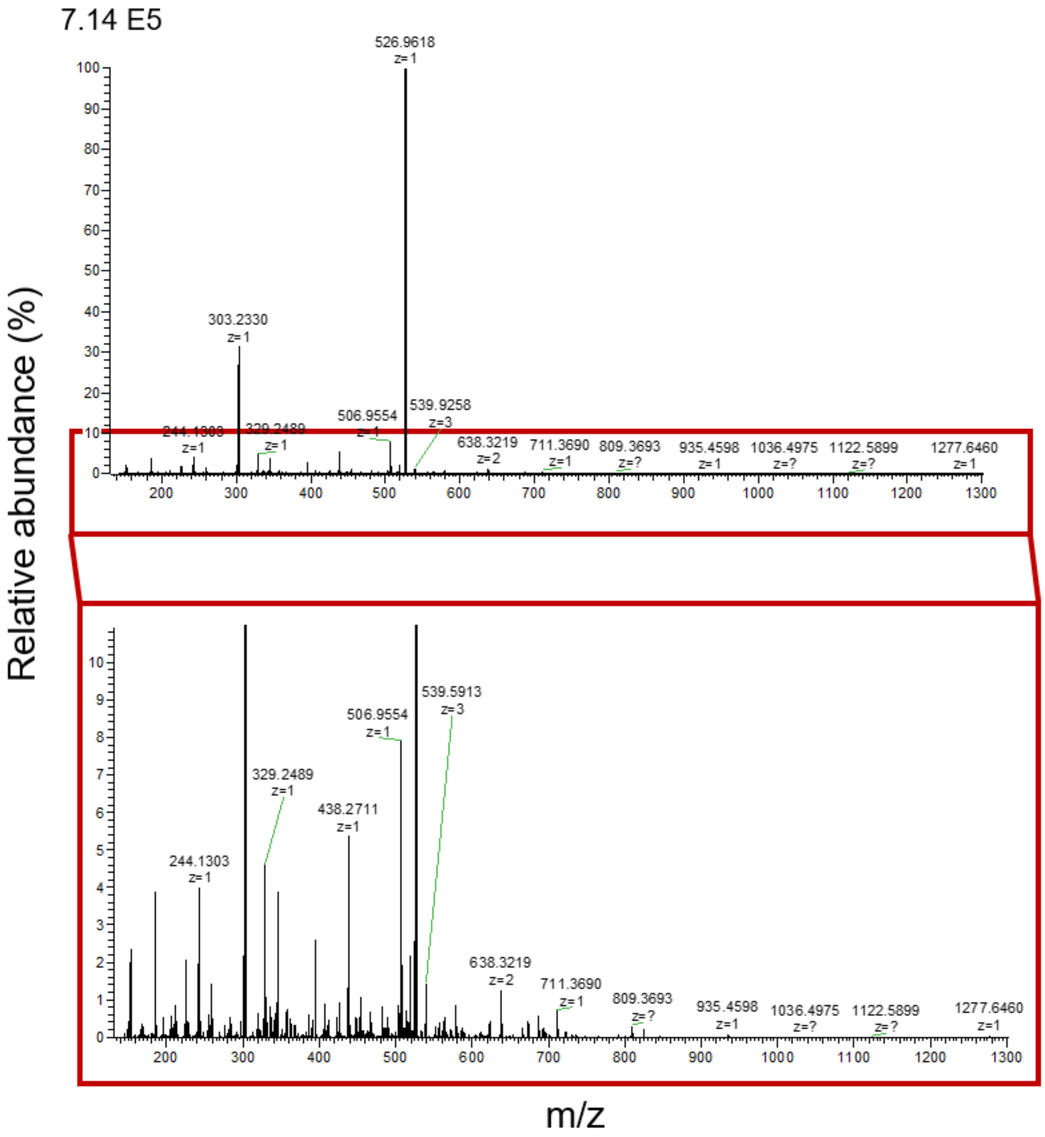


Fig. S39. MS/MS spectrum (top) and zoomed-in MS/MS spectrum (bottom) of the -4 charge state of the 2111 Da at m/z 526.7459.

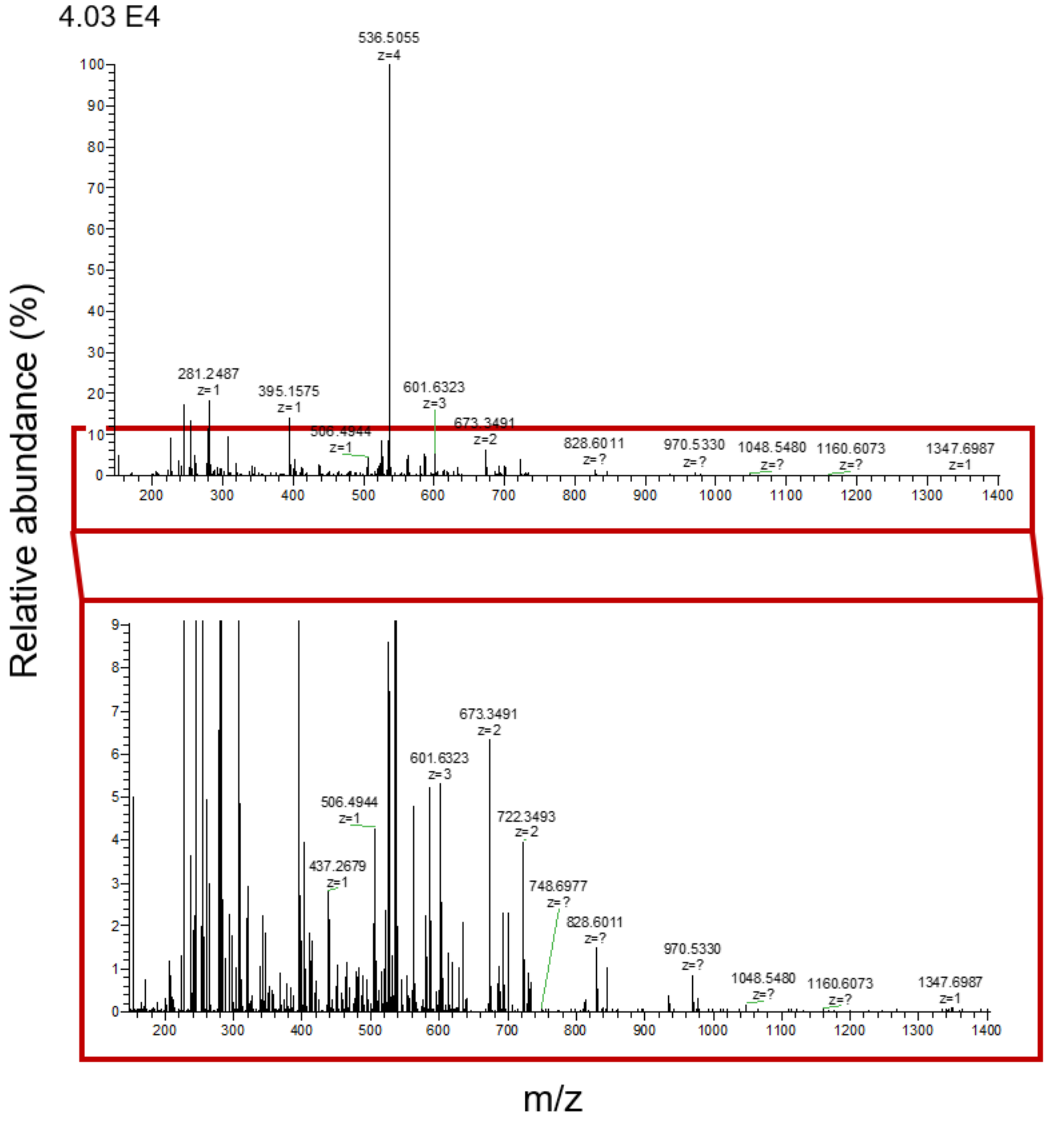


Fig. S40. MS/MS spectrum (top) and zoomed-in MS/MS spectrum (bottom) of the -4 charge state of the 2151 Da at m/z 536.7561.

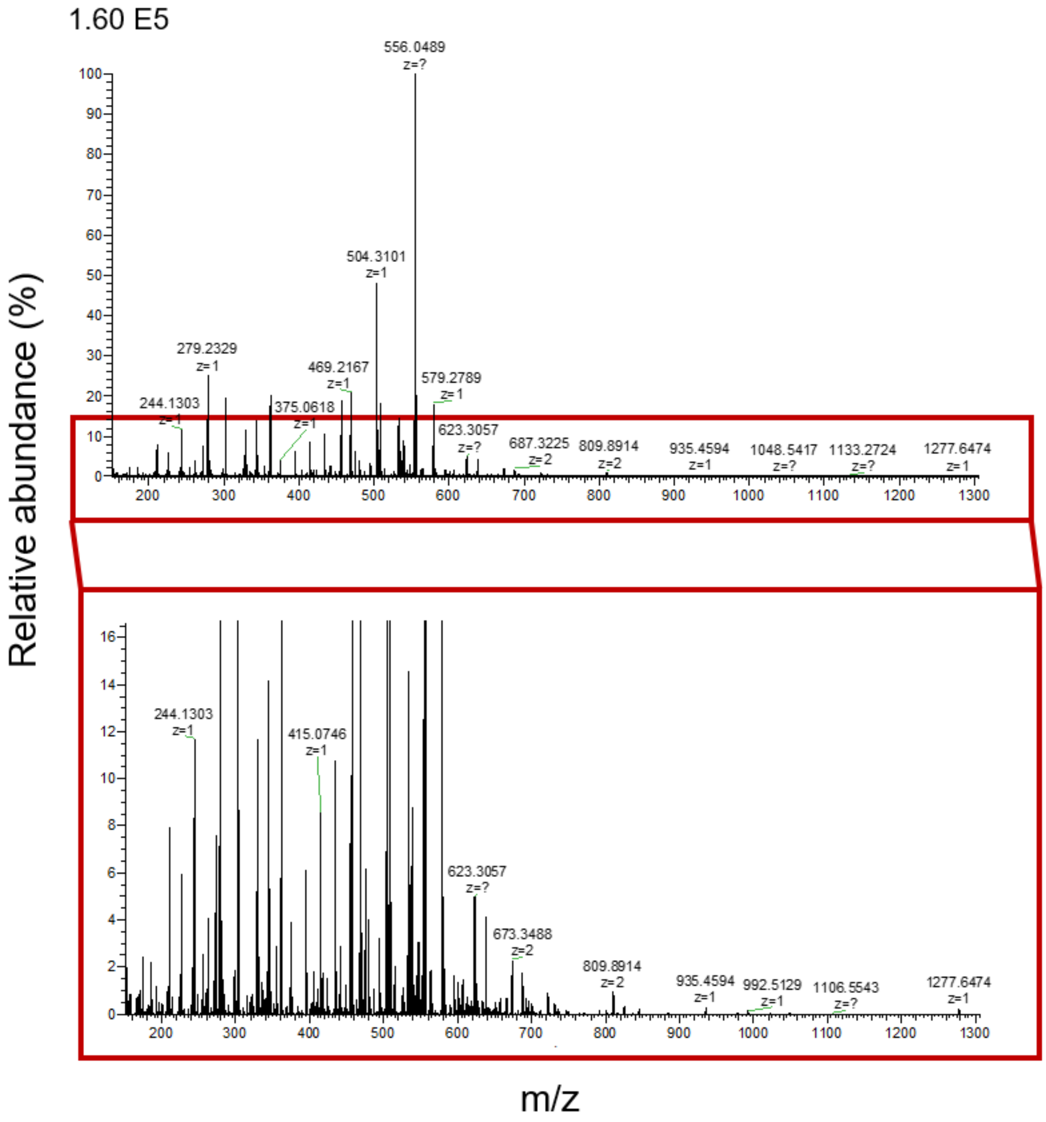


Fig. S41. MS/MS spectrum (top) and zoomed-in MS/MS spectrum (bottom) of the -4 charge state of the 2224 Da at m/z 555.0167.

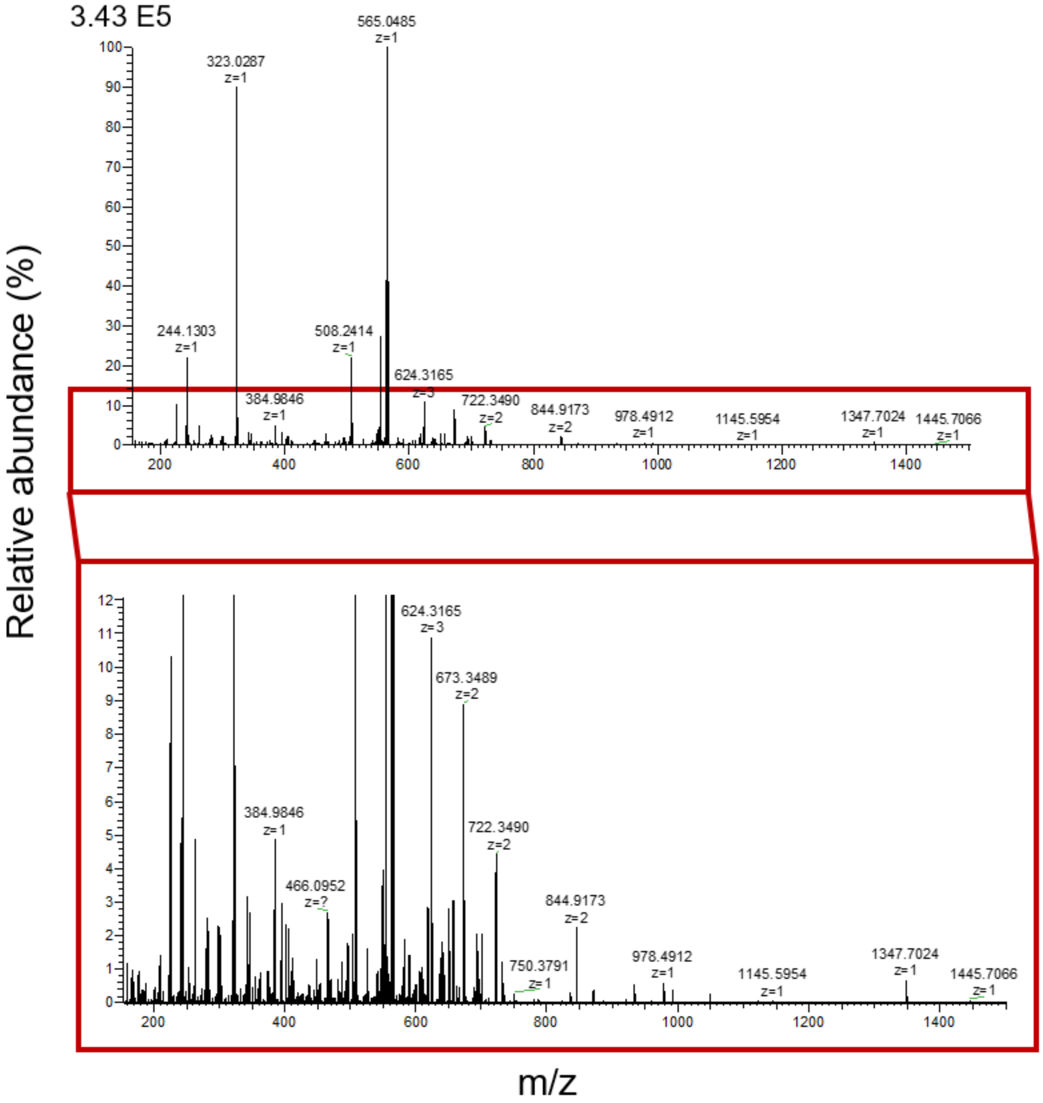


Fig. S42. MS/MS spectrum (top) and zoomed-in MS/MS spectrum (bottom) of the -4 charge state of the 2264 Da at m/z 565.0266.

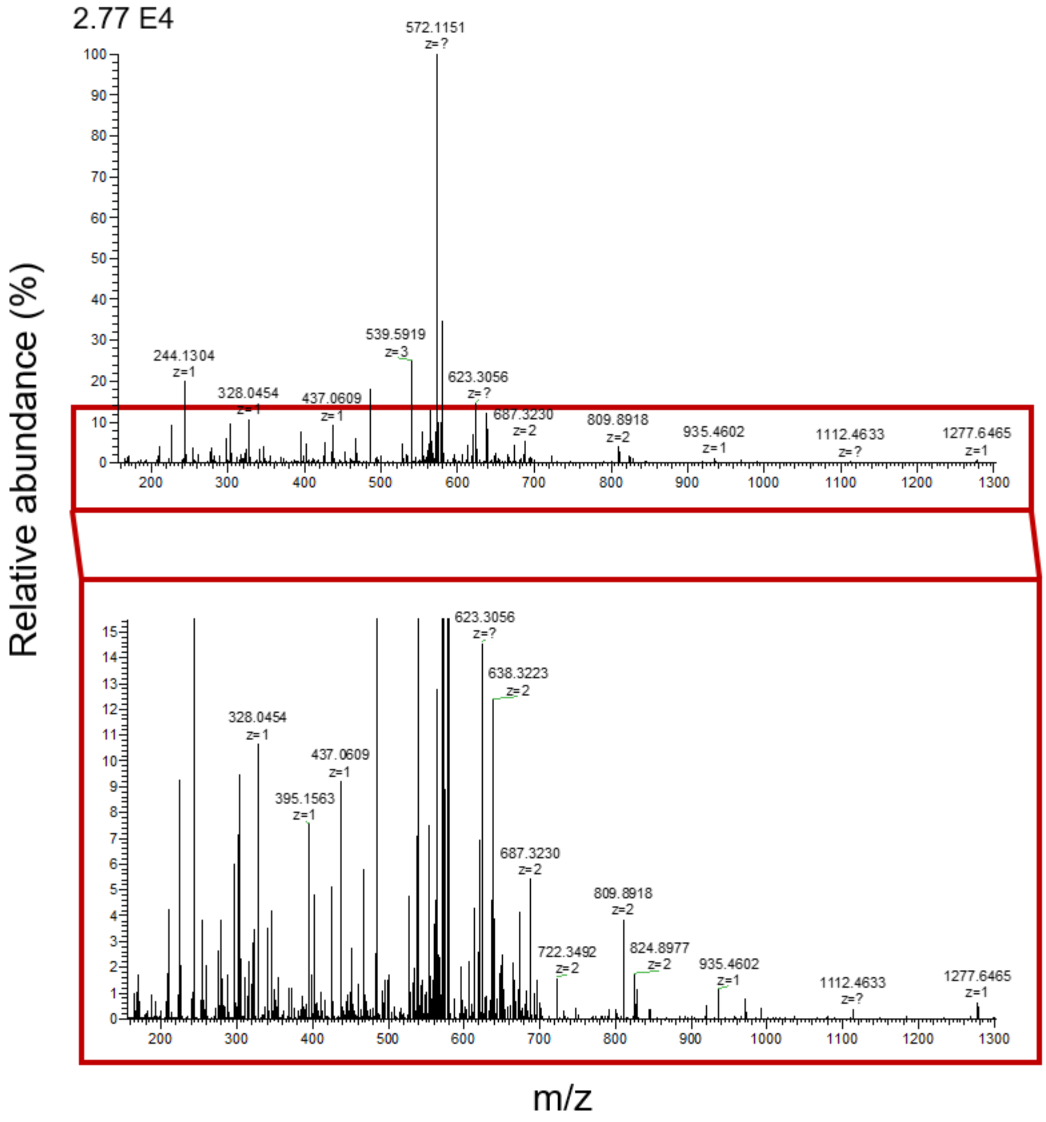


Fig. S43. MS/MS spectrum (top) and zoomed-in MS/MS spectrum (bottom) of the -4 charge state of the 2295 Da at m/z 572.7758.

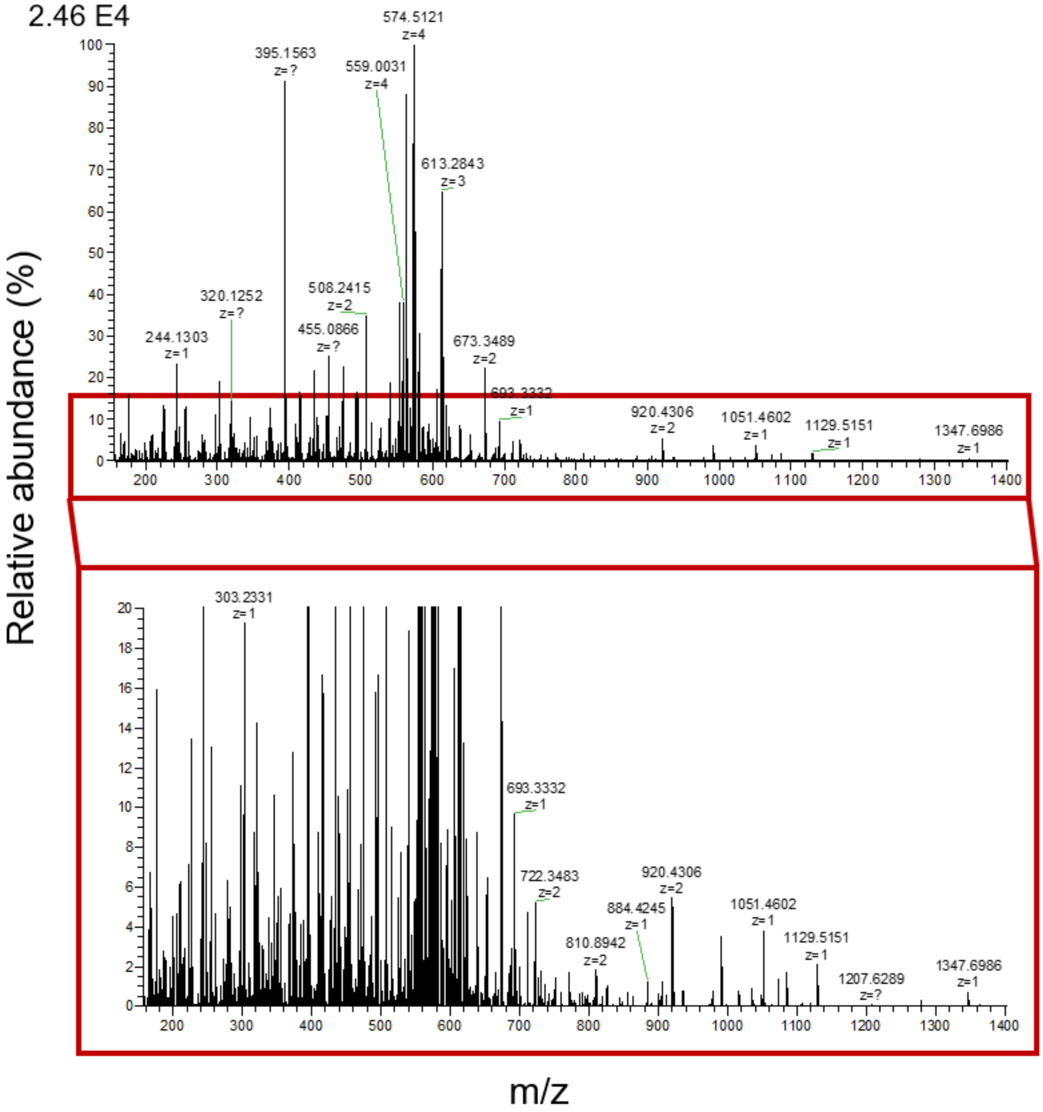


Fig. S44. MS/MS spectrum (top) and zoomed-in MS/MS spectrum (bottom) of the -4 charge state of the 2302 Da at m/z 574.5125.

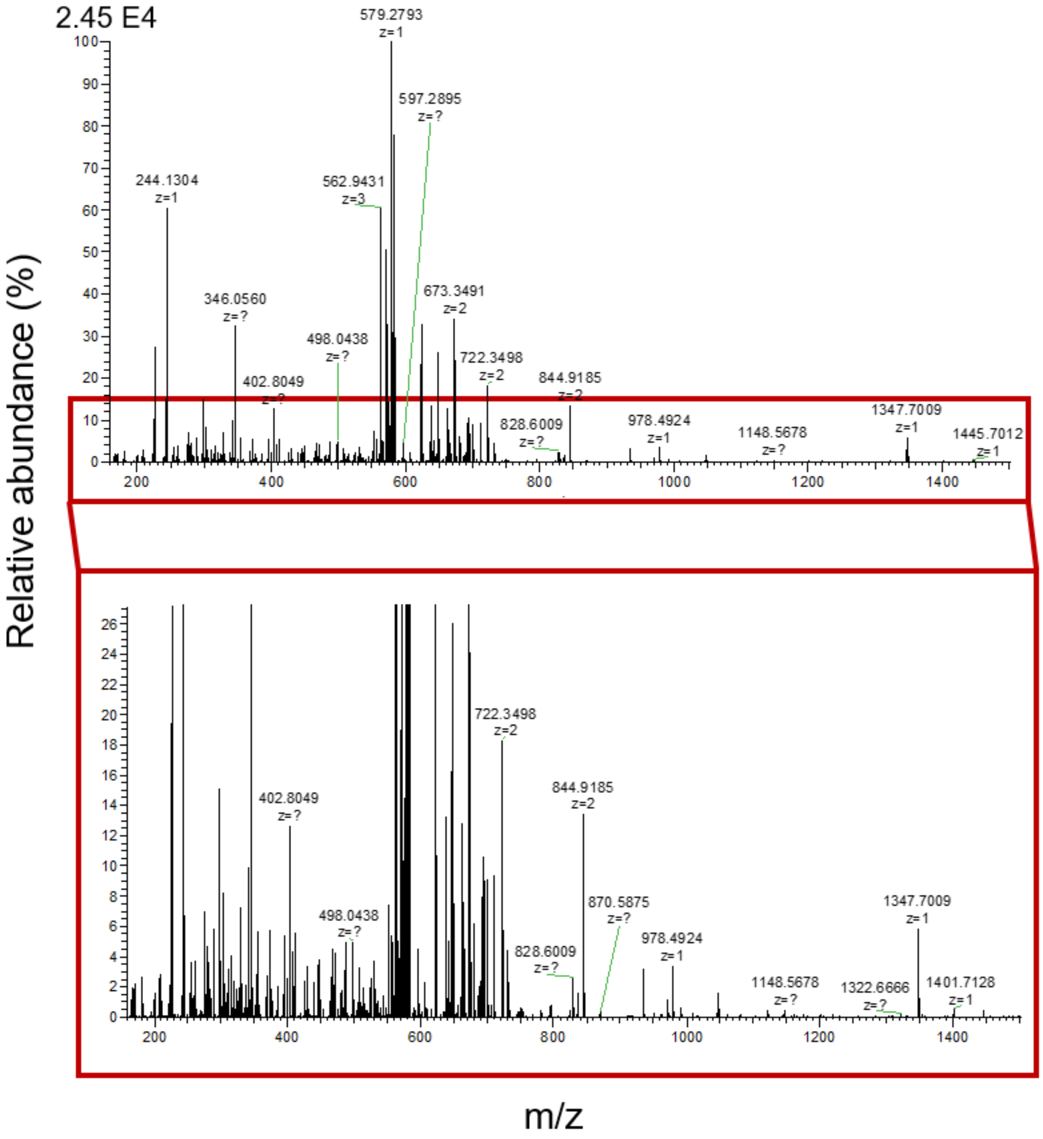


Fig. S45. MS/MS spectrum (top) and zoomed-in MS/MS spectrum (bottom) of the -4 charge state of the 2335 Da at m/z 582.7870.

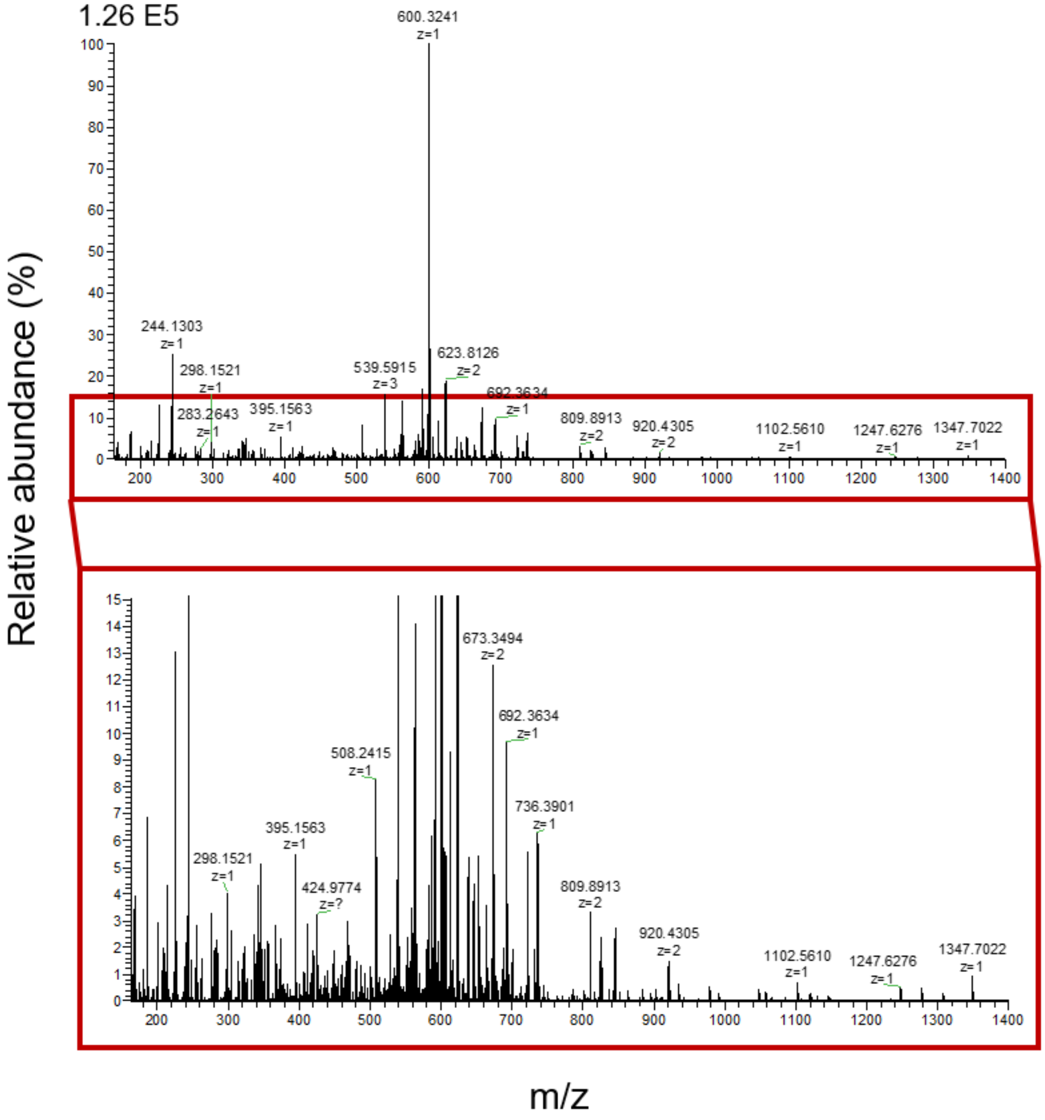


Fig. S46. MS/MS spectrum (top) and zoomed-in MS/MS spectrum (bottom) of the -4 charge state of the 2408 Da at m/z 601.0472.

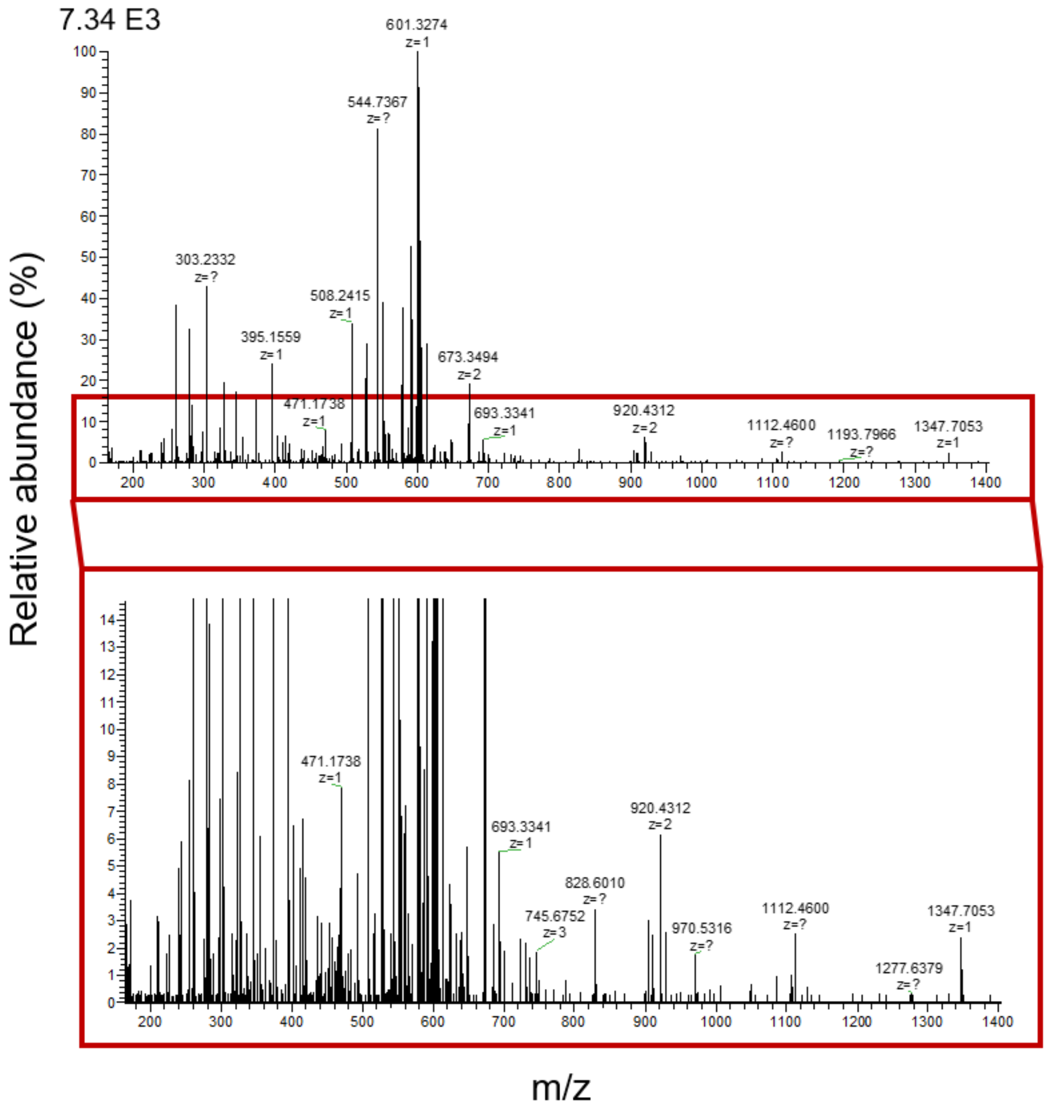


Fig. S47. MS/MS spectrum (top) and zoomed-in MS/MS spectrum (bottom) of the -4 charge state of the 2415 Da at m/z 602.7839.

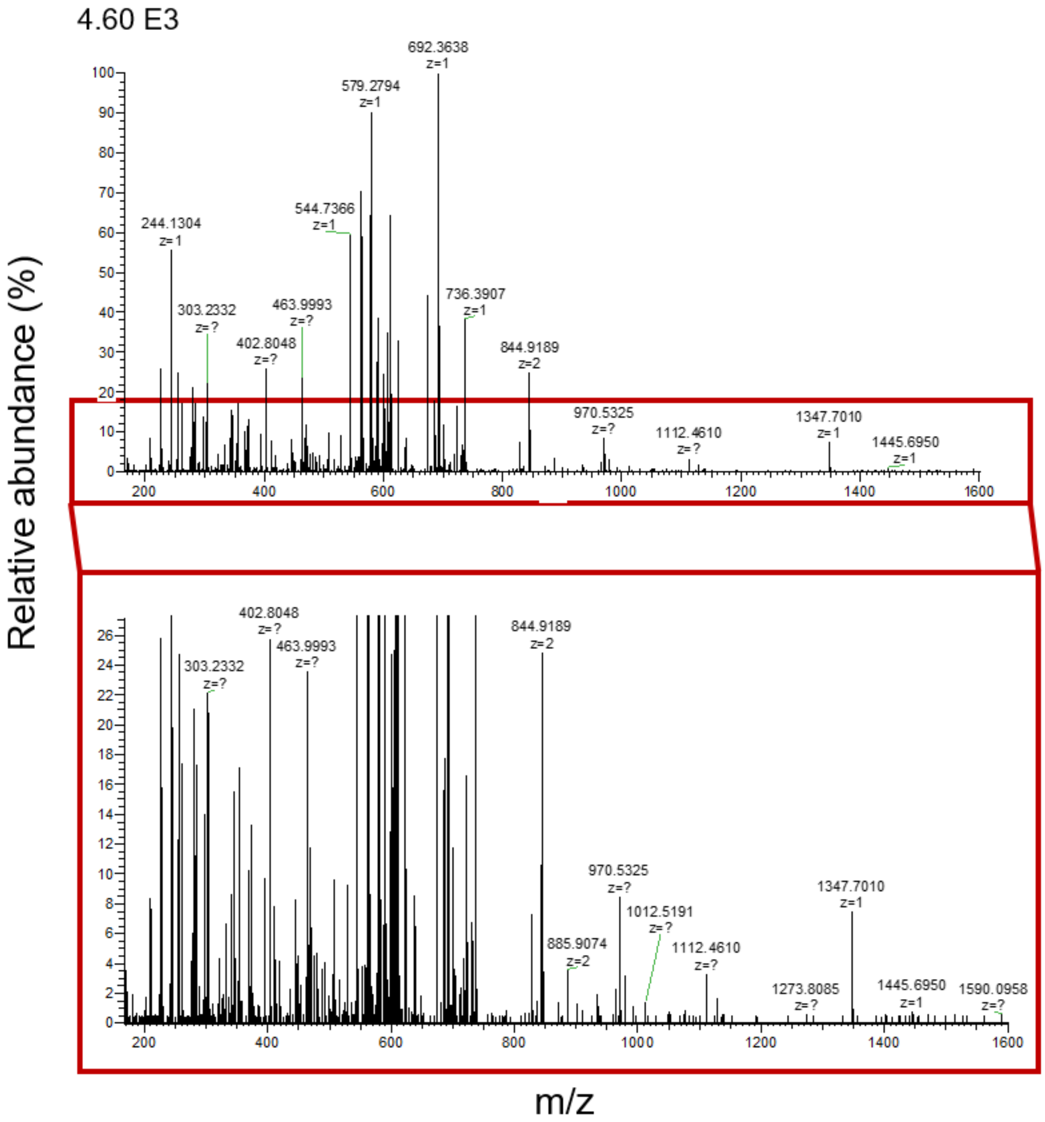


Fig. S48. MS/MS spectrum (top) and zoomed-in MS/MS spectrum (bottom) of the -4 charge state of the 2448 Da at m/z 611.0578.

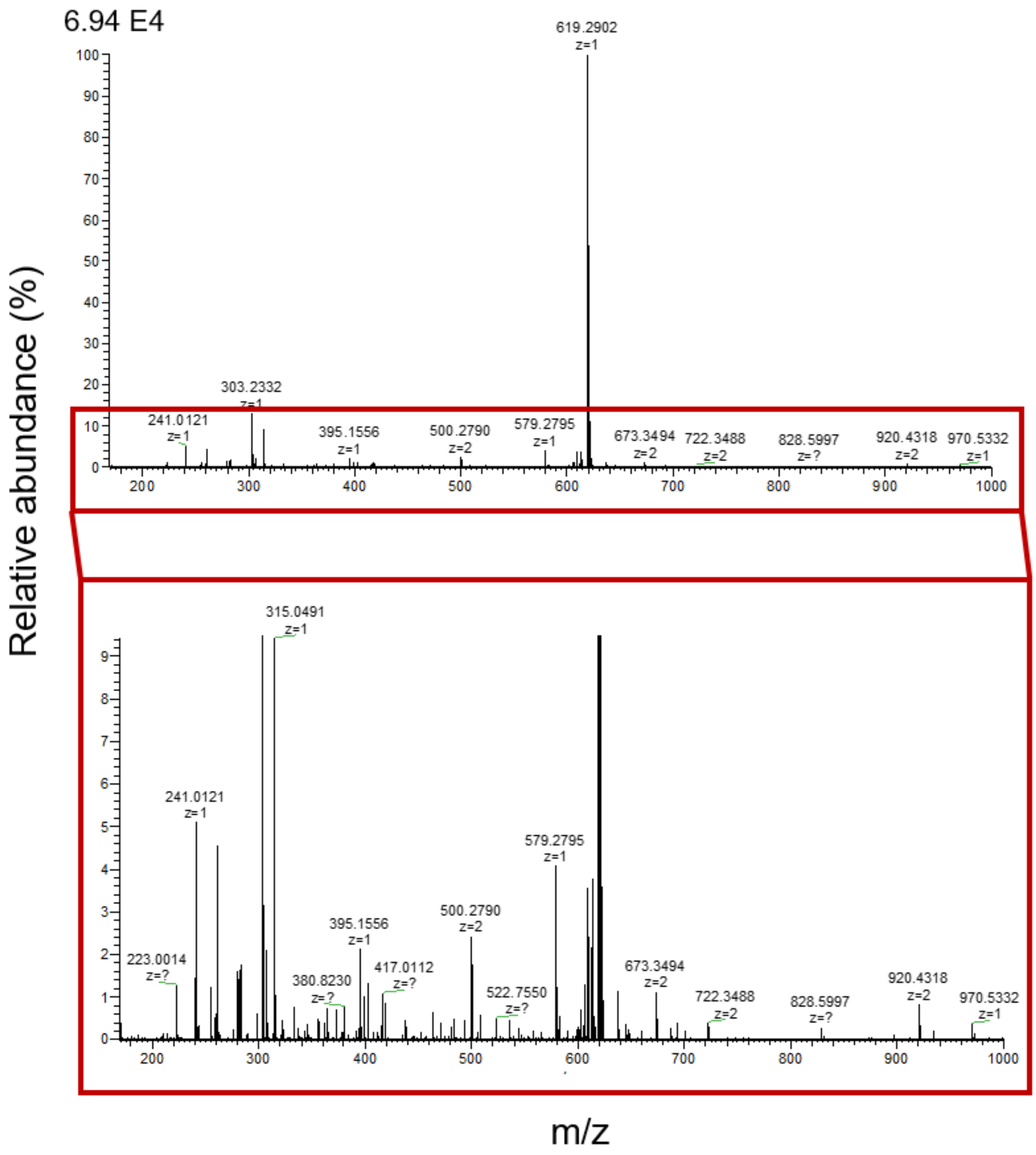


Fig. S49. MS/MS spectrum (top) and zoomed-in MS/MS spectrum (bottom) of the -4 charge state of the 2486 Da at m/z 620.5431.

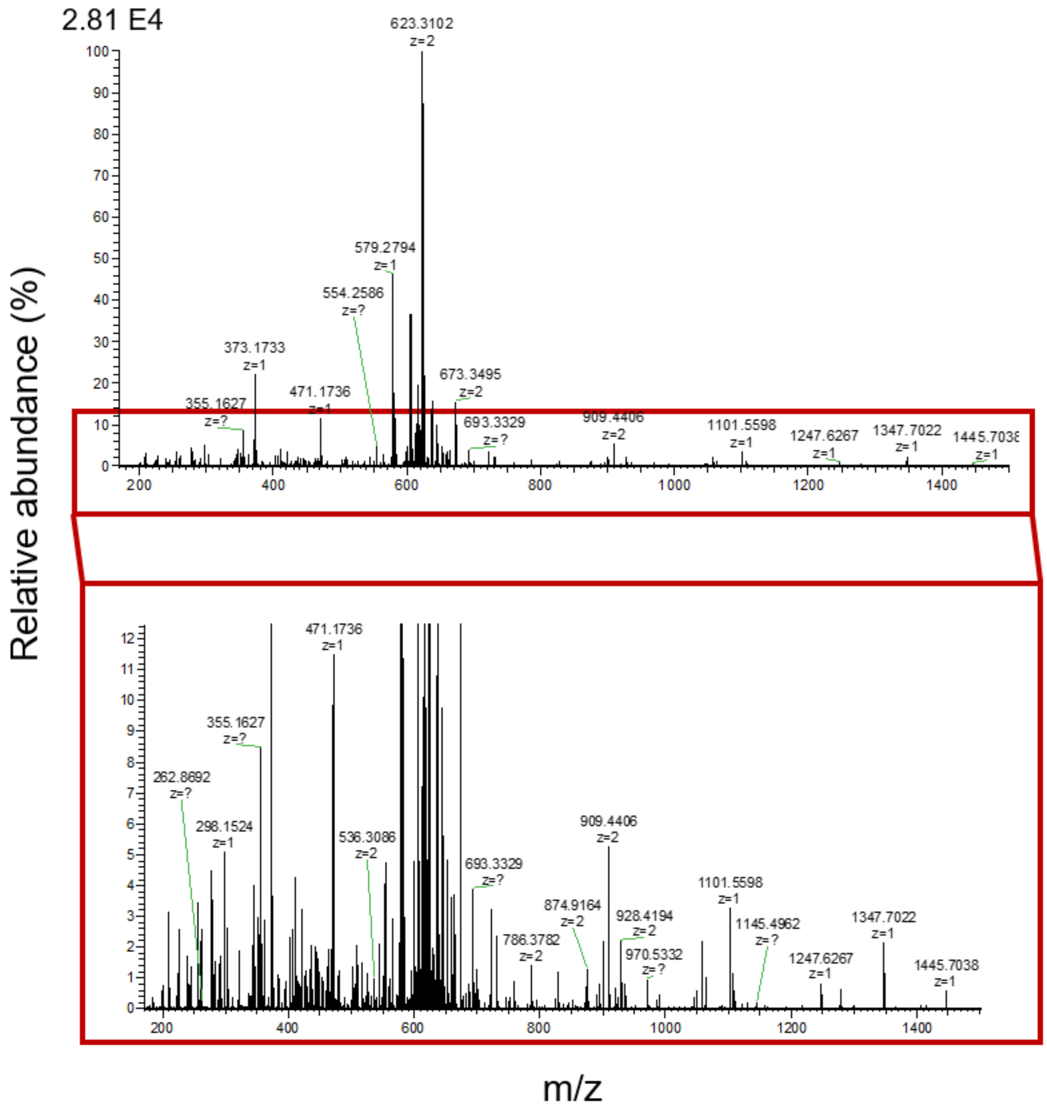


Fig. S50. MS/MS spectrum (top) and zoomed-in MS/MS spectrum (bottom) of the -4 charge state of the 2502 Da at m/z 624.5357.

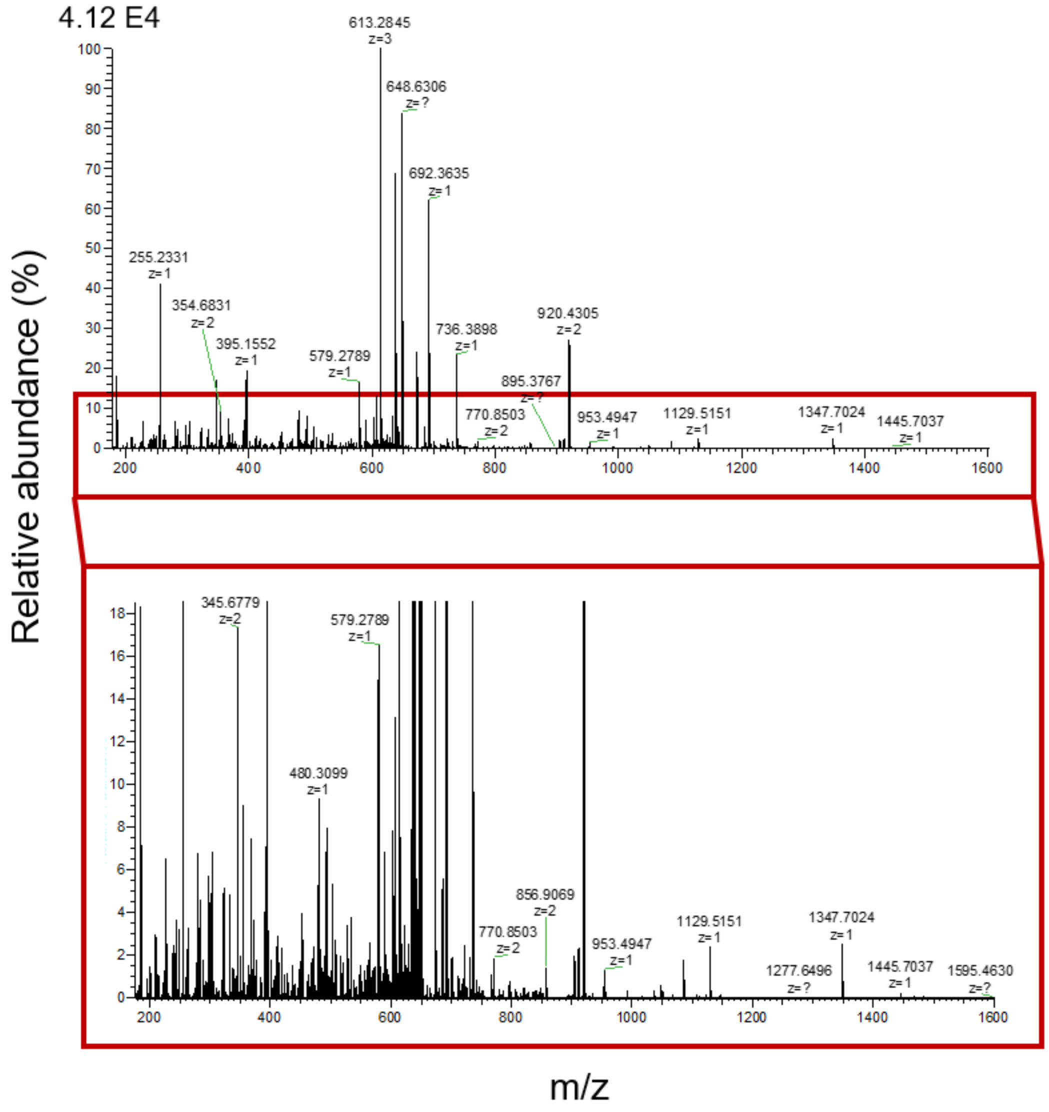


Fig. S51. MS/MS spectrum (top) and zoomed-in MS/MS spectrum (bottom) of the -4 charge state of the 2599 Da at m/z 648.8137.

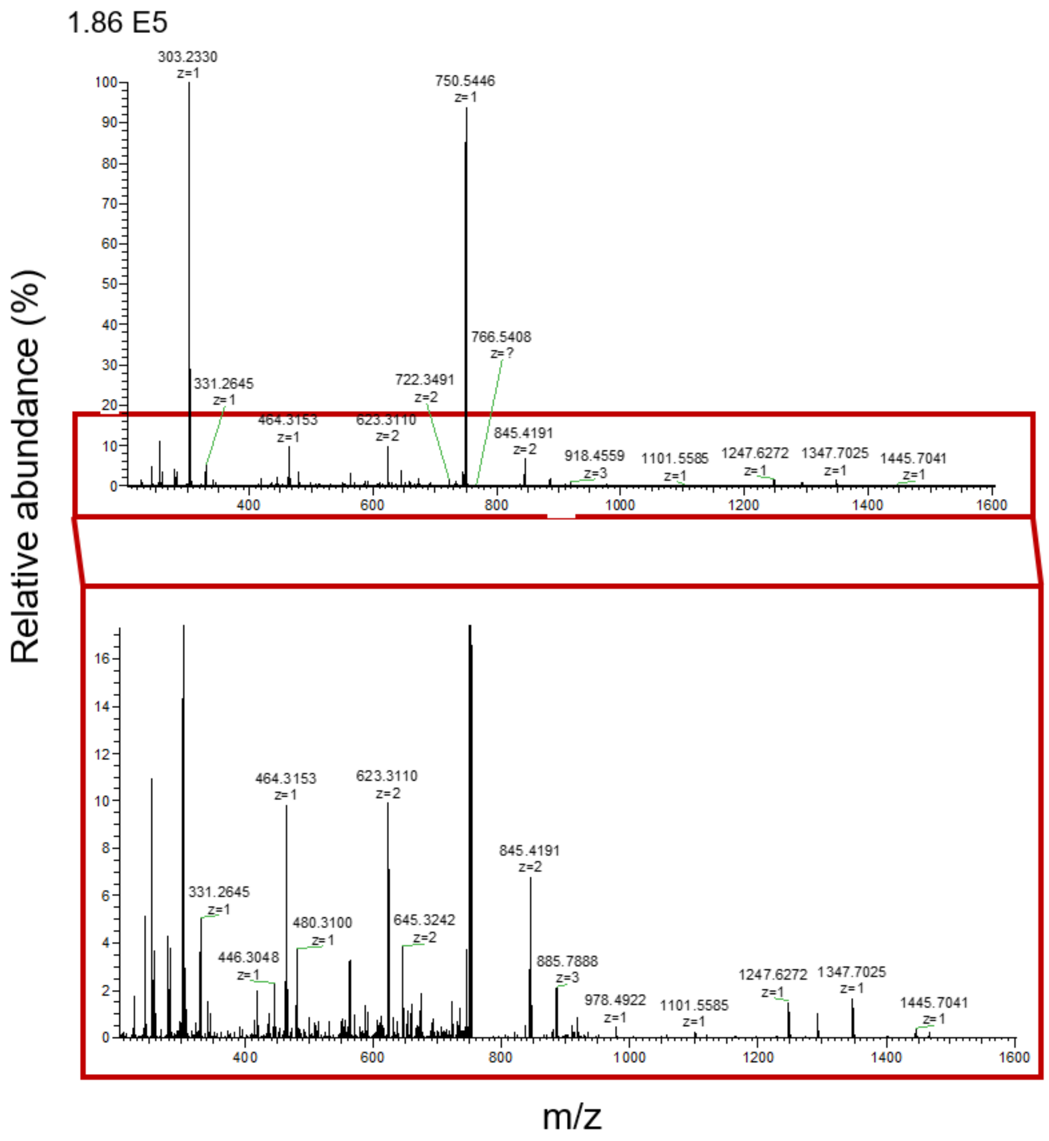


Fig. S52. MS/MS spectrum (top) and zoomed-in MS/MS spectrum (bottom) of the -4 charge state of the 3007 Da at m/z 750.8808.

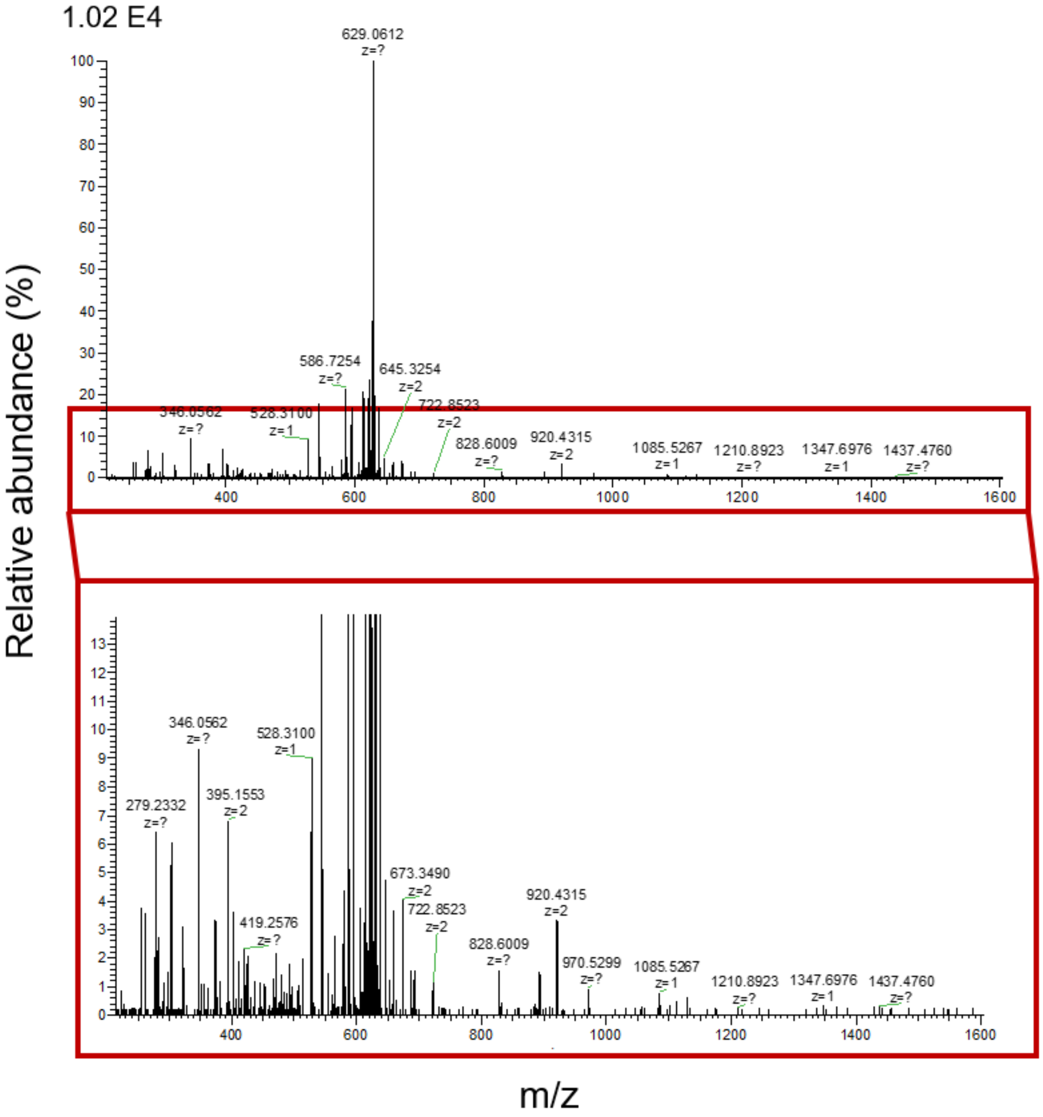


Fig. S53. MS/MS spectrum (top) and zoomed-in MS/MS spectrum (bottom) of the -5 charge state of the 3154 Da at m/z 629.9023.

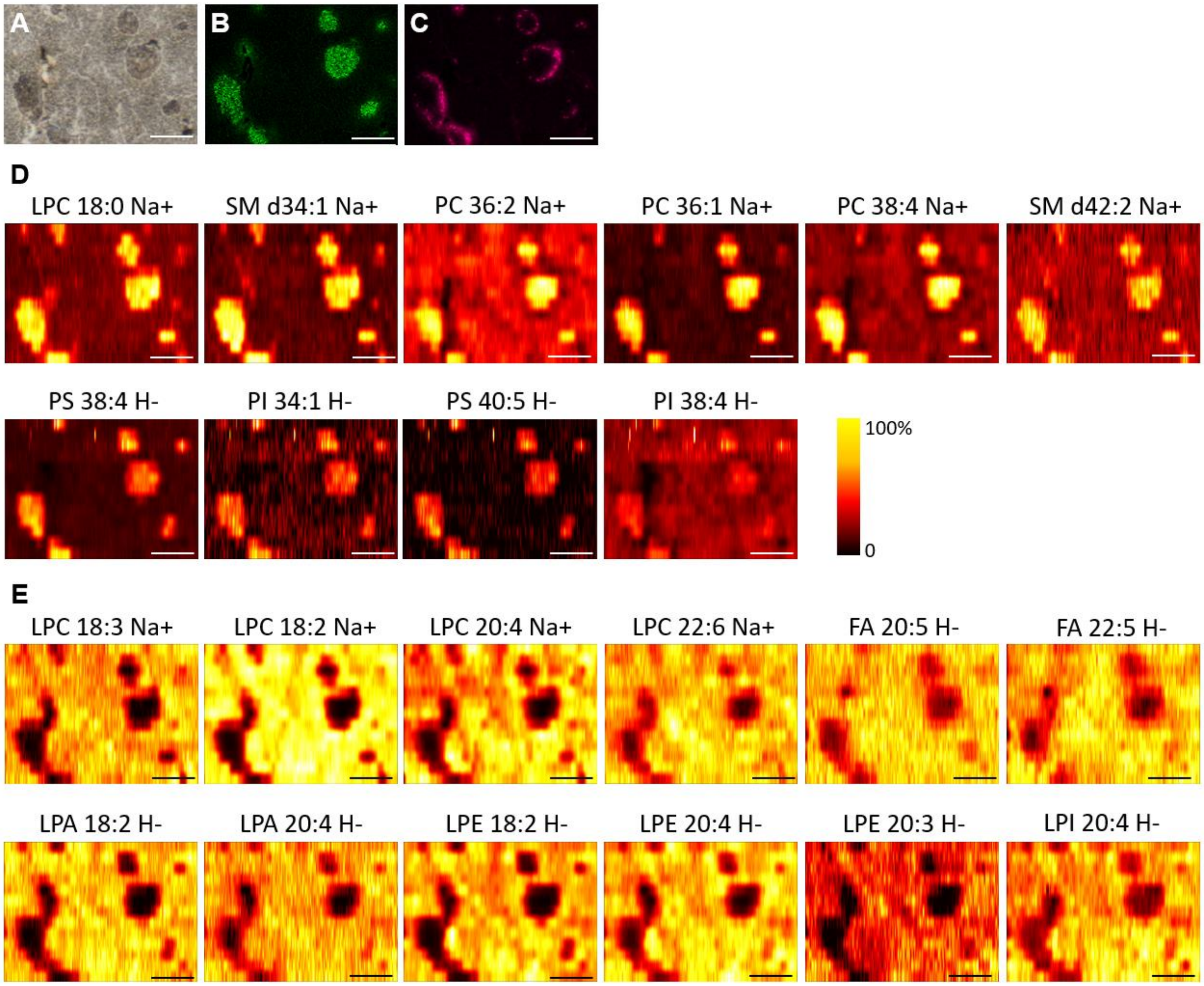


Fig. S54. Ion images of lipids and metabolites obtained from replicate 1.

(A) Brightfield optical image of the analyzed region of the pancreatic tissue section. IF image of (B) INS and (C) glucagon on the adjacent section. Ion images of lipids and metabolites normalized to TIC with (D) species showing enhanced abundance in β-cells and (E) species showing diminished abundance in islets of Langerhans. Scale bar: 250 µm.

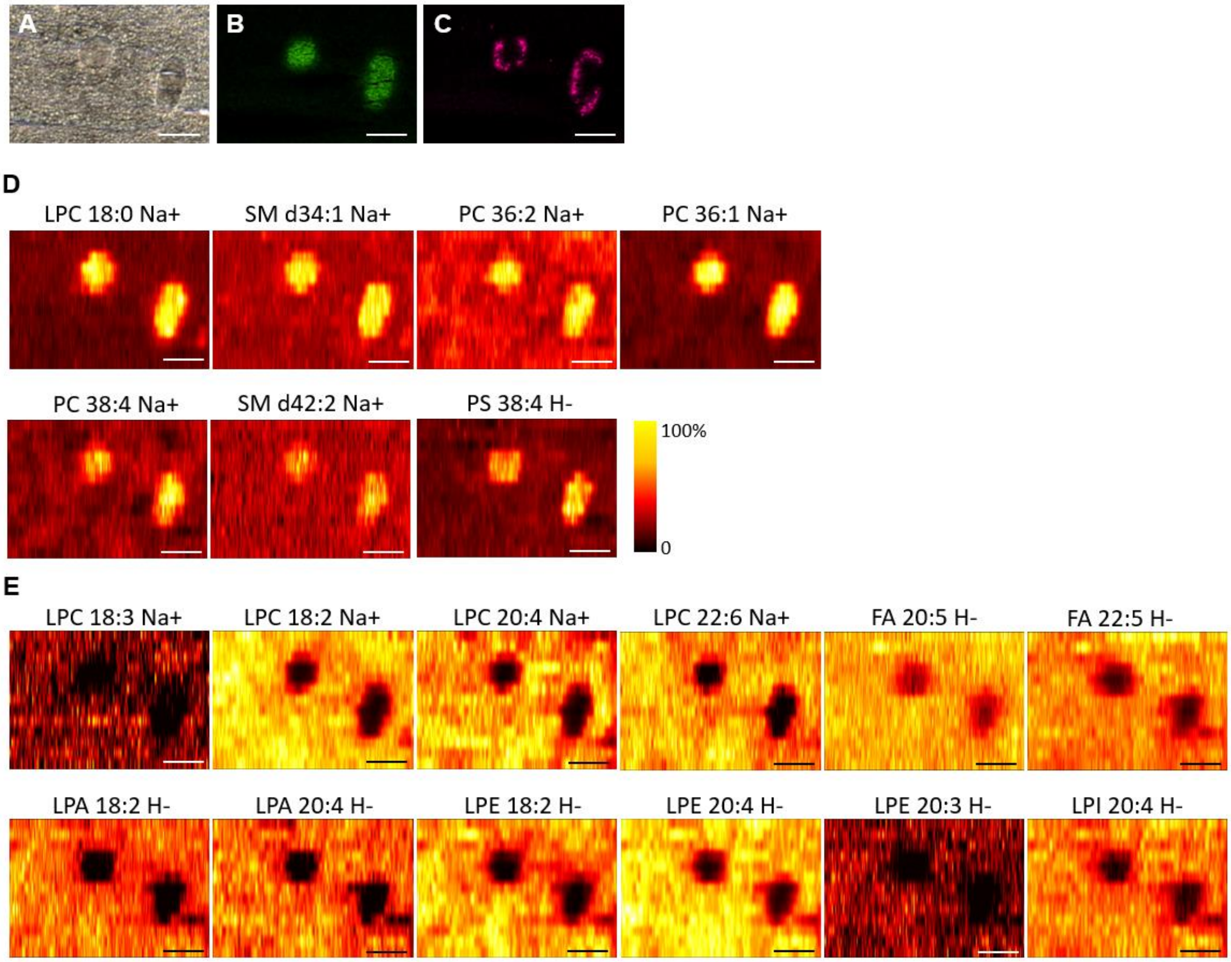


Fig. S55. Ion images of lipids and metabolites obtained from replicate 2.

(A) Brightfield optical image of the analyzed region of the pancreatic tissue section. IF image of (B) INS and (C) glucagon on the adjacent section. Ion images of lipids and metabolites normalized to TIC with (D) species showing enhanced abundance in β-cells and (E) species showing diminished abundance in islets of Langerhans. Scale bar: 250 µm.

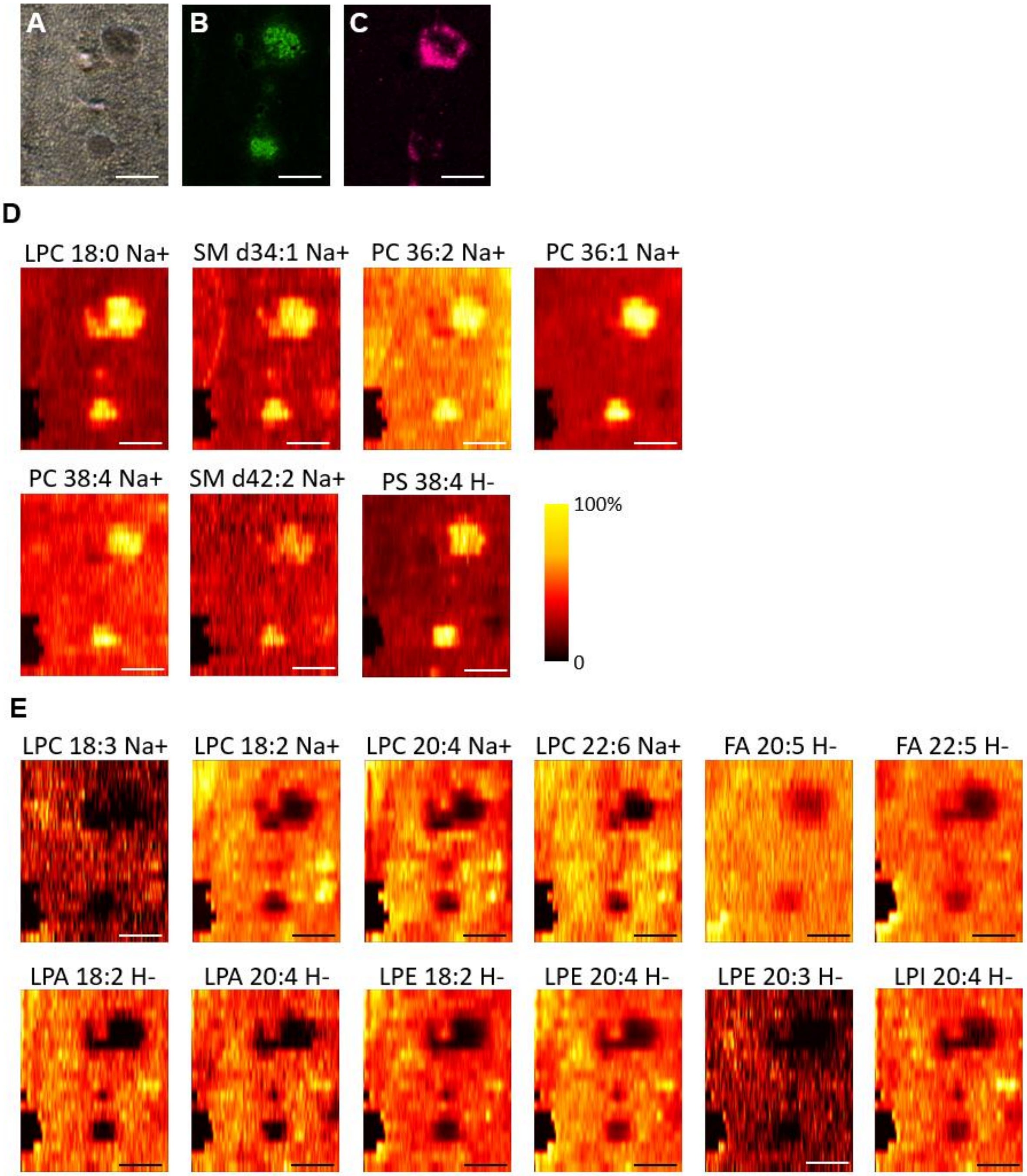


Fig. S56. Ion images of lipids and metabolites obtained from replicate 3.

(A) Brightfield optical image of the analyzed region of the pancreatic tissue section. IF image of (B) INS and (C) glucagon on the adjacent section. Ion images of lipids and metabolites normalized to TIC with (D) species showing enhanced abundance in β-cells and (E) species showing diminished abundance in islets of Langerhans. Scale bar: 250 µm.

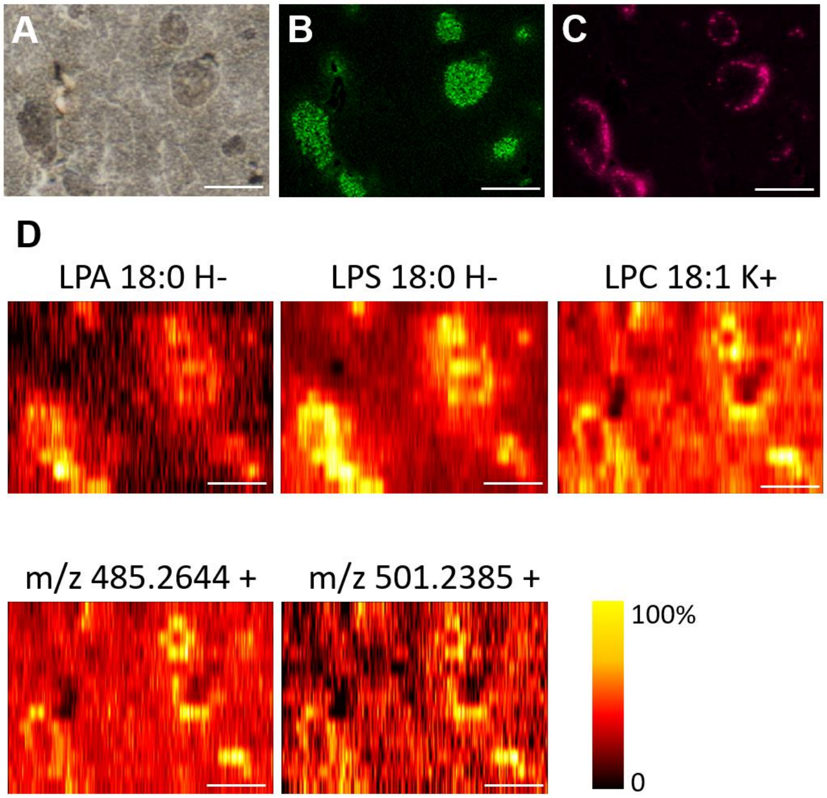


Fig. S57. Ion images of species showing signal enhancement in cell clusters at the periphery of each islet.

(A) Brightfield optical image of the analyzed region of the pancreatic tissue section. IF image of (B) INS and (C) glucagon on the adjacent section. (D) TIC-normalized ion images of species showing enhanced abundance in cell clusters at the periphery of each islet. Scale bar: 250 µm.

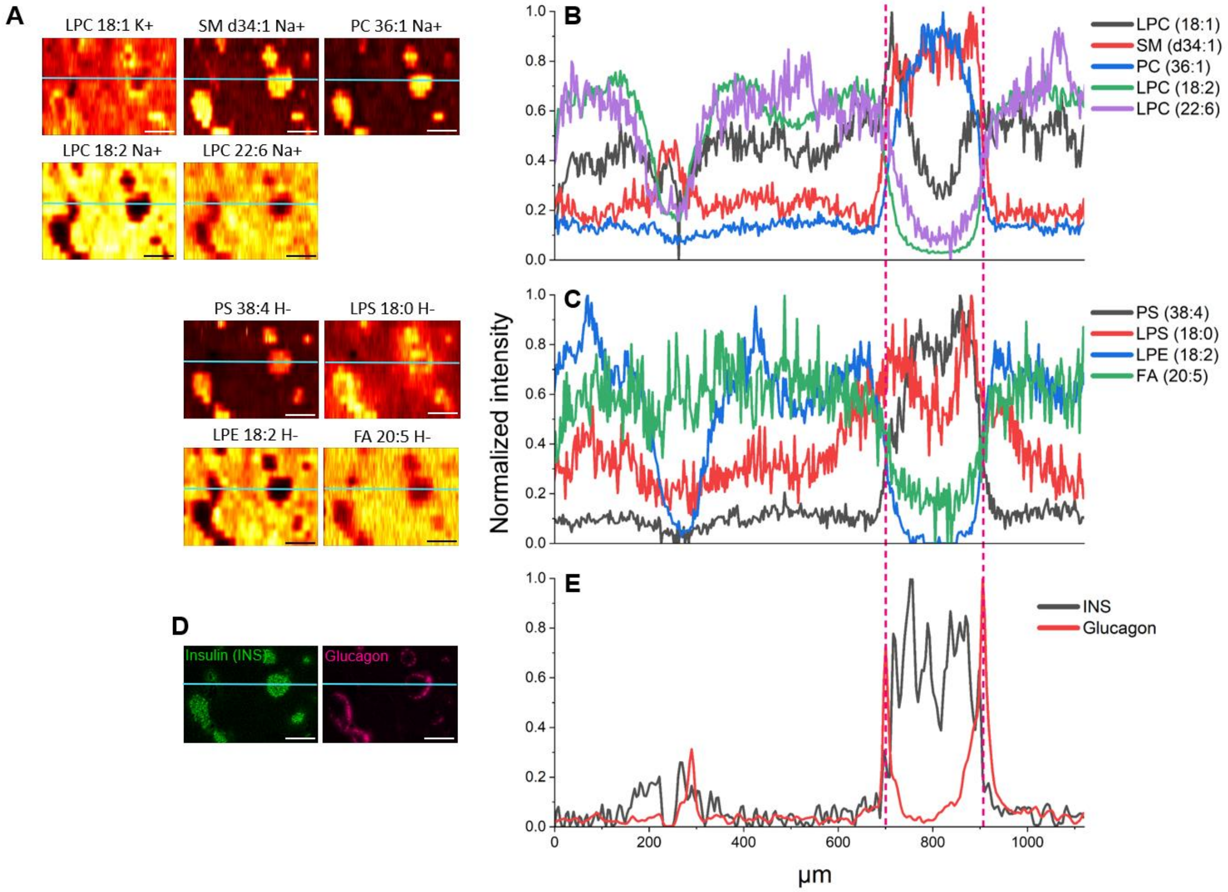


Fig. S58. Extracted line profiles of lipids and metabolites from ion images and extracted line profiles of insulin and glucagon from IF images.

(A) The ion images of lipids with the light blue trace marking the location of the extracted line profiles. Overlay of the line profiles of lipids detected in (B) positive ionization mode and (C) negative ionization mode. (D) The insulin and glucagon IF images with the light blue trace marking the location of (E) the extracted line profiles. Scale bar: 250 µm. The signal intensities were self-normalized.
